# Selection-driven adaptation to the extreme Antarctic environment in the Emperor penguin

**DOI:** 10.1101/2021.12.14.471946

**Authors:** Federica Pirri, Lino Ometto, Silvia Fuselli, Flávia A. N. Fernandes, Lorena Ancona, Céline Le Bohec, Lorenzo Zane, Emiliano Trucchi

## Abstract

The eco-evolutionary history of penguins is profoundly influenced by their shift from temperate to cold environments. Breeding only in Antarctica during the winter, the Emperor penguin appears as an extreme outcome of this process, with unique features related to insulation, heat production and energy management. However, whether this species actually diverged from a less cold-adapted ancestor, thus more similar in ecology to its sister species, the King penguin, is still an open question. As the Antarctic niche shift likely resulted in vast changes in selective pressure experienced by the Emperor penguin, the identification and relative quantification of the genomic signatures of selection, unique to each of these sister species, could answer this question. Applying a suite of phylogeny-based methods on 7,651 orthologous gene alignments of seven penguins and 13 other birds, we identified a set of candidate genes showing significantly different selection regimes either in the Emperor or in the King penguin lineage. Our comparative approach unveils a more pervasive selection shift in the Emperor penguin, supporting the hypothesis that its extreme cold adaptation is a derived state from a more King penguin-like ecology. Among the candidate genes under selection in the Emperor penguin, four genes (TRPM8, LEPR, CRB1, and SFI1) were identified before in other cold adapted vertebrates, while, on the other hand, 161 genes can be assigned to functional pathways relevant to cold adaptation (e.g., cardiovascular system, lipid, fatty acid and glucose metabolism, insulation, etc.). Our results show that extreme cold adaptation in the Emperor penguin largely involved unique genetic options which, however, affect metabolic and physiological traits common to other cold-adapted homeotherms.

## Introduction

Adaptation to severe Arctic and Antarctic temperatures is rare among terrestrial vertebrates and restricted to warm-blooded lineages (Storey and Storey, 1992; Blix, 2016). The main challenge for adaptation to extreme cold is keeping adequately high core body temperature, which in homeotherms can be obtained by the combination of several physiological, morphological, and behavioral responses (Allen, 1877; Scholander, 1955; Cannon and Nedergaard, 2010; Tattersall et al., 2012; Blix, 2016; Roussel et al., 2020): *i)* Minimizing heat dissipation from the body surface *e.g*., by reducing the surface area-to-volume ratio, by raising insulating hair or feathers, by decreasing peripheral circulation, and/or by balling up or huddling; *ii)* Increasing heat production via shivering and nonshivering thermogenesis through brown adipocytes; *iii)* Temporarily hibernating.

In contrast, the genetic basis of cold adaptation in homeotherms is not well understood. Only few candidate genes were found in common across cold-adapted mammals and birds (Yudin et al., 2017; Wollenberg Valero et al., 2014), suggesting that cold adaptation is likely the result of selection on different genes, which are nevertheless relevant to the same set of physiological and metabolic functions. For example, only four candidate genes, which are related to cardiovascular function, were found in common in three out of six mammals dwelling in/near the Arctic or Antarctica (Yudin et al., 2017), even if different candidate genes are involved anyway in heart and vascular development and regulation (Liu et al., 2014; Vianna et al., 2020). Different genes acting in fatty-acid metabolism have been identified to be under positive selection in the polar bear and the Arctic fox (Liu et al., 2014; Castruita et al., 2020; Kumar et al., 2015). In mammals, the molecular mechanisms underpinning heat production via non-shivering thermogenesis have been extensively investigated and have been mainly linked to UCP-1, a mitochondrial uncoupling protein involved in the generation of heat in brown adipose tissue (Lowell and Spiegelman, 2000). Non-shivering thermogenesis is instead not well understood in birds, where it is not clear whether the avian homologue of UCP-1 (i.e., avian uncoupling protein, avUCP) has a similar thermogenic role or is mainly required against oxidative stress (Talbot et al., 2004). However, the same non-shivering thermogenic pathways could be used in birds (Tigano et al., 2018), and even in ectothermic vertebrates like reptiles (Akashi et al., 2016).

Birds are known for their fast adaptation rate, which allowed the colonization of a huge diversity of environments, including polar ones (Zhang et al., 2014). In particular, the clade of penguins, which likely originated in temperate environments, successfully diversified in the cold Antarctic and sub-Antarctic ecosystems (Pan et al., 2019; Vianna et al., 2020), featuring unique adaptations for insulation, heat production and energy management (Scholander, 1955; Rowland et al., 2015). Our understanding of the underlying genetic determinants of such adaptations is still rather scarce. Testing about one third of the total genes across all penguin genomes, Vianna et al. (2020) identified blood pressure, cardiovascular regulation, and oxygen metabolism in muscles as functions, potentially involved in thermoregulation, which have been the targets of positive selection. Analyses of *Spheniscus* and *Pygoscelis* mitochondrial genomes also revealed a correlation between the pattern of diversity of the ND4 gene and sea surface temperature, suggesting this gene is involved in climate adaptation (Ramos et al., 2018). Moreover, Emperor and Adelie penguin genomes show the highest rate of duplication of *β*-keratin genes as compared to non-penguin birds, suggesting important changes in feathers and skin structure during their evolution to increase core body insulation (Li et al., 2014). Alternative gene pathways related to lipid metabolism and phototransduction appeared also to have been under selection in these two species (Li et al., 2014).

The Emperor penguin (*Aptenodytes forsteri*) is the only warm-blooded vertebrate thriving and breeding during the harshest Antarctic winter, facing profound seasonal changes in daylight length as well as severe cold and wind conditions (Blix, 2016; Goldsmith and Sladen, 1961). To withstand such hostile environment, the Emperor penguin shows multiple morphological, physiological, behavioral adaptations, like improved thermoregulation systems in the head, wings, and legs (Frost et al., 1975; Thomas and Fordyce, 2008), and efficient energy storage management system for long-term fasting (Groscolas, 1990; Cherel et al., 1994; Groscolas and Robin, 2001). Conversely, its sister species, the King penguin (*A. patagonicus*), breeds exclusively in year-round ice-free sub-Antarctic islands and in Tierra del Fuego. Extreme cold adaptation in the Emperor penguin has been suggested to be a derived feature from a less cold-adapted ancestor, likely more ecologically similar to the King penguin (Vianna et al., 2020). Such marked ecological transition should have left a clear signature of selection change across the genome of the Emperor penguin, with some genes becoming the novel targets of positive selection while others getting released from previous selective pressures associated with the ancestral habitat.

Here, we apply phylogeny-based tests to identify genes that markedly changed in their selection regime during the evolutionary history of the Emperor penguin, using its less cold-adapted sister species, the King penguin, as a control. If the common ancestor ecology was similar to the King penguin one (*i.e*., not so cold-adapted), we expect a more intense selection shift across the Emperor penguin genome, with genes under positive selection related to adaptations to cold. By using a phylogenetic framework including seven species of penguins and 13 other birds, we compare the pattern of molecular evolution (Yang, 2007; Wertheim et al., 2015) between Emperor and King penguins across 7,651 orthologous genes and explore the gene ontology terms to identify the molecular functions that may have undergone positive selection in the Emperor penguin. To allow for a broader comparison with other cold-adapted vertebrates, we also investigated the overlap between the biological functions of the candidate genes identified in the Emperor penguin and the metabolic and physiological functions related to cold adaptation compiled from previous studies.

## Methods

### Orthologous coding sequences identification

In order to test for selective signatures in coding sequences of Emperor and King penguins, we implemented a comparative phylogenomic analysis. We selected at least one species for each extant penguin genus based on the phylogeny of Pan et al. (2019), and other bird species representing the clade of Core Waterbirds (which includes penguins), the tropicbirds (one species only), and some more distant species from the Core Landbirds according to Jarvis et al. (2014). The resulting dataset included seven penguin species (*Eudyptula minor minor*, *Spheniscus magellanicus*, *Eudyptes chrysolophus*, *Pygoscelis papua*, *Pygoscelis adeliae*, *Aptenodytes patagonicus* and *Aptenodytes forsteri*) and 13 additional bird species (*Phaethon lepturus*, *Eurypyga helias*, *Gavia stellata*, *Fulmarus glacialis*, *Phalacrocorax carbo*, *Nipponia nippon*, *Egretta garzetta*, *Pelecanus crispus*, *Haliaeetus leucocephalus*, *Tyto alba*, *Cariama cristata*, *Corvus brachyrhynchos* and, as a more distant outgroup, *Opisthocomus hoazin*). All coding sequences (CDS) of each of these twenty bird species were downloaded from GigaDB and Genbank (*SI Appendix*, Table S1).

One-to-one orthologs were identified by applying a reciprocal best-hit approach using pairwise BLAST searches with an e-value cutoff of 1e-15, a nucleotide sequence identity of at least 70%, and a fraction of aligned CDS of at least 60% (Ometto et al., 2013). Only CDS longer than 150 bp that were a reciprocal best-hit between the Emperor penguin and the other species were retained. The orthologous gene sequences were then aligned with MAFFT (Madeira et al., 2019) and the alignments trimmed to maintain the open reading frame using a custom perl script. The resulting nucleotide alignments were re-aligned using the PRANK algorithm (Löytynoja, 2013) in TranslatorX (Abascal et al., 2010), which aligns protein-coding sequences based on their corresponding amino acid translations. Since the results of *dN/dS* analysis could have been affected by poorly aligned regions or by regions that were too different to be considered truly orthologs, we removed CDS alignments that included internal stop codons and used a custom perl script to remove problematic regions as in Han et al. (2009) (see also Ramasamy et al., 2016). Furthermore, as phylogenetic-based selection tests are not able to properly deal with alignment gaps (Yang, 2007), we filtered all the alignments with a custom perl script and kept only sites that were unambiguously present in at least 16 of the 20 sequences and always present in both our species of interest (*i.e*., King and Emperor penguins). The minimum length of an alignment for subsequent analyses after gap removal was set to 150 bp (50 codons).

### Identification of selection regime shifts

We first used CODEML in the PAML package (Yang, 2007) to estimate synonymous (*dS*) and nonsynonymous (*dN*) substitution rates and to identify genes which are characterized by lineage-specific ω values (*i.e*., *dN/dS*, which is a proxy for the level of past selective pressure in the gene). In particular, we separately investigated the two scenarios of different ω in the King or in the Emperor lineage. If one branch in the phylogeny shows a value of ω (ω_i_) significantly larger than in the other branches (*i.e*., ω_b_; background ω), such a foreground lineage may have been targeted by positive (Darwinian) or relaxed selection (allowing the accumulation of non synonymous substitutions), whereas when ω_i_ is lower than ω_b_, the foreground lineage may have evolved under stronger selective constraints (*e.g*., purifying selection).

We first ran the two-ratio branch model (one ω for the foreground branch, another ω for the background branches; set parameters “model=2, NSsites=0, fix_omega=0”) and the one-ratio branch model (one ω estimate for all branches, as null model; set parameters “model=0, NSsites=0, fix_omega=0”) on the unrooted phylogenetic tree of the species of interest. We determined the topology of such a tree (Fig. 1A) by manually combining the total evidence nucleotide tree of the avian family (Jarvis et al., 2014) and the phylogenomic reconstruction of penguins (Pan et al., 2019). The two models (two-ratios vs. one-ratio) were compared by likelihood ratio tests (LRTs). False discovery rates (FDR) were computed using the qvalue package (Storey at al., 2017) and the p.adjust function in R (R Core Team, 2013) using the Benjamini-Hochberg procedure to adjust for multiple testing. An FDR or *p*-adjusted significant threshold of 0.05 was used.

**Figure 1.**
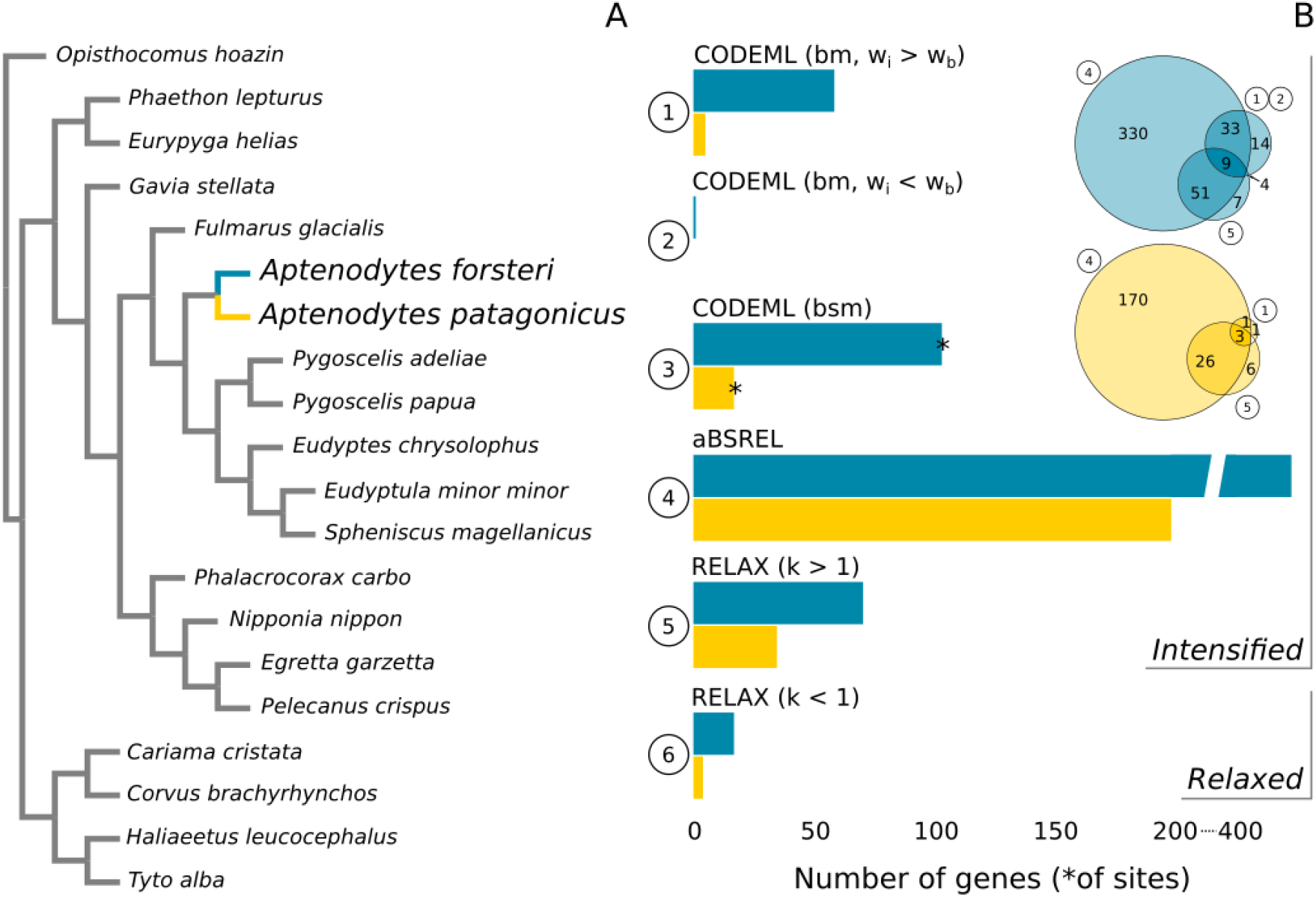
**A.** Phylogenetic tree based on Jarvis et al. 2014 and Pan et al. 2019. The Emperor and the King penguin are highlighted in blue and yellow, respectively. Note that branch length is not to scale. **B.** Comparison between Emperor and King penguins for genes with FDR > 0.05 in each of the tests performed; bm: branch model; bsm: branch-site model; w_i_: ω in the target species; w_b_: background ω; For sake of completeness, we also show the genes putatively under selection according to RELAX (with *K* > 1). **Inset**. Venn diagram showing the overlap among CODEML (bm), aBSREL, and RELAX (*K* > 1). Note that the total number of genes is different between the Emperor and the King penguins and the size of the circles scales to the maximum in each of the two graphs. The overlap between CODEML (bm) or/and aBSREL with RELAX (*K* < 1) is 3, 1, and 1 gene, respectively (not shown).

We then used a branch-site model to test for sites under selection in the candidate genes from the previous test. The parameters for the null model were set as “model=2, NSsites=2, fix_omega=1”, while the parameters for the alternative model were set as “model=2, NSsites=2, fix_omega=0”. LRT and FDR were computed as for the branch model tests. To control for misalignments that could have biased the results, we visually checked the sequence alignment of all candidate genes under selection using MEGA7 (Kumar et al., 2016).

We also applied aBSREL from the HyPhy package, a branch-site test that includes an adaptive branch-site random effects likelihood model (Smith et al., 2015; Pond et al., 2005), to our set of orthologous coding sequences. In contrast to the branch-site test in CODEML, which assumes 4 ω-rate classes for each branch and assigns each site to one of these classes, the aBSREL test uses AIC_c_ to infer the optimal number of ω-rate categories per branch, not making the assumption that all branches exhibit the same degree of substitution rate heterogeneity (Smith et al., 2015). Although we expect a broad overlap between CODEML and aBSREL results, a higher sensitivity should characterize the latter approach (Smith et al., 2015). Signatures of positive selection were searched by setting *a priori* the King and the Emperor lineage as *test* branches in the phylogeny. LRT was performed by comparing the full model to a null model where branches were not allowed to have rate classes of ω > 1. A Benjamini-Hochberg correction was used to control the probability of making false discoveries and only tests with adjusted p-values < 0.05 were considered significant.

Beside presenting novel drivers of selection, the major ecological shift occurring in the Emperor penguin lineage should have also released some of the selective constraints characterizing the ancestral ecological niche. As a consequence, some genes could show a signature of relaxed selection in this lineage, a higher number than in the King penguin. To test for relaxation of selective constraints on a specific lineage we used RELAX from the HyPhy package, a general hypothesis testing framework that determines whether the strength of natural selection has been relaxed or intensified along a set of test branches defined *a priori* in a phylogenetic tree (Wertheim et al., 2015; Pond et al., 2005). It estimates a selection intensity parameter *K*, in which a significant *K* > 1 indicates intensification in the selection strength, whereas a significant *K* < 1 indicates relaxation in the strength of selection in the test branches (Wertheim et al., 2015). We tested whether selection pressure has increased or decreased in either the King or in the Emperor lineage as compared to the rest of the phylogeny. In the null model, the selection intensity parameter *K* was set to 1 for all the branches of the phylogenetic tree, whereas in the alternative model the parameter *K* was inferred for every tested branch. The increase or relaxation of selection was validated by a LRT with 1 degree of freedom. Again, the Benjamini-Hochberg procedure was used to adjust for multiple testing with adjusted p-values < 0.05 considered significant.

### Functional characterization of candidate genes under selection

To test whether the set of candidate genes for positive selection in the Emperor penguin lineage has a relevance for cold adaptation, we tested these genes for functional GO terms enrichment by using the g:GOSt function in g:Profiler (Raudvere et al., 2019). GO terms were assigned to candidate genes based on the Ensembl GO predictions for the flycatcher (*Ficedula albicollis*). Significantly enriched categories included at least two genes, and the Benjamini-Hochberg method was used for multiple testing correction to estimate significance (at *p* < 0.05). We used REVIGO (Supek et al., 2011) to summarize the resulting lists of GO terms in order to obtain a non-redundant and more easily interpretable set of GO terms. GO terms enrichment was performed on two lists of genes: *i)* candidate genes supported by both CODEML and aBSREL and *ii)* candidate genes supported by either CODEML or aBSREL.

To compare our results with the recent literature on genetics of cold adaptation, we compiled a list of biological/molecular functions characterizing the candidate genes for cold adaptation retrieved in previous studies in vertebrates (Table 1). The list included: *cardiovascular activity and regulation, skin thickness, immunity, lipid and fatty acid metabolism, glucose (including insulin) metabolism, thyroid hormones, non-shivering thermogenesis, shivering thermogenesis, response to oxidative stress, stress response, homeostasis, circadian rhythm, phototransduction, mitochondrial activity, feathers development*. We checked whether any of the GO terms enriched in candidate genes supported by either CODEML or aBSREL could be assigned to any of the 15 biological/molecular functions listed above. In addition, we assigned, whenever possible, genes from this list to the same biological/molecular functions, using the gene function description from the human gene database GeneCards (Stelzer et al., 2016).

**Table 1.**
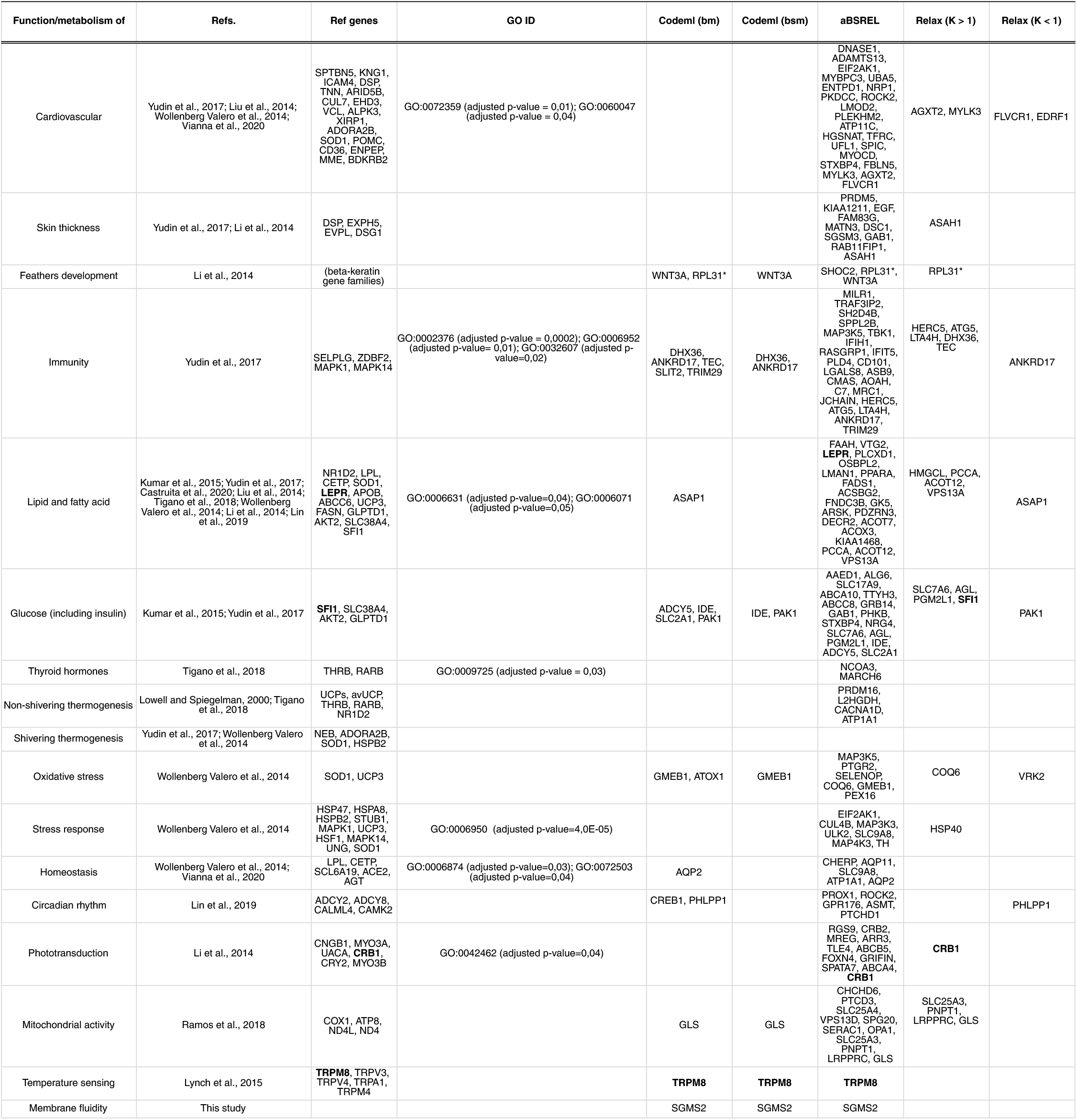
Candidate genes for positive/relaxed selection assigned to biological/molecular functions putatively related to cold adaptation from previous studies. In bold, genes identified as being under positive selection also in other cold adapted vertebrates. *Weak evidence of gene expression change in a study on feathers development (Ng et al., 2015).

## Results

### Signature of selection shift in the Emperor penguin lineage

We identified 7,651 orthologous coding sequences across seven penguin species and 13 other birds, corresponding to about 50% of the total number of genes in an avian genome (Zhang et al., 2014). Across all of the tests, by applying a significance threshold of FDR < 0.05, we consistently identified more candidate genes which underwent a shift in their selection regime (intensified or relaxed) in the Emperor penguin lineage than in the King penguin one (Fig. 1B). Even though we found a much larger number of genes putatively under selection using the aBSREL model (*SI Appendix*, Table S2, S3), the overlap between the CODEML (with ω_i_ > ω_b_) branch model (*SI Appendix*, Table S4) or RELAX (with *K* > 1; *SI Appendix*, Table S5, S6) with aBSREL was on average 80% (Fig. 1B, inset).

Using the CODEML branch test, we found 59 candidate genes with signals of positive or relaxed selection in the Emperor penguin, showing a ω significantly greater than the background, and one candidate gene under purifying selection showing a lower ω value than the background. In comparison, only five genes, with ω greater than the background, were retained as candidates of positive or relaxed selection in the King penguin lineage. Although greater than those of the background branches, the ω values of most of the candidate genes in the Emperor or the King penguin, still remain lower than one, making it difficult to distinguish between positive selection and relaxation of purifying selection. The CODEML branch-site test indicated a total of 104 sites in the 60 candidate genes in the Emperor lineage, whereas 17 were suggested in the five genes in the King lineage (Fig. 1B; *SI Appendix*, Table S7). On the other hand, aBSREL identified a much larger number of candidate genes under positive selection than CODEML branch model (423 in the Emperor lineage and 200 in the King lineage). Between 70% and 80% of the CODEML candidate genes (42 and 4 genes considering the Emperor and the King lineage, respectively) were also significant in aBSREL results (Fig. 1B). According to RELAX, 17 genes in the Emperor lineage and in four genes in the King lineage bear a significant signature of relaxed selection (K < 1; Fig. 1B; *SI Appendix*, Table S8, S9). Concerning the Emperor lineage, five of the 17 genes (*i.e*., FLVCR1, ANKRD17, ASAP1, PAK1, PHLPP1) are also candidates in either CODEML (branch model, ω_i_ > ω_b_), aBSREL, or both, further supporting the signal of relaxed purifying selection.

### Biological functions enrichment in Emperor penguin candidate genes

After correcting for multiple tests and using REVIGO’s redundancy elimination algorithm, we found 16 enriched GO biological process terms in candidate genes for positive selection suggested by both CODEML and aBSREL (*SI Appendix*, Table S10). Some of these GOs are related to heart and muscle development (GO:0003306, GO:1901863), lipid (GO:0033993), glucose (GO:0071333) and sphingolipid (*i.e*., ceramide) metabolism (GO:1905371). When considering all candidate genes supported by either CODEML or aBSREL, we retrieved 34 enriched GO biological process terms (*SI Appendix*, Table S11), 12 of which could be assigned to one of the biological/molecular functions putatively related to cold adaptation from previous studies (Table 1). In addition, qualitatively screening the biological functions of all candidate genes using the human database GeneCards, we identified 161 genes which could be assigned to the biological/molecular functions identified in previous studies (Table 1). Four genes identified as under selection in the Emperor penguin were already found in previous studies: TRPM8 (*temperature sensing*), indicated as under selection by both CODEML (brach and brach-site models) and aBSREL; LEPR (*lipid and fatty acid metabolism*) and CRB1 (*phototransduction*) were suggested as candidate genes under selection by aBSREL; SFI1 (*glucose metabolism*) which showed a significant signal of intensified selection in the RELAX test (K > 1). In addition, the sphingomyelin synthase2 (SGMS2), shows a biological function that appears as markedly related to cold adaptation in the Emperor penguin, potentially regulating biological membrane fluidity at low temperatures (Wang et al., 2014). This function (named as *membrane fluidity*) was added to the list of biological/molecular functions putatively related to cold adaptation (Table 1).

## Discussion

### Emperor penguin shift towards a novel ecological niche

Among the orthologous genes tested in our analyses, a larger fraction shows signatures of novel selection regimes in the Emperor penguin lineage than in the King penguin, either as intensification or relaxation of the selection pressure (Fig. 1B). One possible explanation of this pattern is that the ancestor of both species had ecological preferences which were more similar to the King penguin, while the Antarctic ecology of the Emperor penguin is a derived, though rather recent (ca. 1-2 Mya; Gavryushkina et al., 2017), adaptation. A general shift from warmer to colder habitats has been suggested for penguins in general (Vianna et al., 2020). King and Emperor penguins critically differ in their breeding range, as they reproduce in temperate-cold sub-Antarctic islands with average winter temperature of ca. 3 °C, or on the Antarctic sea-ice featuring average winter temperature around −25 °C, sometimes at −60 °C, respectively. Upon colonization of Antarctica, novel selective pressures are expected to have appeared while others, characterizing the former ecology, should have relaxed. Relaxed purifying selection can in fact be an additional source of evolutionary novelties (Hunt et al., 2011), as it can have non-linear consequences on a trait, including stabilizing or balancing selection, pseudogenization or, on the contrary, recruitment for a different function (Lathi et al., 2009).

Rather fast adaptation to polar lifestyle is not new in homeothermic vertebrates, as suggested for the recent divergence (less than 0.5 Mya) of the polar bear from the brown bear (Liu et al., 2014, Castruita et al., 2020), or of the Arctic fox from its common ancestor with red fox (ca. 2.9 Mya; Kumar et al., 2015). One question, which could be highly relevant in the ongoing climate change scenario, is whether extreme cold adaptation is an evolutionary *cul de sac*, *i.e*., a derived suite of traits from which reverting to a less extreme cold ecology is hampered. Interestingly, none of the extant rhino species, which are all adapted to warm climate, descend from any of the three cold-adapted species we know about and which are all extinct (Liu et al., 2021). A similar evolutionary endpoint could have characterized the diversification of elephants, with the extinction of the cold-adapted mammoths (Lynch et al., 2015).

An alternative explanation for the higher number of genes showing evidence of selection shifts in the Emperor penguin is that selection was more efficient along the Emperor lineage due to its larger and more constant population size through time, as revealed in previous studies (Trucchi et al., 2014, Cristofari et al., 2016), when compared to the King penguin markedly oscillating demographic trajectory (Cristofari et al., 2018, Trucchi et al., 2019). In fact, at low population size genetic drift overwhelms selection, leading to the fixation of both synonymous and nonsynonymous variants, and potentially blurring the *dN/dS* ratios. However, the higher signature of relaxed purifying selection in the Emperor penguin contrasts with this alternative explanation.

### Looking for a common genetic underlying of adaptation to cold in homeotherms

A large fraction of the candidate genes under selection in the Emperor penguin are involved in most of the functional pathways relevant to cold adaptation identified in previous studies (Table 1). Nonetheless, we discovered only four genes in common with those previously identified in other cold-adapted vertebrates (notably, two of which were found in the mammoth). This result is not surprising given the suggested polygenic basis of most phenotypic traits (Barghi et al., 2020) which could be shaped by the contribution of a quite large set of genes (Boyle et al., 2017). As also emerged in other vertebrates (see refs in Table 1), traits related to cardiovascular function, lipid and fatty acid metabolism, glucose metabolism, oxidative stress and stress response, insulation (including skin thickness and feathers development), phototransduction and mitochondrial activity show the largest number of candidate genes under selection in the Emperor penguin (Table 1).

Genes involved in fatty-acid metabolism have been identified to be under positive selection both in polar bear and Arctic fox, indicating similar evolutionary constraints on fat metabolism in these two cold-adapted mammal species (Liu et al 2014; Kumar et al., 2015). The storage of subcutaneous fat is also crucial in the Emperor penguin, both because it represents the main source of energy during the long fasting periods (Blem, 1990; Cherel et al., 1994; Groscolas et al., 1990) and because it provides thermal insulation (Kooyman et al., 1976). Moreover, fatty-acids can stimulate muscle thermogenic processes in birds and therefore may be a very important component of the adaptive response to cold temperatures in penguins (Duchamp et al., 1999; Toyomizu et al., 2002; Talbot et al., 2004; Rey et al., 2010). Accordingly, our results revealed candidate genes under selection in the Emperor penguin such as PPARa, regulating eating behavior (Fu et al., 2003), controlling lipid absorption in the intestine (Poirier et al., 2001) and fatty-acid oxidation (Lemberger et al., 1996), ADCY5, associated with body weight (Li and Li, 2019), and LEPR, the leptin receptor. LEPR is involved in fat and glucose metabolism, in appetite-regulating through its effects on food intake and energy consumption (Zhang et al., 1994; Halaas et al., 1995; Pelleymounter et al., 1995), and in adaptive thermogenesis (Yang et al., 2011). Yet, the adipostat activity of leptin has not been demonstrated in Aves, where the expression pattern differs greatly from that of mammals, as it seems to be missing from the adipose tissue (Friedman-Einat and Eyal Seroussi, 2019).

While shivering thermogenesis might be the main thermogenic mechanism in birds following short-term cold exposure (Teulier et al., 2010), cold acclimated birds show non-shivering thermogenesis mediated by avUCP expression within the skeletal muscle (Talbot et al., 2004). According to our analyses, some of the candidate genes could be assigned to the non-shivering thermogenesis category (Table 1), but none of them can be unambiguously associated with shivering thermogenesis. Among the former, Na,K-ATPase (ATP1A1) is a membrane enzyme that utilizes energy derived from the hydrolysis of ATP to pump Na+, wasting energy as heat, thus playing a significant role in thermal tolerance and energy balance (Geering et al., 1987; Iannello et al., 2007). It was demonstrated that its expression is affected by heat stress (Sonna et al., 2002) and consistently increases during mammal hibernation (Vermillion et al., 2015). L2HGDH was found to be associated with the TCA cycle, electron transport and glycolysis (Oldham et al., 2006) and it was identified as one of the candidate genes under positive selection in three high-altitude passerine birds (Hao et al., 2019). PRDM16 is a zinc-finger protein that activates brown fat-selective genes responsible for mitochondrial biogenesis and oxidative metabolism, while repressing the expression of a wide range of genes selective for white fat cells (Seale et al., 2007; Kajimura et al., 2015). This protein appears to play a role also in the development and function of beige cells (Ohno et al., 2012; Seale et al., 2011). Retinoic acid and thyroid hormones, whose candidate binding sites have been found in a mammal UCP-1 enhancer (Lowell and Spiegelman, 2000). These hormones, together with another nuclear receptor (NR1D2), have been suggested to be involved in thermal adaptation in birds (Tigano et al., 2018). NR1D2 as well as MARCH6 and NCOA3, also involved in thyroid hormone regulation and action (Zelcer et al., 2014; Ishii et al., 2021), influencing baseline temperature (Elliott et al., 2013) and thermoregulation in response to cold stimuli in birds (Vézina et al., 2015), were present among our candidate genes.

One of the most promising candidate genes under selection is TRPM8, which encodes the sensor for noxious cold temperature (Yin et al., 2018). Setting the physiological range of temperature tolerance and, ultimately, the width of the geographical habitat (Matos-Cruz et al., 2017), any biological thermosensory apparatus should be under strong evolutionary pressures to trigger specific responses to noxious high or low temperatures (Myers et al., 2009). Indeed, evolutionary tuning of five temperature-sensitive transient receptor potential channels, including TRPM8, has been likely key in the adaptation of the Woolly mammoth to the Arctic (Lynch et al., 2015). A previous study demonstrated the crucial role of a single-point mutation located at site 906 (as per *Gallus gallus* coordinates in Yin et al., 2018; 919 as per coordinates in Yang et al., 2020) for the activation of TRPM8 pore domain channel in the Emperor penguin (Yang et al., 2020). Interestingly our comparative selection scan found instead evidence of positive selection at two other sites (*i.e*., I1058T, Y1069M). After aligning 541 vertebrates TRPM8 ortholog sequences available in GenBank (accessed on 22/11/2021), we found that the substitution at site 906 is not unique to Emperor penguins but it is instead widespread in birds with different ecology and habitat preference, including warm tropical regions. Also our candidate substitution Y1069M is common in penguins and other birds. Conversely, the substitution I1058T (as per *Gallus gallus* coordinates in Yin et al., 2018) is extremely rare in birds, being present in one other species only (*Sitta europea*). This substitution is also rare in mammals and reptiles where it appears in seven and one species only, respectively. We located the single-point mutation I1058T just after the last ultra-conserved residue in one of the three helices (CTDH2) of the C-terminal cytoplasmic domain (Yin et al., 2018). Further analyses should be conducted to determine if and how this substitution affects noxious cold sensing in Emperor penguins.

### Limitations of this study

There is no overlap between our set of candidate genes under selection and those discovered in previous studies on penguins (Li et al., 2014; Vianna et al., 2020). On one hand, selection tests based on phylogenies are, of course, influenced by size and composition of the set of species included in the tree. In contrast with Li et al. (2014), where 48 bird species of which only two penguins were analyzed, we selected non-target species both in close (seven penguins) and in more distant (13 other birds) clades. On the other hand, the bioinformatic pipeline applied for identifying ortholog coding sequences across the species in the phylogeny may determine which genes are included or excluded. Our pipeline, successfully tested in *Drosophila* (Ometto et al., 2013), aligned ca. 30% more genes than in Vianna et al. (2020) and slightly less than in Li et al. (2014). We also note that in Vianna et al. (2020), the selection scan was performed for 18 penguin species, not only on the King and Emperor ones, and searched for candidate genes in all of the penguin lineages.

Although we found an appreciable overlap between the results of CODEML and aBSREL, the latter suggested a lot more candidate genes (Fig. 1B). This could be due either to higher sensitivity of aBSREL, as due to the branch-site model applied, or to a higher false negative rate in CODEML. One big difference between these two methods is that aBSREL makes no assumptions about the selective regime on background branches while CODEML assumes negative or no selection on background branches. This could lead to a higher false negative rate when the evolutionary process along background branches deviates significantly from modeling assumptions (Kosakovsky Pond et al., 2009). In general, phylogenetic tests of adaptive evolution are not capable of avoiding false positives because they do not consider multi nucleotide mutations (simultaneous mutations at two or three codon positions) which can instead be more common than expected (Venkat et al., 2018).

Concerning our characterization of the candidate genes to associate them to the functional categories we identified from previous studies on cold adaptation, we reckon that this is by far a simple (*i.e*., gene characterization was based on GeneCards description only) and qualitative (*i.e*., not based on a statistical test) comparison, aiming at contextualizing our results within the published literature. As sequence orthology does not necessarily equal functional orthology, we acknowledge that a functional characterization largely based on a human model (and sometimes on a chicken model) is a strong assumption for a distant non-model organism.

## Acknowledgements

ET was supported by the PNRA16_00164 (“Programma Nazionale di Ricerca in Antartide. Bando PNRA 5 aprile 2016, n. 651. – Linea B “Genomica degli adattamenti estremi alla vita in Antartide”).

## Notes

### Competing Interest Statement

The authors have declared no competing interest.

## References

Abascal, F., Zardoya, R. & Telford, M.J. 2010 TranslatorX: multiple alignment of nucleotide sequences guided by amino acid translations. Nucleic Acids Research 38, 7–13.

Akashi, H.D., Cádiz Díaz, A., Shigenobu, S., Makino, T. & Kawata, M. 2016 Differentially expressed genes associated with adaptation to different thermal environments in three sympatric Cuban Anolis lizards. Molecular Ecology 25, 2273–2285.

Allen, J.A. 1877 The influence of Physical conditions in the genesis of species. Radical Review 1, 108–140.

Barghi, N., Hermisson, J. & Schlötterer, C. 2020 Polygenic adaptation: a unifying framework to understand positive selection. Nature Reviews Genetics 21, 769–781.

Blem, C.R. 1990 Avian energy storage. Current Ornithology 7, 59–113.

Blix, A.S. 2016 Adaptations to polar life in mammals and birds. Journal of Experimental Biology 219, 1093–1105.

Boyle, E.A., Li, Y.I. & Pritchard J.K. 2017 An Expanded View of Complex Traits: From Polygenic to Omnigenic. Cell 169, 1177–1186.

Cannon, B. & Nedergaard, J. 2010 Thyroid hormones: igniting brown fat via the brain. Nature Medicine 16, 965–967.

Castruita, J.A.S., Westbury, M.V. & Lorenzen, E.D. 2020 Analyses of key genes involved in Arctic adaptation in polar bears suggest selection on both standing variation and de novo mutations played an important role. BMC Genomics 21, 1–8.

Cherel, Y., Gilles, J., Handrich, Y. & Le Maho, Y. 1994 Nutrient reserve dynamics and energetics during long-term fasting in the king penguin (Aptenodytes patagonicus). Journal of Zoology 234, 1–12.

Cristofari, R., Bertorelle, G., Ancel, A., et al. 2016 Full circumpolar migration ensures evolutionary utility in the Emperor penguin. Nature Communications 7, 1–9.

Cristofari, R., Liu, X., Bonadonna, F., et al. 2018 Climate-driven range shifts of the king penguin in a fragmented ecosystem. Nature Climate Change 8, 245–251.

Duchamp, C., Marmonier, F., Denjean, F., Lachuer, J., et al. 1999 Regulatory, cellular and molecular aspects of avian muscle non-shivering thermogenesis. Ornis Fennica 76, 151–165.

Elliott, K.H., Welcker, J., Gaston, A.J., et al 2013 Thyroid hormones correlate with resting metabolic rate, not daily energy expenditure, in two charadriiform seabirds. Biology Open 2, 580–586.

Friedman-Einat, M. & Seroussi, E. 2019 Avian Leptin: Bird’s-Eye View of the Evolution of Vertebrate Energy-Balance Control. Trends in Endocrinology and Metabolism 30, 819–832.

Frost, P.G.H., Siegfried, W.R. & Greenwood, P.J., 1975 Arterio-venous heat exchange systems in the Jackass penguin Spheniscus demersus. Journal of Zoology 175, 231–241.

Fu, J., Gaetani, S., Oveisi, F., Lo Verme, J., et al. 2003 Oleylethanolamide regulates feeding and body weight through activation of the nuclear receptor PPAR-alpha. Nature 425, 90–93.

Gavryushkina, A., Heath, T.A., Ksepka, D.T., Stadler, T., et al. 2017 Bayesian total-evidence dating reveals the recent crown radiation of penguins. Systematic biology 66, 57–73.

Geering, K., Kraehenbuhl, J.P. & Rossier, B.C. 1987 Maturation of the catalytic alpha unit of Na, K-ATPase during intracellular transport. Journal of Cell Biology 105, 2613–2619.

Goldsmith, R. & Sladen, W.J. 1961 Temperature regulation of some Antarctic penguins. The Journal of Physiology 157, 251–262.

Groscolas, R. 1990 Metabolic adaptations to fasting in emperor and king penguins. In Penguin Biology (ed. LS Davis & JT Darby), 269–296. San Diego: Academic Press.

Groscolas, R. & Robin, J.P., 2001 Long-term fasting and re-feeding in penguins. Comparative Biochemistry and Physiology Part A: Molecular & Integrative Physiology 128, 645–655.

Halaas, J.L., Gajiwala, K.S., Maffei, M., Cohen, S.L., et al. 1995 Weight-reducing effects of the plasma protein encoded by the obese gene. Science 269, 543–546.

Han, M.V., Demuth, J.P., McGrath, C.L., Casola, C. & Hahn, M.W. 2009 Adaptive evolution of young gene duplicates in mammals. Genome Research 19, 859–867.

Hao, Y., Xiong, Y., Cheng, Y., Song, G., et al. 2019 Comparative transcriptomics of 3 high-altitude passerine birds and their low-altitude relatives. PNAS 116, 11851–11856.

Hunt, B.G., Ometto, L., Wurmb, Y., et al. 2011 Relaxed selection is a precursor to the evolution of phenotypic plasticity. PNAS 108, 15936–15941.

Iannello, S., Milazzo, P. & Belfiore, F. 2007 Animal and human tissue Na,K-ATPase in normal and insulin-resistant states: regulation, behaviour and interpretative hypothesis on NEFA effects. Obesity Reviews 8, 231–51.

Ishii, S., Amano, I., & Koibuchi, N. 2021 The Role of Thyroid Hormone in the Regulation of Cerebellar Development. Endocrinology and Metabolism 36, 703–716.

Jarvis, E.D., Mirarab, S., Aberer, A.J., et al. 2014 Whole genome analyses resolve early branches in the tree of life of modern birds. Science 346, 1320–1331.

Kajimura, S., Spiegelman, B.M. & Seale, P. 2015 Brown and Beige Fat: Physiological Roles beyond Heat Generation. Cell Metabolism 22, 546–559.

Kooyman, G.L., Gentry, R.L., Bergman W.P. & Hammel, H. T. 1976 Heat loss in penguins during immersion and compression. Comparative Biochemistry and Physiology Part A 54, 75–80.

Kosakovsky Pond, S.L., Poon, A.F.Y. & Frost, S.D.W. 2009 Estimating selection pressures on alignments of coding sequences. The phylogenetic handbook: a practical approach to phylogenetic analysis and hypothesis testing. Cambridge Univ. Press, Cambridge, UK, 419–490.

Kumar, V., Kutschera, V.E. & Nilsson, M.A. 2015 Genetic signatures of adaptation revealed from transcriptome sequencing of Arctic and red foxes. BMC Genomics 16, 1–13.

Kumar, S., Stecher, G. & Tamura, K. 2016 MEGA7: Molecular Evolutionary Genetics Analysis version 7.0 for bigger datasets. Molecular Biology and Evolution 33, 1870–1874.

Lemberger, T., Saladin, R., Vazquez, M., Assimacopoulos, F., et al. 1996 Expression of the peroxisome proliferator-activated receptor alpha gene is stimulated by stress and follows a diurnal rhythm. Journal of Biological Chemistry 271, 1764–1769.

Li, F.G. & Li, H. 2019 A time-dependent genome-wide SNP-SNP interaction analysis of chicken body weight. BMC Genomics 20, 1–9.

Li, C., Zhang, Y., Li, J. et al. 2014 Two Antarctic penguin genomes reveal insights into their evolutionary history and molecular changes related to the Antarctic environment. Gigascience 3, 2047–2217X.

Lin, Z., Chen, L., Chen, X., Zhong, Y., Yang, Y., Xia, W., Liu, C., Zhu, W., Wang, H., Yan, B. & Yang, Y. 2019 Biological adaptations in the Arctic cervid, the reindeer (Rangifer tarandus). Science 364(6446).

Liu, S., Lorenzen, E.D., Fumagalli, M., et al. 2014 Population genomics reveal recent speciation and rapid evolutionary adaptation in polar bears. Cell 157, 785–794.

Lowell, B.B. & Spiegelman, B.M. 2000 Towards a molecular understanding of adaptive thermogenesis. Nature 404, 652–660.

Löytynoja, A. 2013 Phylogeny-aware alignment with PRANK. Multiple Sequence Alignment Methods. Methods in Molecular Biology (Methods and Protocols) 1079, 155–170.

Lynch, V.J, Bedoya-Reina, O.C., Ratan, A., Sulak, M., Drautz-Moses, D.I., et al. 2015 Elephantid genomes reveal the molecular bases of woolly mammoth adaptations to the Arctic. Cell Reports 12, 217–228.

Madeira, F., Park, Y.M., Lee J., et al. 2019 The EMBL-EBI search and sequence analysis tools APIs in 2019. Nucleic Acids Research 47, 636–641.

Matos-Cruz, V., Schneider, E.R., Mastrotto, M., et al. 2017 Molecular Prerequisites for Diminished Cold Sensitivity in Ground Squirrels and Hamsters. Cell Reports 21, 3329–3337.

Myers, B.R., Sigal, Y.M. & Julius D. 2009 Evolution of Thermal Response Properties in a Cold-Activated TRP Channel. PloS one 4, e5741.

Ng, C.S., Chen, C.K., Fan, W.L., Wu, P., Wu, S.M., Chen, J.J., Lai, Y.T., Mao, C.T., Lu, M.Y.J., Chen, D.R. & Lin, Z.S. 2015 Transcriptomic analyses of regenerating adult feathers in chicken. BMC Genomics 16, 1–16.

Ohno, H., Shinoda, K., Spiegelman, B.M. & Kajimura, S. 2012 PPARg agonists induce a white-to-brown fat conversion through stabilization of PRDM16 protein. Cell Metabolism 15, 395–404.

Oldham, M.C., Horvath, S. & Geschwind, D. H. 2006 Conservation and evolution of gene coexpression networks in human and chimpanzee brains. PNAS 103, 17973–17978.

Ometto, L., Cestaro, A., Ramasamy S., et al. 2013 Linking Genomics and Ecology to Investigate the Complex Evolution of an Invasive Drosophila Pest. Genome Biology and Evolution 5, 745–757.

Pan, H., Cole, T.L., Bi, X., et al. 2019 High-coverage genomes to elucidate the evolution of penguins. GigaScience 8, 1–17.

Pelleymounter, M.A., Cullen, M.J., Baker, M.B., Hecht, R., et al. 1995 Effects of the obese gene product on body weight regulation in ob/ob mice. Science 269, 540–543.

Poirier, H., Niot, I., Monnot, M.C., Braissant, O., et al. 2001 Differential involvement of peroxi-some-proliferator-activated receptors alpha and delta in fibrate and fatty-acid-mediated inductions of the gene encoding liver fatty-acid-binding protein in the liver and the small intestine. Biochemical Journal 355, 481–488.

Pond, S.K., Frost, S. & Muse, S.V. 2005 HyPhy: hypothesis testing using phylogenies. Bioinformatics 21, 676–679.

R Core Team. 2013 R: A language and environment for statistical computing, R Foundation for Statistical Computing, Vienna, Austria. http://www.R-project.org/.

Ramasamy, S., Ometto, L., Crava, C.M., Revadi, S., Kaur, R., Horner, D.S., et al. 2016 The evolution of olfactory gene families in Drosophila and the genomic basis of chemical-ecological adaptation in Drosophila suzukii. Genome Biology and Evolution 8, 2297–2311.

Ramos, B., González-Acuña, D., Loyola, D.E., et al. 2018 Landscape genomics: natural selection drives the evolution of mitogenome in penguins. BMC Genomics 19, 1–17.

Raudvere, U., Kolberg, L., Kuzmin, I. et al. 2019 g:Profiler: a web server for functional enrichment analysis and conversions of gene lists. Nucleic Acids Research 47, W191–W198.

Rey, B., Roussel, D., Romestaing, C., Belouze, M., et al. 2010 Up-regulation of avian uncoupling protein in cold-acclimated and hyperthyroid ducklings prevents reactive oxygen species production by skeletal muscle mitochondria. BMC Physiology 10, 1–12.

Roussel, D., Le Coadic, M., Rouanet, J.L. & Duchamp, C. 2020 Skeletal muscle metabolism in sea-acclimatized king penguins. I. Thermogenic mechanisms. Journal of Experimental Biology 223, p.jeb233668.

Rowland, L.A., Bal, N.C., & Periasamy, M. 2015 The role of skeletal-muscle-based thermogenic mechanisms in vertebrate endothermy. Biological Reviews 90, 1279–1297.

Scholander, P.F. 1955 Evolution of climatic adaptation in homeotherms. Evolution 9, 15–26.

Seale, P., Kajimura, S., Yang, W., Chin, S., et al. 2007 Transcriptional control of brown fat determination by PRDM16. Cell Metabolism 6, 38–54.

Seale, P., Conroe, H.M., Estall, J., Kajimura, S., et al. 2011 Prdm16 determines the thermogenic program of subcutaneous white adipose tissue in mice. Journal of Clinical Investigation 121, 96–105.

Smith, M.D., Wertheim, J.O., Weaver, S. 2015 Less Is More: An Adaptive Branch-Site Random Effects Model for Efficient Detection of Episodic Diversifying Selection. Molecular Biology and Evolution 32, 1342–1353.

Sonna, L.A., Fujita, J., Gaffin, S.L. & Lilly C.M. 2002 Effects of heat and cold stress on mammalian gene expression. Journal of Applied Physiology 92, 1725–1742.

Stelzer, G., Rosen, R., Plaschkes, I., Zimmerman, S., et al. The GeneCards Suite: From Gene Data Mining to Disease Genome Sequence Analysis. Current Protocols in Bioinformatics 54, 1–30.

Storey, K.B. & Storey J.M. 1992 Natural freeze tolerance in ectothermic vertebrates. Annual review of physiology 54, 619–637.

Storey, J.D., Bass, A.J., Dabney, A. & Robinson D. 2017 qvalue: Q-value estimation for false discovery rate control. R package version 2.15.0. http://github.com/StoreyLab/qvalue

Supek, F., Bošnjak, M., Škunca, N. & Šmuc T. 2011 REVIGO summarizes and visualizes long lists of gene ontology terms. PloS one 6, e21800.

Talbot, D., Duchamp, C., Rey, B. et al. 2004 Uncoupling protein and ATP/ADP carrier increase mitochondrial proton conductance after cold adaptation of king penguins. Journal of physiology 558, 123–135.

Tattersall, G.J., Sinclair, B.J., Withers, P.C., et al. 2012 Coping with thermal challenges: physiological adaptations to environmental temperatures. Comprehensive Physiology 2, 2151–202.

Teulier, L., Rouanet, J.L., Letexier, D. & Romestaing, C. 2010 Cold-acclimation-induced non-shivering thermogenesis in birds is associated with upregulation of avian UCP but not with innate uncoupling or altered ATP efficiency. The Journal of Experimental Biology 213, 2476–2482.

Thomas, D.B. & Fordyce R.E. 2008 The heterothermic loophole exploited by penguins. Australian Journal of Zoology 55, 317–321.

Tigano, A., Reiertsen, T.K., Walters, J.R., Friesen, V.L. 2018 A complex copy number variant underlies differences in both colour plumage and cold adaptation in a dimorphic seabird. BioRxiv, 507384.

Toyomizu, M., Ueda, M., Sato, S., Seki, Y., et al. 2002 Cold-induced mitochondrial uncoupling and expression of chicken UCP and ANT mRNA in chicken skeletal muscle. FEBS Letters 529, 313–318.

Trucchi, E., Gratton, P., Whittington, J.D., et al. 2014 King penguin demography since the last glaciation inferred from genome-wide data. Proceedings of the Royal Society B: Biological Sciences 281, 20140528.

Trucchi, E., Cristofari, R. & Le Bohec, C. 2019 Reply to:’The role of ocean dynamics in king penguin range estimation. Nature Climate Change 9, 122–122.

Venkat, A., Hahn, M.W. & Thornton, J.W. 2018 Multinucleotide mutations cause false inferences of lineage-specific positive selection. Nature Ecology and Evolution 2, 1280–1288.

Vermillion, K.L., Anderson, K.J., Hampton, M. & Andrews, M.T. 2015 Gene expression changes controlling distinct adaptations in the heart and skeletal muscle of a hibernating mammal. Physiological Genomics 47, 58–74.

Vézina, F., Gustowska, A., Jalvingh, K.M., Chastel, O. & Piersma T., 2015 Hormonal correlates and thermoregulatory consequences of molting on metabolic rate in a northerly wintering shorebird. Physiological and biochemical zoology 82, 129–142.

Vianna, J.A., Fernandes, F., Frugone M.J., et al. 2020 Genome-wide analyses reveal drivers of penguin diversification. PNAS 117, 22303–22310.

Wang, Q., Tan, X. & Jiao S. 2014 Analyzing Cold Tolerance Mechanism in Transgenic Zebrafish (Danio rerio). PloS one 9, e102492.

Wertheim, J.O., Murrell, B., Smith, M.D., Kosakovsky Pond, S.L. & Scheffler K. 2015 RELAX: detecting relaxed selection in a phylogenetic framework. Molecular biology and evolution 32, 820–832.

Wollenberg Valero, K.C., Pathak, R., Prajapati, I., Bankston, S., et al. 2014 A candidate multimodal functional genetic network for thermal adaptation. PeerJ 2, e578.

Yang, Z. 2007 PAML 4: a program package for phylogenetic analysis by maximum likelihood. Molecular Biology and Evolution 24, 1586–1591.

Yang, J., Bromage, T.G., Zhao, Q., Xu, B.H., Gao, W.L., Tian, H.F., Tang, H.J., Liu, D.W. & Zhao, X.Q. 2011 Functional evolution of leptin of Ochotona curzoniae in adaptive thermogenesis driven by cold environmental stress. PloS one 6, e19833.

Yang, S., Lu, X., Wang, Y., et al. 2020 A paradigm of thermal adaptation in penguins and elephants by tuning cold activation in TRPM8. PNAS 117, 8633–8638.

Yin, Y., Wu, M., Zubcevic, L., et al. 2018 Structure of the cold- and menthol-sensing ion channel TRPM8. Science 359, 237–241.

Yudin, N.S., Larkin, D.M. & Ignatieva E.V. 2017 A compendium and functional characterization of mammalian genes involved in adaptation to Arctic or Antarctic environments. BMC Genetics 18, 33–43.

Zelcer, N., Sharpe, L.J., Loregger, A., Kristiana, I., Cook, E.C., Phan, L., Stevenson J. & Brown A.J. 2014 The E3 ubiquitin ligase MARCH6 degrades squalene monooxygenase and affects 3-hydroxy-3-methyl-glutaryl coenzyme A reductase and the cholesterol synthesis pathway. Molecular and cellular biology 34, 1262–1270.

Zhang, Y., Proenca, R., Maffei, M., Barone, M., et al. 1994 Positional cloning of the mouse obese gene and its human homologue. Nature 372, 425–432.

Zhang, G., Li, C., Li, Q., et al. 2014 Comparative genomics reveals insights into avian genome evolution and adaptation. Science 346, 1311–1320.

