## Supplementary Material for "Selection-driven adaptation to the extreme Antarctic environment in the Emperor penguin"

**This PDF file includes:**
Tables S1 to S11

**Table S1. GigaDB and Genbank accession ID of the coding sequences (CDS) of the twenty bird species selected for the phylogeny**

| Species | Accession ID | Database |
|---|---|---|
| *Phaethon lepturus* | GCA_000687285.1 | GenBank |
| *Eurypyga helias* | GCA_000690775.1 | GenBank |
| *Gavia stellata* | GCA_000690875.1 | GenBank |
| *Fulmarus glacialis* | GCA_000690835.1 | GenBank |
| *Phalacrocorax carbo* | GCA_000708925.1 | GenBank |
| *Nipponia nippon* | GCA_000708225.1 | GenBank |
| *Egretta garzetta* | GCA_000687185.1 | GenBank |
| *Pelecanus crispus* | GCA_000687375.1 | GenBank |
| *Haliaeetus leucocephalus* | GCA_000737465.1 | GenBank |
| *Tyto alba* | GCA_000687205.1 | GenBank |
| *Cariama cristata* | GCA_000690535.1 | GenBank |
| *Corvus brachyrhynchos* | GCA_000691975.1 | GenBank |
| *Opisthocomus hoazin* | GCA_000692075.1 | GenBank |
| *Aptenodytes forsteri* | Aptenodytes_forsteri | GigaDB |
| *Aptenodytes patagonicus* | KP FORT 001 | GigaDB |
| *Pygoscelis adeliae* | Pygoscelis_adeliae | GigaDB |
| *Pygoscelis papua* | Gentoo penguin DNA -4 | GigaDB |
| *Eudyptula minor minor* | Gonzo | GigaDB |
| *Spheniscus magellanicus* | AH 6 | GigaDB |
| *Eudyptes chrysolophus* | MP PEI 1 | GigaDB |

Table S2. Candidate genes predicted to be under positive selection by aBSREL models in *A. forsteri* (FDR < 0.05)

| Gene ID | Gene name | Full adaptive model | Full adaptive model (non-synonymous subs/site) | Full adaptive model (synonymous subs/site) | LRT | Adjusted p-value |
|---|---|---|---|---|---|---|
| AF_67 | RAB11FIP1 | 0,03978832569 | 0,03756351761 | 0,002224808077 | 15,23266442 | 0,004735497959 |
| AF_126 | SIX4 | 0,3441046301 | 0,343446601 | 0,0006580290316 | 14,22141505 | 0,007197335296 |
| AF_224 | RECK | 0,1463463798 | 0,1459333673 | 0,0004130124426 | 24,53948108 | 0,0001031920533 |
| AF_262 | CTNNB1 | 0,1493238387 | 0,1491977367 | 0,0001261019944 | 22,04681764 | 0,0002728009514 |
| AF_284 | MATN3 | 18,92142651 | 18,91413236 | 0,007294147998 | 16,11095243 | 0,003275988575 |
| AF_393 | AAED1 | 0,01953908628 | 0,01932694688 | 0,0002121394001 | 10,96131878 | 0,02844513253 |
| AF_407 | DAPK1 | 0,8098414789 | 0,8086136977 | 0,001227781231 | 44,15418911 | 0,00000003670802221 |
| AF_471 | HFM1 | 0,00879112879 | 0,007957081776 | 0,0008340470144 | 15,8110221 | 0,003733917226 |
| AF_475 | ZNF326 | 0,01009790176 | 0,009474285168 | 0,0006236165973 | 10,90570251 | 0,02908619585 |
| AF_505 | VTG2 | 0,01056709205 | 0,009231676141 | 0,001335415905 | 12,3313369 | 0,01597452425 |
| AF_532 | FAAH | 0,1486163283 | 0,148387298 | 0,0002290302404 | 19,72381036 | 0,0006899259259 |
| AF_539 | ATP1A1 | 0,0045410133 | 0,003540288914 | 0,001000724387 | 19,40315246 | 0,0007940719858 |
| AF_569 | BLVRA | 0,009176526122 | 0,009169842134 | 0,000006683988159 | 13,1744513 | 0,01108689193 |
| AF_617 | ORAI2 | 0,1204426367 | 0,1191761458 | 0,001266490874 | 22,835274 | 0,0002067553426 |
| AF_653 | CHD1 | 0,00366681623 | 0,003661335911 | 0,000005480319194 | 13,31745026 | 0,01041231951 |
| AF_753 | SLCO2B1 | 1,53892817 | 1,537672809 | 0,001255361024 | 22,45030622 | 0,000233533378 |
| AF_755 | PGM2L1 | 2,548265627 | 2,546378343 | 0,001887283342 | 14,48694696 | 0,006529131699 |
| AF_767 | KIAA1211L | 0,05639786074 | 0,05358806488 | 0,002809795854 | 19,32306804 | 0,0008225005249 |
| AF_798 | EEF2KMT | 7,480391256 | 7,47683077 | 0,003560485915 | 32,43752956 | 0,000003532612265 |
| AF_812 | CCDC78 | 0,4645971468 | 0,4620782485 | 0,002518898309 | 40,67091371 | 0,0000001141742667 |
| AF_827 | DECR2 | 1,127009506 | 1,1266572 | 0,0003523059627 | 12,56959359 | 0,01433239235 |
| AF_864 | DNASE1 | 0,08281133932 | 0,07238054746 | 0,01043079186 | 13,78534023 | 0,00868524499 |
| AF_867 | CLCN7 | 0,02132897875 | 0,01988934777 | 0,001439630972 | 31,85409517 | 0,000004434647767 |
| AF_964 | CCNL1 | 0,142891058 | 0,1422133439 | 0,0006777140744 | 15,15261744 | 0,004869908448 |
| AF_1044 | ACMSD | 6,074454798 | 6,07150117 | 0,002953628114 | 10,97731619 | 0,02834500441 |
| AF_1081 | UBA5 | 0,02760754756 | 0,02759661161 | 0,00001093594528 | 17,17620541 | 0,0020476022 |
| AF_1154 | TRAIP | 0,02395098152 | 0,02108769185 | 0,002863289671 | 15,19354682 | 0,004811779067 |
| AF_1397 | WDR91 | 1,151496155 | 1,150012696 | 0,001483458679 | 15,31660935 | 0,004590771994 |
| AF_1408 | NUP205 | 0,5655074625 | 0,5635384856 | 0,001968976899 | 24,16836947 | 0,0001187902981 |
| AF_1424 | SPPL2B | 0,09318646639 | 0,09309602034 | 0,00009044604672 | 28,79132923 | 0,00001635403894 |
| AF_1497 | ATG5 | 0,8201047647 | 0,8197896349 | 0,0003151298028 | 29,29253855 | 0,00001334030508 |
| AF_1557 | RASGRP1 | 0,007786515805 | 0,006253846927 | 0,001532668879 | 17,53637812 | 0,001753367932 |
| AF_1588 | AQP2 | 0,01382013194 | 0,01262341776 | 0,001196714184 | 16,50612816 | 0,002761966709 |
| AF_1620 | NADSYN1 | 0,02463090053 | 0,02292391302 | 0,001706987508 | 41,37933785 | 0,00000009315064475 |
| AF_1642 | FADS1 | 3,628145165 | 3,625037933 | 0,003107232386 | 10,20621951 | 0,03929380701 |
| AF_1775 | PCCA | 0,7648488704 | 0,7639763486 | 0,000872521782 | 17,93037528 | 0,001491480849 |
| AF_1783 | DOCK9 | 0,01792926443 | 0,01723753967 | 0,0006917247652 | 13,15480668 | 0,01116276863 |
| AF_1787 | IPO5 | 0,1220430759 | 0,1210062034 | 0,001036872438 | 23,41509412 | 0,0001612436106 |
| AF_1799 | CNOT6L | 0,001534889751 | 0,001475056985 | 0,00005983276553 | 12,23167376 | 0,01670106848 |
| AF_1807 | LOC103906701 | 0,366533191 | 0,3612674233 | 0,005265767622 | 13,2091632 | 0,01096226353 |
| AF_1816 | JCHAIN | 10,77552827 | 10,76832797 | 0,007200292626 | 21,97362283 | 0,000279273065 |
| AF_1882 | KIF20A | 0,04869304532 | 0,04764856366 | 0,001044481659 | 42,14103884 | 0,00000007952623791 |
| AF_1887 | TMEM68 | 0,1204860162 | 0,1162487623 | 0,004237253928 | 24,76516954 | 0,00009295190657 |
| AF_1912 | USP53 | 0,006054971834 | 0,005812362898 | 0,000242608936 | 25,82288114 | 0,00005818049088 |
| AF_1917 | PRDM5 | 4,079948322 | 4,076727551 | 0,003220770996 | 14,76791604 | 0,005728264803 |
| AF_1929 | PTCD3 | 0,00921734749 | 0,007922864097 | 0,001294483393 | 14,21802588 | 0,007197335296 |
| AF_1975 | ATXN1L | 0,02137468629 | 0,02136486398 | 0,000009822306982 | 22,58436207 | 0,0002253074507 |
| AF_2053 | ADAD1 | 4,033811871 | 4,030321859 | 0,003490012375 | 29,26333547 | 0,00001337397837 |
| AF_2077 | MYRF | 0,2893620799 | 0,2886663865 | 0,0006956933596 | 22,51520215 | 0,0002308235853 |
| AF_2083 | LMAN1 | 0,00295427087 | 0,002950300764 | 0,000003970105988 | 12,26214011 | 0,01649402119 |
| AF_2109 | PLEKHM2 | 1,269284393 | 1,268165549 | 0,001118843695 | 18,95591141 | 0,0009647814647 |
| AF_2179 | VPS13D | 0,008764636026 | 0,008260184461 | 0,0005044515649 | 11,03963978 | 0,02773965306 |
| AF_2266 | BOC | 0,002312002236 | 0,001873143668 | 0,0004388585682 | 11,62245584 | 0,0216244603 |
| AF_2282 | NME7 | 0,06470575529 | 0,06461877215 | 0,00008698313518 | 12,15978793 | 0,01721832703 |
| AF_2333 | DACT1 | 0,01632246372 | 0,01565816008 | 0,00066430364 | 25,27772448 | 0,00007315760713 |
| AF_2373 | PSMC6 | 1,675165805 | 1,665237367 | 0,009928437524 | 14,66774347 | 0,006003237488 |
| AF_2396 | L2HGDH | 0,007668108554 | 0,005914862833 | 0,001753245721 | 10,50968082 | 0,03483108728 |
| AF_2501 | CBX7 | 0,1420097652 | 0,1419000797 | 0,0001096855037 | 22,35473643 | 0,0002416773208 |
| AF_2518 | SGSM3 | 0,5203066568 | 0,5189417723 | 0,001364884486 | 22,67445553 | 0,0002208660149 |
| AF_2675 | SPG20 | 0,02931811527 | 0,02723157383 | 0,002086541444 | 9,8522667 | 0,04598483916 |
| AF_2773 | TBC1D9 | 0,004914651648 | 0,002685978369 | 0,00222867328 | 16,33249122 | 0,002977310999 |
| AF_2789 | GAB1 | 0,04923557324 | 0,04912591907 | 0,0001096541685 | 27,40643562 | 0,0000306161936 |
| AF_2812 | COCH | 0,01731891179 | 0,01730826488 | 0,00001064690551 | 14,23614743 | 0,007197335296 |
| AF_2852 | BRMS1L | 0,01132246109 | 0,01017352006 | 0,001148941035 | 12,51335502 | 0,01465849019 |
| AF_2853 | MBIP | 0,04193802384 | 0,03967564561 | 0,002262378234 | 15,61848747 | 0,004050264847 |
| AF_2889 | AGL | 0,1418661565 | 0,1390376657 | 0,002828490814 | 66,87906083 | 2,40E-12 |
| AF_2912 | ABCA4 | 0,3394585731 | 0,3384368122 | 0,001021760962 | 30,7158949 | 0,000007145043674 |
| AF_2934 | BRDT | 1,171623807 | 1,170830269 | 0,0007935385179 | 36,82858007 | 0,00000064207998 |
| AF_2954 | JMJD1C | 0,1866937015 | 0,1856161309 | 0,001077570657 | 23,46840075 | 0,0001586972209 |
| AF_3056 | SLC6A17 | 0,2825163628 | 0,2819921229 | 0,0005242398926 | 13,34553852 | 0,01033031961 |
| AF_3175 | COQ9 | 11,28423583 | 11,28178076 | 0,002455069428 | 42,046746 | 0,00000007974715523 |
| AF_3429 | ZCCHC4 | 0,02145941441 | 0,01973715793 | 0,001722256484 | 21,72239629 | 0,0003069250126 |
| AF_3445 | TMA16 | 0,4944993733 | 0,4944324818 | 0,00006689145004 | 41,66894926 | 0,00000009188883773 |
| AF_3475 | GALNT7 | 0,05854879404 | 0,03891157761 | 0,01963721643 | 22,57746357 | 0,0002253074507 |
| AF_3490 | SPCS3 | 0,06083411427 | 0,05559048664 | 0,00524362763 | 13,6583595 | 0,009124118166 |
| AF_3520 | SUSD5 | 0,2260853005 | 0,2255291055 | 0,0005561950056 | 14,18489382 | 0,007288902697 |
| AF_3559 | STARD3NL | 1,781957483 | 1,780455617 | 0,0015018653 | 17,60083887 | 0,001712473949 |
| AF_3752 | HNF4G | 0,07383319862 | 0,07376917012 | 0,00006402850067 | 14,92828196 | 0,005340961046 |
| AF_3784 | NEDD4 | 0,007867344004 | 0,00744343847 | 0,0004239055339 | 29,64929325 | 0,00001143422481 |
| AF_3839 | MAGI3 | 0,01492588857 | 0,0119220488 | 0,003003839769 | 26,5290658 | 0,00004373746908 |
| AF_3995 | ITPR3 | 0,01359517896 | 0,01153521953 | 0,002059959425 | 32,66550056 | 0,000003252830472 |
| AF_4022 | ZNF76 | 2,075821736 | 2,072063911 | 0,003757825333 | 19,81570631 | 0,0006654562324 |
| AF_4157 | FANCA | 0,04163997639 | 0,03987345235 | 0,00176652404 | 12,81989673 | 0,01289845235 |
| AF_4405 | COPB2 | 0,003382067176 | 0,003077928579 | 0,0003041385963 | 11,80625502 | 0,02000600088 |
| AF_4464 | RPAC2 | 0,0635537822 | 0,05574498825 | 0,007808793955 | 13,70892113 | 0,008979953594 |
| AF_4473 | LOC103894128 | 0,01944187994 | 0,01886969321 | 0,0005721867356 | 29,91099906 | 0,00001041890587 |
| AF_4476 | EXOC1 | 0,1065868812 | 0,1057529498 | 0,0008339313523 | 43,77873236 | 0,00000004054858036 |
| AF_4505 | KIAA2022 | 0,03285119131 | 0,03164966737 | 0,001201523936 | 25,68993935 | 0,00006057186228 |
| AF_4510 | NME8 | 0,02258568456 | 0,0219900496 | 0,0005956349615 | 33,36194355 | 0,000002452653903 |
| AF_4525 | DPY19L1 | 0,8182403043 | 0,8165752796 | 0,001665024678 | 18,42291996 | 0,001219095109 |
| AF_4576 | HERC5 | 10,15353877 | 10,15012397 | 0,003414797855 | 15,18142184 | 0,004823372845 |
| AF_4632 | NPNT | 0,915919142 | 0,9144191119 | 0,001500030088 | 13,66937384 | 0,009117862086 |
| AF_4640 | SGMS2 | 0,469739201 | 0,4695225555 | 0,0002166455254 | 26,67318644 | 0,0000419456323 |
| AF_4656 | EGF | 0,003541120473 | 0,00214877122 | 0,001392349253 | 10,25543171 | 0,03870671253 |
| AF_4666 | HGSNAT | 0,7933310732 | 0,7927564759 | 0,0005745972484 | 33,50895364 | 0,000002402727129 |
| AF_4780 | KIF13A | 0,00945767291 | 0,008894608783 | 0,000563064127 | 15,44083728 | 0,004412036259 |
| AF_4800 | JAKMIP1 | 0,006780313096 | 0,005263738593 | 0,001516574503 | 12,62705578 | 0,01396409478 |

| | | | | | | |
|---|---|---|---|---|---|---|
| AF_4804 | EVC2 | 0,009139362228 | 0,008791149845 | 0,0003482123829 | 41,18885575 | 0,00000009459493518 |
| AF_4871 | TP53BP2 | 0,008828493449 | 0,007884811957 | 0,0009436814923 | 14,35638971 | 0,006899963935 |
| AF_4919 | ZNF512B | 0,0494718192 | 0,04486307598 | 0,004608743225 | 14,94127993 | 0,005334210714 |
| AF_4948 | SLC17A9 | 9,592798121 | 9,587249289 | 0,005548831908 | 11,60112729 | 0,02176278925 |
| AF_4967 | OSBPL2 | 0,1569758082 | 0,1560461371 | 0,0009296711032 | 26,49456859 | 0,0000440602787 |
| AF_5022 | HERC1 | 0,005245301286 | 0,004417134627 | 0,0008281666587 | 27,9951148 | 0,00002303997516 |
| AF_5025 | TBC1D2B | 0,004162930193 | 0,00339313016 | 0,0007698000328 | 9,688079163 | 0,04937725887 |
| AF_5042 | RBM19 | 0,01000283162 | 0,00937909807 | 0,0006237335461 | 33,92347968 | 0,000002025680885 |
| AF_5053 | IRAK1BP1 | 0,06337511533 | 0,06334588215 | 0,00002923317857 | 14,2532382 | 0,007197335296 |
| AF_5071 | TFAP2C | 0,7188867334 | 0,7185770624 | 0,0003096710258 | 34,15064323 | 0,000001842483609 |
| AF_5128 | ACOT7 | 29,60381824 | 29,59718598 | 0,0066322606 | 21,13433397 | 0,0003926042443 |
| AF_5155 | PRDM16 | 0,009940803745 | 0,009488432163 | 0,0004523715827 | 37,61013056 | 0,0000004575096084 |
| AF_5226 | ACAP3 | 1,003614538 | 1,001131029 | 0,002483509415 | 12,99795666 | 0,01200732941 |
| AF_5343 | EP400 | 0,1150294599 | 0,1103427452 | 0,004686714648 | 18,10481487 | 0,001403732804 |
| AF_5352 | ZDHHC8 | 0,03830667807 | 0,03705876978 | 0,001247908287 | 33,46172869 | 0,000002416338058 |
| AF_5369 | SEPTIN2 | 1,909258359 | 1,907081141 | 0,002177218284 | 25,88565162 | 0,0000569794185 |
| AF_5373 | UFD1L | 0,02801107365 | 0,02662521742 | 0,001385856228 | 12,0102065 | 0,01825336473 |
| AF_5390 | SCN8A | 0,007425941976 | 0,007375021108 | 0,0000509208685 | 33,20317029 | 0,000002610779329 |
| AF_5464 | CD101 | 0,01898172726 | 0,0167812132 | 0,002200514066 | 13,46780486 | 0,009805205741 |
| AF_5508 | C7 | 0,01332810461 | 0,01144798466 | 0,001880119954 | 22,40326497 | 0,0002374761789 |
| AF_5516 | SELENOP | 0,2466970662 | 0,242742043 | 0,00395502321 | 13,83298998 | 0,008507205667 |
| AF_5623 | IDUA | 0,2397789785 | 0,2378589348 | 0,001920043743 | 57,05977363 | 0,00000000001663416904 |
| AF_5652 | NEK4 | 0,7581716173 | 0,7564679075 | 0,001703709806 | 17,47838987 | 0,001790203317 |
| AF_5751 | MAP4K3 | 14,66609571 | 14,66040348 | 0,005692227596 | 41,17534904 | 0,00000009459493518 |
| AF_5762 | HNRNPLL | 0,007797190854 | 0,007232229832 | 0,000564961022 | 19,21583022 | 0,0008593548994 |
| AF_5846 | GRB14 | 0,01006703456 | 0,009351716978 | 0,0007153175826 | 19,30402232 | 0,0008262364449 |
| AF_5849 | IFIH1 | 1,112654948 | 1,111564765 | 0,001090183169 | 26,36421253 | 0,00004612420485 |
| AF_5866 | MYO10 | 0,02321004545 | 0,02320028941 | 0,000009756041587 | 11,71040062 | 0,02082314261 |
| AF_5911 | STK36 | 0,03436092576 | 0,03374225233 | 0,0006186734243 | 19,93302898 | 0,0006412721739 |
| AF_5943 | SMARCAL1 | 0,5482235134 | 0,5473190464 | 0,0009044670416 | 39,60681089 | 0,0000001821791806 |
| AF_5979 | DIRC2 | 0,01305891456 | 0,01084909637 | 0,002209818196 | 9,799296511 | 0,04689719347 |
| AF_5984 | ADCY5 | 1,103921479 | 1,103588253 | 0,0003332260808 | 74,26338727 | 2,12E-13 |
| AF_5987 | CCDC14 | 0,0305091642 | 0,02925655797 | 0,001252606235 | 26,99928615 | 0,00003694608098 |
| AF_6080 | ARAP2 | 0,05158722253 | 0,05116430874 | 0,000422913793 | 13,86523737 | 0,008425725666 |
| AF_6230 | CUL4B | 0,007016997298 | 0,004677539947 | 0,00233945735 | 11,44550269 | 0,02328629733 |
| AF_6305 | UFL1 | 0,5520319313 | 0,5509267844 | 0,001105146894 | 20,37581776 | 0,0005483757567 |
| AF_6321 | COQ3 | 0,009966581185 | 0,008790617018 | 0,001175964167 | 11,62806416 | 0,0216244603 |
| AF_6396 | MATN2 | 76,93773028 | 76,92720736 | 0,01052292833 | 60,01790401 | 4,54E-11 |
| AF_6428 | PLCG1 | 3,766835923 | 3,763865664 | 0,002970258406 | 28,17008723 | 0,00002182395621 |
| AF_6444 | ATRIP | 0,826562458 | 0,825510247 | 0,001052211022 | 31,0343879 | 0,000006257541396 |
| AF_6455 | NBAS | 0,001412499983 | 0,001080740158 | 0,0003317598248 | 10,96005133 | 0,02844513253 |
| AF_6460 | GREB1 | 0,03241624331 | 0,03175392508 | 0,0006623182301 | 64,38662242 | 6,26E-12 |
| AF_6466 | ROCK2 | 0,008449948599 | 0,007868163878 | 0,0005817847213 | 25,78843661 | 0,00005818049088 |
| AF_6467 | PQLC3 | 40,56335263 | 40,55436553 | 0,008987104862 | 11,18368032 | 0,02608280139 |
| AF_6523 | AKAP11 | 0,00825123253 | 0,007359456234 | 0,0008917762966 | 32,3046771 | 0,000003717759927 |
| AF_6540 | DIAPH3 | 0,03012633944 | 0,02931124288 | 0,0008150965607 | 18,24482378 | 0,001320680642 |
| AF_6604 | MRC1 | 0,00239331999 | 0,001176886811 | 0,001216433179 | 10,32843756 | 0,03775007209 |
| AF_6608 | SLC39A12 | 0,008581028928 | 0,008577155282 | 0,000003873645773 | 16,69277537 | 0,002556544635 |
| AF_6700 | EDEM3 | 0,004357704614 | 0,003198556722 | 0,001159147892 | 12,22295038 | 0,01672638451 |
| AF_6764 | STX6 | 0,124444855 | 0,1216400366 | 0,002804818449 | 21,23120164 | 0,0003762162058 |
| AF_6792 | MAP3K3 | 1,58664796 | 1,584931662 | 0,001716297786 | 22,65415592 | 0,0002215112826 |
| AF_6794 | DLEC1 | 2,110929476 | 2,110066959 | 0,0008625175625 | 45,75506885 | 0,00000002140975404 |
| AF_6831 | WNT3A | 3,94639955 | 3,945296909 | 0,001102640964 | 33,38096408 | 0,000002452653903 |
| AF_6861 | HRAS | 2,322802122 | 2,321018907 | 0,001783214186 | 25,33336037 | 0,0000717847392 |
| AF_6879 | AP2A2 | 0,004726739816 | 0,003995685639 | 0,0007310541775 | 18,04358598 | 0,001440876688 |
| AF_6922 | ATP11A | 0,2737805531 | 0,2726085633 | 0,001171989812 | 14,25117298 | 0,007197335296 |
| AF_6996 | RPL31 | 0,2160326998 | 0,216014005 | 0,00001869487392 | 13,19615707 | 0,01100020325 |
| AF_7017 | GMEB1 | 2,931421898 | 2,929578889 | 0,001843009364 | 15,39300465 | 0,004502227211 |
| AF_7112 | SRSF10 | 1,34082902 | 1,340098115 | 0,0007309049577 | 20,61811772 | 0,0004934399512 |
| AF_7253 | GTF2IRD1 | 0,02196983565 | 0,01865322187 | 0,003316613786 | 24,15078056 | 0,0001187902981 |
| AF_7276 | UNC13C | 0,02223902557 | 0,02183811405 | 0,0004009115199 | 49,26832903 | 0,000000005478789589 |
| AF_7370 | TMC3 | 0,03235078533 | 0,03197082473 | 0,0003799606008 | 17,45212477 | 0,001805484889 |
| AF_7436 | ELP2 | 0,03568769623 | 0,03566942553 | 0,00001827070224 | 21,55770364 | 0,0003252035147 |
| AF_7489 | CUNH18orf63 | 0,01907021904 | 0,01764266773 | 0,00142755131 | 18,01064022 | 0,001458156507 |
| AF_7503 | MYLK3 | 0,05456947156 | 0,05277131118 | 0,00179816038 | 15,93248378 | 0,003554357963 |
| AF_7512 | PHKB | 0,01341722578 | 0,01100354306 | 0,002413682718 | 19,41628267 | 0,000792875358 |
| AF_7564 | SLC7A6 | 0,2615421031 | 0,2613760966 | 0,0001660065391 | 29,83809698 | 0,00001066821507 |
| AF_7642 | IARS2 | 2,015634505 | 2,014124206 | 0,001510298873 | 19,93224513 | 0,0006412721739 |
| AF_7649 | ASB9 | 1,937031878 | 1,936164617 | 0,0008672608589 | 28,00820986 | 0,00002303997516 |
| AF_7678 | CDKL5 | 0,005686387081 | 0,002764532136 | 0,002921854945 | 14,12189587 | 0,007453344956 |
| AF_7709 | SMS | 0,01585477328 | 0,01276152485 | 0,003093248426 | 21,31755911 | 0,0003624335635 |
| AF_7711 | PTCHD1 | 0,274252728 | 0,2734066306 | 0,000846097412 | 10,32456479 | 0,03775007209 |
| AF_7751 | EXPH5 | 0,006877410336 | 0,004919859205 | 0,001957551131 | 10,98527823 | 0,02830494523 |
| AF_7762 | MARCH6 | 1,196042983 | 1,194843402 | 0,001199580813 | 12,94650879 | 0,0122108129 |
| AF_7895 | GLS | 0,0153220458 | 0,01531663302 | 0,000005412780182 | 29,30605089 | 0,00001334030508 |
| AF_7908 | ATP6V1C1 | 0,0673397153 | 0,06350905571 | 0,003830659589 | 32,54548799 | 0,000003399662175 |
| AF_8088 | GAN | 0,08461075488 | 0,08273966745 | 0,001871087426 | 15,37909205 | 0,004516559759 |
| AF_8090 | PKD1L2 | 0,4965526433 | 0,4950327396 | 0,001519903617 | 22,01723792 | 0,0002750727968 |
| AF_8127 | SLC25A4 | 0,07242602985 | 0,06855132334 | 0,003874706503 | 30,76283202 | 0,00000707319326 |
| AF_8156 | ASAH1 | 0,180978473 | 0,1790564627 | 0,001922010334 | 56,10039996 | 0,0000000000230379858 |
| AF_8173 | SLC7A2 | 0,2158252023 | 0,2057370341 | 0,01008816826 | 35,56437263 | 0,0000010038002 |
| AF_8191 | KIAA1211 | 1,347269666 | 1,345599928 | 0,00166973789 | 18,15845725 | 0,001372802496 |
| AF_8264 | MYBPC3 | 0,002465328147 | 0,001323866208 | 0,001141461939 | 10,81497555 | 0,03037211338 |
| AF_8271 | ARFGAP2 | 1,215035571 | 1,212752196 | 0,002283375027 | 13,64285207 | 0,009151103905 |
| AF_8290 | PEX16 | 0,02533374716 | 0,0228588797 | 0,002474867463 | 13,70609651 | 0,008979953594 |
| AF_8317 | ITPKA | 0,07948197886 | 0,07942590878 | 0,00005607008548 | 19,80603098 | 0,0006654562324 |
| AF_8333 | PLA2G4E | 1,045559116 | 1,044903361 | 0,0006557548897 | 26,80361516 | 0,00003969861473 |
| AF_8336 | AOAH | 1,004717416 | 1,003816178 | 0,0009012381161 | 17,93449351 | 0,001491480849 |
| AF_8341 | CAPN3 | 1,393148687 | 1,391217407 | 0,001931279959 | 26,55568843 | 0,00004359321896 |
| AF_8360 | PTGR2 | 6,114255848 | 6,111060549 | 0,003195298252 | 12,39452189 | 0,01551853998 |
| AF_8401 | SEC24C | 0,08123627963 | 0,08067922173 | 0,000557907892 | 53,03899402 | 0,00000000009332595946 |
| AF_8439 | NOLC1 | 0,2877836602 | 0,2870802633 | 0,0007033969589 | 15,31851306 | 0,004590771994 |
| AF_8507 | IFIT5 | 0,8703109592 | 0,8688983555 | 0,001412603683 | 17,81251205 | 0,001553340243 |
| AF_8543 | IDE | 0,06954494136 | 0,0652883135 | 0,004256627859 | 40,78200642 | 0,0000001114783137 |
| AF_8564 | ENTPD1 | 0,9232658192 | 0,9226229107 | 0,0006429084951 | 44,69119138 | 0,00000003252280495 |
| AF_8615 | SLC2A1 | 1,46749876 | 1,466357737 | 0,001141023196 | 12,12370595 | 0,01748439292 |
| AF_8676 | SMC3 | 0,001555045025 | 0,001225269526 | 0,0003297754991 | 11,3339364 | 0,02450283608 |
| AF_8679 | SHOC2 | 0,006466755992 | 0,006047314469 | 0,0004194415235 | 13,45599039 | 0,009832643163 |
| AF_8703 | TDRD1 | 0,02383866312 | 0,02297535507 | 0,0008633080505 | 16,1104096 | 0,003275988575 |
| AF_8721 | CCDC172 | 0,02139381718 | 0,02138911033 | 0,000004706850261 | 20,22886338 | 0,000577192499 |

| | | | | | | |
|---|---|---|---|---|---|---|
| AF_8749 | MCMBP | 0,01768397501 | 0,01629924446 | 0,001384730553 | 24,41705036 | 0,0001087843763 |
| AF_8859 | KIAA0100 | 0,5151053434 | 0,512962425 | 0,002142918406 | 15,74542065 | 0,0038143613 |
| AF_8931 | ULK2 | 0,4685110285 | 0,466283928 | 0,002227100427 | 11,27028372 | 0,02516972731 |
| AF_8948 | RAD51C | 0,2599204861 | 0,2591794804 | 0,0007410057144 | 17,04494169 | 0,002159639103 |
| AF_8972 | INTS2 | 0,02673584555 | 0,02648060925 | 0,0002552363015 | 44,42907002 | 0,0000000347669685 |
| AF_8983 | APPBP2 | 0,009512621027 | 0,008953854224 | 0,0005587668028 | 28,01091184 | 0,00002303997516 |
| AF_9000 | SYNRG | 0,001817541614 | 0,001204579517 | 0,0006129620973 | 11,12583421 | 0,0267139225 |
| AF_9101 | LRPPRC | 0,01882777214 | 0,01808437273 | 0,0007433994089 | 32,83424921 | 0,000003038031195 |
| AF_9132 | HDHD2 | 0,04337220122 | 0,04336310223 | 0,000009098987899 | 23,64911572 | 0,0001484958218 |
| AF_9208 | FRMPD4 | 0,003030208313 | 0,002473209253 | 0,0005569990596 | 12,1091944 | 0,01756255054 |
| AF_9256 | TLK1 | 0,002431916282 | 0,001967820711 | 0,0004640955707 | 10,36047204 | 0,03735043747 |
| AF_9261 | DYNC1I2 | 1,176785899 | 1,175581336 | 0,001204562944 | 31,45697981 | 0,000005205579133 |
| AF_9269 | PPP1R9A | 0,01063592033 | 0,009171691092 | 0,001464229241 | 12,685698 | 0,01367824218 |
| AF_9324 | NRP1 | 0,1333378255 | 0,1329186701 | 0,0004191554272 | 15,36390424 | 0,004527384238 |
| AF_9395 | LEPR | 0,01815337243 | 0,01753063618 | 0,0006227362465 | 17,30625269 | 0,001934517579 |
| AF_9464 | NRG3 | 0,1428630036 | 0,1375500413 | 0,005312962331 | 21,96118644 | 0,000279273065 |
| AF_9465 | SH2D4B | 0,9731184149 | 0,9724262155 | 0,0006921994229 | 34,19633193 | 0,000001836086873 |
| AF_9481 | NRG4 | 0,1593165742 | 0,159262557 | 0,00005401721511 | 10,07382642 | 0,04170853662 |
| AF_9570 | GLIPR1 | 5,004217405 | 5,002901418 | 0,001315987432 | 14,55531131 | 0,006330469463 |
| AF_9601 | CACNA1C | 0,06601467318 | 0,06266040964 | 0,003354263541 | 28,91877129 | 0,00001570693992 |
| AF_9671 | CMAS | 0,8330908392 | 0,8321350861 | 0,0009557530784 | 24,1412006 | 0,0001187902981 |
| AF_9733 | RAB28 | 0,02591849541 | 0,02391896919 | 0,001999526222 | 16,38930276 | 0,002917141322 |
| AF_9766 | NCAPG | 0,003322747161 | 0,002163973817 | 0,001158773344 | 10,38238793 | 0,03703197152 |
| AF_9868 | KIAA1468 | 0,3420213356 | 0,3409977429 | 0,001023592702 | 10,52841969 | 0,03465890023 |
| AF_9957 | RASSF6 | 4,081119081 | 4,079204661 | 0,00191442074 | 18,80027671 | 0,001033089575 |
| AF_9961 | ANKRD17 | 0,3210492809 | 0,3201732193 | 0,000876061543 | 22,69096804 | 0,0002206534446 |
| AF_9984 | XPO4 | 0,006674037121 | 0,005944214685 | 0,0007298224363 | 21,91195547 | 0,0002816939081 |
| AF_10151 | CTNNBL1 | 0,5995043483 | 0,5982384568 | 0,001265891573 | 17,88148044 | 0,001515023888 |
| AF_10182 | PROX1 | 0,3310229367 | 0,3305490473 | 0,0004738893888 | 34,93819995 | 0,000001291340669 |
| AF_10189 | FLVCR1 | 1,160623638 | 1,159496641 | 0,001126996925 | 20,18185253 | 0,0005877013744 |
| AF_10200 | INTS7 | 0,005519745352 | 0,003355153893 | 0,002164591459 | 13,53318841 | 0,009547827175 |
| AF_10205 | TRAF5 | 0,9876421331 | 0,9852680321 | 0,002374100997 | 21,63483425 | 0,0003147808264 |
| AF_10208 | HHAT | 0,07034864676 | 0,06917693409 | 0,001171712673 | 25,80326173 | 0,00005818049088 |
| AF_10241 | HN1 | 0,2806232341 | 0,2805158291 | 0,0001074049677 | 11,55207272 | 0,02224746507 |
| AF_10274 | MYCBPAP | 0,4444195016 | 0,4434113147 | 0,001008186909 | 17,97118255 | 0,001480514134 |
| AF_10344 | HOMER3 | 1,606971453 | 1,605087682 | 0,001883771142 | 20,01990655 | 0,0006236505749 |
| AF_10462 | NCLN | 1,50075604 | 1,49811764 | 0,00263839976 | 9,826268444 | 0,04648243294 |
| AF_10487 | FCHO2 | 0,03962909106 | 0,03745220037 | 0,002176890685 | 28,82779723 | 0,00001624661897 |
| AF_10514 | PCDH11X | 0,005605937955 | 0,005599075267 | 0,000006862688305 | 40,51142824 | 0,0000001199216584 |
| AF_10519 | SMARCB1 | 0,1795297329 | 0,1775822607 | 0,001947472196 | 22,22507981 | 0,0002544694618 |
| AF_10646 | RALGPS1 | 0,1207896812 | 0,1171060447 | 0,003683636471 | 11,25503775 | 0,02529419107 |
| AF_10666 | GUF1 | 0,02711422799 | 0,02040706234 | 0,006707165651 | 15,23696085 | 0,004735497959 |
| AF_10728 | MYOCD | 1,392528108 | 1,390967346 | 0,001560762587 | 9,799544073 | 0,04689719347 |
| AF_10832 | POC5 | 4,685859869 | 4,684948015 | 0,000911853748 | 23,64804072 | 0,0001484958218 |
| AF_10875 | FAM189A1 | 0,00678089685 | 0,006010508014 | 0,0007703888352 | 15,9491091 | 0,003538791404 |
| AF_10879 | TARS3 | 1,685980369 | 1,684493039 | 0,00148733002 | 15,88845053 | 0,003605374903 |
| AF_10888 | MYO1E | 0,3490732525 | 0,3480539258 | 0,001019326722 | 19,9444929 | 0,0006412721739 |
| AF_10980 | IQCH | 0,06409245137 | 0,06260364698 | 0,001488804388 | 35,62799999 | 0,0000009934392667 |
| AF_11037 | OVOSTATIN | 0,01068600381 | 0,00978594134 | 0,0009000624739 | 24,98791252 | 0,00008385862313 |
| AF_11098 | CRB2 | 0,05078646755 | 0,04743731574 | 0,003349151809 | 44,08538814 | 0,00000003670802221 |
| AF_11099 | STRBP | 0,005906829765 | 0,004926662138 | 0,0009801676271 | 13,33619199 | 0,01034666184 |
| AF_11108 | ZBTB26 | 0,005145463897 | 0,005137646074 | 0,000007817822596 | 15,14047698 | 0,004869908448 |
| AF_11206 | COQ6 | 0,03072372939 | 0,02753137104 | 0,003192358352 | 17,29328301 | 0,001938818244 |
| AF_11306 | SPATA7 | 0,154469859 | 0,1514558425 | 0,003014016512 | 22,92118804 | 0,0001995063419 |
| AF_11340 | FBLN5 | 0,01399570927 | 0,01214473967 | 0,001850969608 | 10,2330742 | 0,03893079788 |
| AF_11365 | SPI | 0,01256683467 | 0,01076210572 | 0,001804728947 | 13,0311361 | 0,01184437918 |
| AF_11394 | PKDCC | 0,06646315002 | 0,06639261544 | 0,00007053458193 | 10,23745471 | 0,03893079788 |
| AF_11417 | DYNC1H1 | 0,04078894646 | 0,04020711951 | 0,0005818269522 | 20,22921994 | 0,000577192499 |
| AF_11476 | CEP170B | 0,01953696314 | 0,01848662137 | 0,001050341773 | 39,5591675 | 0,0000001821791806 |
| AF_11477 | PLD4 | 1,975296584 | 1,973675324 | 0,001621260392 | 29,66419432 | 0,00001143422481 |
| AF_11502 | ACOT12 | 0,9144216886 | 0,9136783643 | 0,0007433242721 | 20,79618341 | 0,0004570611158 |
| AF_11533 | ANKRD27 | 0,3692949929 | 0,368360055 | 0,0009349379122 | 10,00309342 | 0,04291507833 |
| AF_11577 | TBK1 | 1,779249672 | 1,777257898 | 0,001991774174 | 23,40579309 | 0,0001612436106 |
| AF_11606 | ACSBG2 | 1,029856358 | 1,02816679 | 0,001689567121 | 17,83250184 | 0,001544704043 |
| AF_11735 | UBASH3B | 0,001752235249 | 0,001648236651 | 0,0001039985983 | 10,2294175 | 0,03893079788 |
| AF_11796 | CYFIP1 | 0,272226405 | 0,2718186587 | 0,0004077463288 | 32,91868003 | 0,000002960691945 |
| AF_11804 | PLCXD1 | 0,2166149969 | 0,2111653288 | 0,005449668119 | 14,47459566 | 0,006546817455 |
| AF_11820 | ASMT | 9,637590343 | 9,633554807 | 0,004035535762 | 31,52550751 | 0,000005101655625 |
| AF_11822 | ZBED1 | 0,006095513195 | 0,005113829529 | 0,0009816836655 | 14,22564464 | 0,007197335296 |
| AF_11851 | PLEKHG1 | 0,01280637439 | 0,01147481866 | 0,001331555722 | 17,88003358 | 0,001515023888 |
| AF_11892 | SERAC1 | 0,01682154245 | 0,01633024826 | 0,0004912941936 | 31,85036482 | 0,00000443464767 |
| AF_11938 | MAP3K5 | 0,3513339573 | 0,3502085801 | 0,001125377227 | 19,61666904 | 0,0007243276299 |
| AF_11986 | ENPP3 | 0,6782808561 | 0,6778047243 | 0,00047613181 | 47,00232291 | 0,00000001394050434 |
| AF_12020 | OTOA | 0,1541432555 | 0,1535220303 | 0,0006212251944 | 15,3592274 | 0,004527384238 |
| AF_12034 | CUNH16orf52 | 0,1305708194 | 0,1305001146 | 0,00007070471932 | 19,05284497 | 0,0009234621018 |
| AF_12116 | CLEC16A | 0,00499552975 | 0,004265390717 | 0,0007301390327 | 16,35552869 | 0,002955017286 |
| AF_12130 | DIS3 | 0,002423790662 | 0,002422649933 | 0,000001140729739 | 11,42668662 | 0,02344502645 |
| AF_12166 | KIF25 | 0,021343334 | 0,02072893779 | 0,0006143962098 | 24,22001669 | 0,0001170622884 |
| AF_12181 | ERMARD | 0,08453182397 | 0,08330574262 | 0,001226081349 | 21,02048312 | 0,0004132307583 |
| AF_12214 | SIPA1L2 | 0,001706956146 | 0,001093460181 | 0,0006134959649 | 11,11927941 | 0,0267317854 |
| AF_12236 | LGALS8 | 5,318558341 | 5,317586561 | 0,0009717796977 | 38,27099521 | 0,0000003376870397 |
| AF_12263 | DENND2D | 3,05952287 | 3,05572448 | 0,003798389525 | 13,34794691 | 0,01033031961 |
| AF_12309 | IPO9 | 0,8042759515 | 0,8031402014 | 0,00113575007 | 30,07490732 | 0,000009723234942 |
| AF_12310 | KDM5B | 0,3279725277 | 0,3273010628 | 0,000671464863 | 42,6317459 | 0,00000006841784304 |
| AF_12349 | RNGTT | 0,006748648503 | 0,006740489868 | 0,000008158634841 | 13,56940592 | 0,009405106785 |
| AF_12459 | GPNMB | 0,2668340778 | 0,2575612884 | 0,009272789413 | 16,57799037 | 0,002674918073 |
| AF_12465 | RAPGEF5 | 0,01889986361 | 0,01839942191 | 0,0005004416999 | 26,98885371 | 0,00003694608098 |
| AF_12469 | ABCB5 | 0,003787852021 | 0,003657991636 | 0,0001298603848 | 20,07487745 | 0,0006133313711 |
| AF_12473 | TWISTNB | 0,008063425918 | 0,008059197212 | 0,000004228706331 | 10,91468463 | 0,02902933432 |
| AF_12524 | ATP11C | 0,0118689251 | 0,01061775949 | 0,001251165609 | 10,31173321 | 0,03790123428 |
| AF_12614 | NCOA3 | 1,044490885 | 1,043192859 | 0,001298026163 | 48,44038076 | 0,000000007463786289 |
| AF_12625 | SLC25A3 | 364,6342247 | 364,6013479 | 0,0328767763 | 15,28595127 | 0,004644702801 |
| AF_12725 | OPA1 | 0,3024994997 | 0,3016890638 | 0,0008104358947 | 9,769219148 | 0,04750345054 |
| AF_12757 | TFRC | 0,005932000962 | 0,003732076954 | 0,002199924008 | 12,66167202 | 0,01376304684 |
| AF_12764 | LRCH3 | 0,7834875496 | 0,7814672727 | 0,002020276954 | 22,1414702 | 0,0002618633167 |
| AF_12765 | IQCG | 0,5251012186 | 0,5242630146 | 0,0008382040036 | 23,16964402 | 0,0001801480674 |
| AF_12795 | ABCC5 | 0,001988698841 | 0,001508580674 | 0,0004801181673 | 11,8363059 | 0,01981558631 |
| AF_12855 | FNDC1 | 0,1200977544 | 0,1194923257 | 0,0006054286713 | 23,88316416 | 0,0001341136431 |
| AF_12857 | FANCL | 0,01104571877 | 0,01103928792 | 0,000006430845254 | 21,69188788 | 0,0003097148811 |
| AF_12861 | PUS10 | 0,1353225192 | 0,1340656052 | 0,001256914046 | 22,19162487 | 0,0002570543295 |

| | | | | | | |
|---|---|---|---|---|---|---|
| AF_12863 | KIAA1841 | 0,01704920592 | 0,01388639616 | 0,003162809757 | 18,32559323 | 0,001274147333 |
| AF_13012 | SLC9A8 | 0,05279039255 | 0,05057658751 | 0,002213805035 | 20,6539129 | 0,000488043873 |
| AF_13038 | MEST | 0,01860537543 | 0,01498521545 | 0,003620159975 | 10,77392452 | 0,03092911775 |
| AF_13198 | NAE1 | 0,03636405251 | 0,0312353947 | 0,005128657805 | 17,10002466 | 0,002109537088 |
| AF_13199 | CCDC79 | 0,01323731664 | 0,009782123075 | 0,003455193565 | 15,14098517 | 0,004869908448 |
| AF_13214 | TRAT1 | 2,851200333 | 2,849313836 | 0,001886497255 | 18,59581788 | 0,001128376368 |
| AF_13261 | MAP3K7CL | 3,318271913 | 3,316266302 | 0,002005610787 | 20,60886772 | 0,0004934399512 |
| AF_13270 | TIAM1 | 0,01886357836 | 0,01845503992 | 0,0004085384402 | 16,61127788 | 0,002652423538 |
| AF_13275 | URB1 | 0,01896577496 | 0,01822561061 | 0,0007401643585 | 26,39976405 | 0,00004575316475 |
| AF_13392 | EFHC2 | 1,91030806 | 1,908301679 | 0,002006380286 | 20,03440441 | 0,0006225083081 |
| AF_13473 | CRB1 | 0,1598558885 | 0,1572670449 | 0,002588843597 | 18,63820068 | 0,001109931475 |
| AF_13632 | DCHS2 | 0,01157540067 | 0,01157276165 | 0,000002639015996 | 11,51891867 | 0,02249999658 |
| AF_13648 | GLRB | 0,01202984642 | 0,01056643496 | 0,001463411457 | 12,88656039 | 0,01254748328 |
| AF_13668 | PDZRN3 | 0,008537851061 | 0,008534366134 | 0,000003484926135 | 28,76720785 | 0,00001636278065 |
| AF_13769 | RGS9 | 0,006021303633 | 0,004998317552 | 0,00102298608 | 16,15572942 | 0,00322796075 |
| AF_13777 | ABCA10 | 1,13778764 | 1,136149473 | 0,001638166458 | 35,36509908 | 0,00000106393119 |
| AF_13824 | PPARA | 0,01131840318 | 0,01078165592 | 0,0005367472563 | 14,17951193 | 0,007288902697 |
| AF_13926 | GPR176 | 0,1215294414 | 0,1214882902 | 0,00004115122313 | 24,33714381 | 0,0001122670847 |
| AF_14001 | SLC4A1AP | 6,570386418 | 6,567875788 | 0,002510630009 | 20,40355977 | 0,0005438954367 |
| AF_14010 | KCNK16 | 1,538622147 | 1,537485662 | 0,001136485816 | 17,64933502 | 0,001678651887 |
| AF_14025 | SLC35B2 | 0,0061324904 | 0,005356154334 | 0,0007763360664 | 10,08545863 | 0,04156478997 |
| AF_14042 | EPHX1 | 0,0521990067 | 0,04907549785 | 0,003123508855 | 11,7272589 | 0,02070406814 |
| AF_14101 | UBE3B | 2,473793829 | 2,472426632 | 0,001367197479 | 26,89042217 | 0,00003840729621 |
| AF_14105 | FOXN4 | 2,251175117 | 2,249105152 | 0,002069964979 | 26,59146545 | 0,00004325480657 |
| AF_14110 | SVOP | 1,260238329 | 1,258970696 | 0,001267633098 | 35,84345646 | 0,0000009115943046 |
| AF_14164 | KNTC1 | 0,005241578998 | 0,004435929073 | 0,0008056499247 | 14,93795178 | 0,005334210714 |
| AF_14165 | RSRC2 | 0,0115592052 | 0,01072946168 | 0,0008297435187 | 15,7489292 | 0,0038143613 |
| AF_14199 | P2RX4 | 6,142745091 | 6,140162724 | 0,00258236709 | 26,24005587 | 0,00004861612489 |
| AF_14240 | DNAH10 | 1,213024613 | 1,211230778 | 0,001793835501 | 14,8797613 | 0,00545342123 |
| AF_14293 | SIN3B | 0,02822956777 | 0,02698157918 | 0,001247988595 | 36,62686662 | 0,0000006853780293 |
| AF_14300 | CHERP | 0,01688180952 | 0,01645649813 | 0,0004253113872 | 12,67719549 | 0,01369625122 |
| AF_14331 | CHAF1A | 0,004575334716 | 0,003729171772 | 0,0008461629439 | 12,96404653 | 0,01214623461 |
| AF_14343 | CEP192 | 6,154238662 | 6,149546456 | 0,004692206798 | 18,74885738 | 0,001054987138 |
| AF_14621 | LOC103903316 | 0,003998493062 | 0,003371692487 | 0,0006268005754 | 10,75989083 | 0,03106953792 |
| AF_14622 | valA | 0,275089235 | 0,2747380807 | 0,0003511543192 | 20,10501314 | 0,0006074542916 |
| AF_14728 | CHCHD6 | 14,11627194 | 14,10496955 | 0,01130238587 | 35,99578438 | 0,0000008638019293 |
| AF_14772 | ARF4 | 0,02031024575 | 0,02029828522 | 0,00001196053032 | 10,12302513 | 0,04088209977 |
| AF_14783 | FAM208A | 0,09765536521 | 0,09743287823 | 0,0002224869763 | 12,70514717 | 0,01358518138 |
| AF_14795 | CACNA1D | 2,014551314 | 2,012220591 | 0,002330722913 | 41,34381426 | 0,00000009315064475 |
| AF_14803 | RHOA | 0,0905631409 | 0,09048798405 | 0,00007515684962 | 10,63463008 | 0,03293118247 |
| AF_14805 | CBFA2T2 | 0,02738147276 | 0,02314399054 | 0,004237482218 | 14,16621036 | 0,007313277971 |
| AF_14808 | EDEM2 | 0,9952138131 | 0,993207423 | 0,002006390082 | 11,23011994 | 0,0255461708 |
| AF_14877 | SUMO1 | 0,4851387184 | 0,4850287723 | 0,0001099460721 | 12,05013368 | 0,01793999278 |
| AF_14884 | TRAK2 | 0,002509145438 | 0,001810790373 | 0,0006983550655 | 11,03558264 | 0,02773965306 |
| AF_14894 | FAM126B | 0,008003581223 | 0,007482301797 | 0,0005212794258 | 13,87806384 | 0,008399359311 |
| AF_14895 | ORC2 | 0,0268451121 | 0,02683086393 | 0,00001424816832 | 41,51900488 | 0,00000009188883773 |
| AF_14923 | GTF3C3 | 0,2500293402 | 0,2486308942 | 0,001398445982 | 17,13629781 | 0,002080219925 |
| AF_14936 | INPP5B | 0,4827641573 | 0,4816089587 | 0,001155198563 | 22,60934945 | 0,000224920156 |
| AF_14953 | STK40 | 0,005088646176 | 0,00434126912 | 0,0007473770558 | 10,72968295 | 0,03146691585 |
| AF_14968 | AGO3 | 0,01866136335 | 0,01754160411 | 0,001119759231 | 11,16372981 | 0,02627703608 |
| AF_15036 | YARS | 0,009833433495 | 0,007559311133 | 0,002274122362 | 11,52724447 | 0,02246667711 |
| AF_15280 | TRPM8 | 0,2532695124 | 0,2523353915 | 0,0009341209363 | 24,31741812 | 0,0001124229825 |
| AF_15307 | MREG | 0,1070936174 | 0,1070283721 | 0,0000652453506 | 9,931593102 | 0,04428207066 |
| AF_15354 | AGXT2 | 0,01022199828 | 0,009563731156 | 0,0006582671236 | 14,21997271 | 0,007197335296 |
| AF_15375 | PDZD2 | 0,0779070751 | 0,0775053774 | 0,0004016977047 | 22,95031747 | 0,0001984538852 |
| AF_15382 | APH1A | 0,0982923112 | 0,09658492472 | 0,001707386476 | 17,54142625 | 0,001753367932 |
| AF_15421 | TLE4 | 0,2576819117 | 0,2572341749 | 0,0004477367506 | 10,33330952 | 0,03775007209 |
| AF_15426 | VPS13A | 0,03807496994 | 0,03713535852 | 0,0009396114258 | 45,66387865 | 0,00000002140975404 |
| AF_15504 | PACS2 | 0,1367160786 | 0,1363119338 | 0,0004041448222 | 26,16736629 | 0,00004994132816 |
| AF_15518 | ACOX3 | 0,02429256467 | 0,02428787386 | 0,000004690809109 | 20,31203329 | 0,0005630121111 |
| AF_15608 | USP6NL | 0,00916706987 | 0,006771909296 | 0,002395160573 | 13,57588248 | 0,009404508786 |
| AF_15646 | HSPA14 | 0,05924882961 | 0,05635405767 | 0,002894771942 | 10,98829738 | 0,02830494523 |
| AF_15710 | FAM179A | 0,01538644056 | 0,01437337311 | 0,001013067449 | 21,90581397 | 0,0002816939081 |
| AF_15763 | DSC1 | 0,005581450024 | 0,005180416633 | 0,0004010333913 | 12,52808445 | 0,01459267036 |
| AF_15828 | WWC1 | 0,03445973961 | 0,03373306233 | 0,0007266772826 | 23,59158114 | 0,0001510705047 |
| AF_15869 | AFAP1L1 | 0,9440083286 | 0,9427131802 | 0,00129514836 | 22,45328945 | 0,000233533378 |
| AF_15924 | UIMC1 | 0,02073651934 | 0,01956924046 | 0,001167278878 | 12,87378737 | 0,01259051098 |
| AF_16096 | SPOCK1 | 0,09207915351 | 0,08871949116 | 0,003359662355 | 14,34198351 | 0,006926075861 |
| AF_16232 | RAC1 | 0,012486128 | 0,01181707786 | 0,0006690501391 | 22,9466486 | 0,0001984538852 |
| AF_16233 | AQP11 | 8,697188071 | 8,694510574 | 0,002677496624 | 20,85154359 | 0,0004471587929 |
| AF_16351 | FOLH1B | 0,03287269411 | 0,03195599949 | 0,0009166946186 | 21,63543681 | 0,0003147808264 |
| AF_16447 | EIF2AK1 | 0,03051825929 | 0,03045053435 | 0,00006772494567 | 13,58446374 | 0,009394039244 |
| AF_16500 | GRIFIN | 0,0458192666 | 0,04581227938 | 0,000006987224221 | 19,18781848 | 0,0008672000429 |
| AF_16502 | TTYH3 | 0,09239911559 | 0,09231317363 | 0,00008594196221 | 42,3911723 | 0,00000007350331379 |
| AF_16567 | TOM1L2 | 0,4674341354 | 0,4667503793 | 0,0006837560419 | 10,004131 | 0,04291507833 |
| AF_16568 | DRC3 | 0,4150412829 | 0,4143344688 | 0,0007068141427 | 15,90768 | 0,003584771225 |
| AF_16581 | FAM83G | 1,306908631 | 1,30438457 | 0,002524060496 | 41,57315335 | 0,00000009188883773 |
| AF_16751 | RASA2 | 0,01030588393 | 0,009700126009 | 0,0006057579245 | 12,10183038 | 0,01757770101 |
| AF_16756 | GK5 | 0,6507002745 | 0,6498188393 | 0,0008814352149 | 11,61919467 | 0,0216244603 |
| AF_16776 | GRIK4 | 2,669365072 | 2,667329241 | 0,002035830826 | 36,59851399 | 0,0000006853780293 |
| AF_16779 | TMEM136 | 0,02006504086 | 0,01877310066 | 0,001291940198 | 15,79180384 | 0,003755474746 |
| AF_16783 | TRIM29 | 0,02194445052 | 0,02065662214 | 0,001287828375 | 17,92918951 | 0,001491480849 |
| AF_16790 | FGD6 | 0,01902910961 | 0,01661895308 | 0,002410156534 | 19,54338983 | 0,0007476379241 |
| AF_16799 | LTA4H | 0,02098161647 | 0,01911363397 | 0,001867982502 | 31,83374595 | 0,000004434647767 |
| AF_16828 | LMOD2 | 3,78860204 | 3,785384772 | 0,003217267648 | 18,54218591 | 0,001153658776 |
| AF_16835 | IQUB | 0,3910552716 | 0,3903328286 | 0,0007224430584 | 19,87043829 | 0,0006544956179 |
| AF_16836 | ATP6AP1 | 0,147821687 | 0,1455122342 | 0,002309452808 | 35,50898966 | 0,000001010545493 |
| AF_16867 | MDFIC | 26,60651394 | 26,60174685 | 0,004767087208 | 18,89987147 | 0,0009874728638 |
| AF_16937 | PKN3 | 0,00501285254 | 0,003824853164 | 0,001187999376 | 11,91091436 | 0,01913714523 |
| AF_16944 | KYAT1 | 0,09659550055 | 0,09074181815 | 0,005853682406 | 31,06970091 | 0,000006233130315 |
| AF_17026 | ADAMTS13 | 11,86514811 | 11,86122736 | 0,003920746695 | 37,10190558 | 0,0000005746430133 |
| AF_17113 | IFFO1 | 0,02146014097 | 0,02145037573 | 0,000009765239648 | 11,73360023 | 0,02069539674 |
| AF_17152 | ZYX | 1,120612987 | 1,119336387 | 0,001276600268 | 21,49276918 | 0,0003339329224 |
| AF_17339 | STIL | 0,09317566221 | 0,09310628826 | 0,00006937395353 | 9,952216005 | 0,04392789491 |
| AF_17367 | CC2D1B | 0,005296772346 | 0,004446239018 | 0,0008505333279 | 13,85139554 | 0,008456593404 |
| AF_17377 | ZYG11B | 0,01372164763 | 0,01282870213 | 0,0008929454923 | 17,47765812 | 0,001790203317 |
| AF_17412 | USP24 | 0,06424886691 | 0,06366709996 | 0,0005817669493 | 12,96304662 | 0,01214623461 |
| AF_17441 | ALG6 | 0,05709463351 | 0,05663136903 | 0,0004632644736 | 36,45413677 | 0,0000007192508452 |
| AF_17459 | SUDS3 | 70,38629047 | 70,37249321 | 0,01379725802 | 46,06709203 | 0,0000000002041132013 |
| AF_17490 | TM9SF2 | 0,008754830559 | 0,006922652649 | 0,001832177911 | 16,59378138 | 0,002664766814 |

| | | | | | | |
|---|---|---|---|---|---|---|
| AF_17538 | FAM162B | 0,6810530544 | 0,6746913071 | 0,006361747253 | 15,07192491 | 0,005022567467 |
| AF_17593 | STXBP4 | 0,007275735737 | 0,005531611466 | 0,001744124272 | 12,09287541 | 0,01760713523 |
| AF_17641 | MILR1 | 0,01563041921 | 0,01426280171 | 0,001367617498 | 11,82657272 | 0,01985738016 |
| AF_17652 | PSMD12 | 0,01214793827 | 0,01024474829 | 0,001903189987 | 16,25940115 | 0,003076318981 |
| AF_17653 | HELZ | 0,2350236519 | 0,2343826218 | 0,0006410300136 | 23,58216135 | 0,0001510705047 |
| AF_17826 | LIMD2 | 16,29431379 | 16,29079534 | 0,0035184548 | 21,92703059 | 0,0002816939081 |
| AF_17836 | CDC27 | 0,03033322529 | 0,02892258853 | 0,001410636764 | 16,78791126 | 0,002447333112 |
| AF_17906 | ARR3 | 0,06929399564 | 0,06736850641 | 0,001925489224 | 10,50859622 | 0,03483108728 |
| AF_17913 | FNDC3A | 0,01323979093 | 0,01203938489 | 0,001200406048 | 14,86787641 | 0,005466557203 |
| AF_17945 | POF1B | 0,003764213756 | 0,003762620378 | 0,000001593377149 | 10,25951784 | 0,03870671253 |
| AF_17947 | FNDC3B | 0,004914860153 | 0,004024141611 | 0,0008907185417 | 12,71984003 | 0,01352527589 |
| AF_18159 | ARSK | 0,008774920918 | 0,008168778641 | 0,0006061422771 | 13,6551254 | 0,009124118166 |
| AF_18225 | WEE2 | 0,02691658058 | 0,02277707934 | 0,004139501242 | 19,8839262 | 0,0006535336967 |
| AF_18243 | SVOPL | 0,1007933191 | 0,09694915952 | 0,003844159566 | 19,84147914 | 0,0006605917169 |
| AF_18269 | TH | 0,009722248323 | 0,009714519737 | 0,000007728586594 | 22,29008073 | 0,00024796371 |
| AF_18295 | CRNKL1 | 0,5838243385 | 0,5832332323 | 0,0005911061323 | 33,73969962 | 0,000002179976614 |
| AF_18346 | PNPT1 | 0,01501970879 | 0,0136120458 | 0,001407662992 | 14,41684037 | 0,006716466652 |
| AF_18631 | ABCC8 | 0,01044263352 | 0,009131848594 | 0,001310784928 | 31,96411953 | 0,000004342938808 |
| AF_18668 | ST5 | 0,004954102911 | 0,003450331129 | 0,001503771781 | 10,0328922 | 0,04247787112 |
| AF_18676 | TMEM41B | 0,01280678167 | 0,01280085299 | 0,000005928681768 | 20,24308439 | 0,000577192499 |
| AF_18686 | RNF141 | 3,57930238 | 3,577819496 | 0,001482883981 | 36,20685391 | 0,0000007951871368 |
| AF_18738 | ELP4 | 2,534276948 | 2,533197341 | 0,001079607515 | 10,26397757 | 0,03870671253 |
| AF_18857 | CELSR1 | 0,04036886118 | 0,03914367914 | 0,001225182043 | 22,48491382 | 0,0002327211435 |
| AF_18889 | ATXN7L1 | 0,005309448674 | 0,004791370254 | 0,0005180784204 | 13,50647971 | 0,009646481236 |
| AF_18986 | TRAF3IP2 | 0,3127519785 | 0,3120624634 | 0,0006895151168 | 11,27010845 | 0,02516972731 |

**Table S3. Candidate genes predicted to be under positive selection by aBSREL models in *A. patagonicus* (FDR < 0.05)**

| Gene ID | Gene name | Full adaptive model | Full adaptive model (non-synonymous subs/site) | Full adaptive model (synonymous subs/site) | LRT | Adjusted p-value |
|---|---|---|---|---|---|---|
| AF_57 | DDB1 | 0,005937256786 | 0,003816996435 | 0,002120260351 | 12,65945415 | 0,02702669133 |
| AF_214 | NFX1 | 0,1215834258 | 0,1214902985 | 0,00009312724318 | 28,31927794 | 0,00003462003615 |
| AF_218 | TNS3 | 0,06475598911 | 0,06245494489 | 0,002301044226 | 41,6297189 | 0,000001076302137 |
| AF_285 | WDR35 | 0,004366266236 | 0,004103820558 | 0,0002624456784 | 12,2118068 | 0,03224737962 |
| AF_341 | OPN4 | 18,64797477 | 18,64446331 | 0,003511456269 | 19,76381782 | 0,001144790352 |
| AF_536 | CASQ2 | 2,30682873 | 2,30593735 | 0,0008913794655 | 44,9319042 | 0,00000003088874603 |
| AF_791 | ABCA3 | 0,0228990592 | 0,02096447155 | 0,001934587652 | 20,15170574 | 0,0009675710921 |
| AF_1084 | DNAJC13 | 0,1260427786 | 0,1255885172 | 0,0004542613353 | 21,1763807 | 0,0006172599229 |
| AF_1498 | AIM1 | 0,4318026211 | 0,4311080841 | 0,0006945370328 | 31,98630951 | 0,000008858236032 |
| AF_1638 | NAT10 | 1,295176535 | 1,293324391 | 0,00185214437 | 12,557389 | 0,02801770992 |
| AF_1698 | TMEM248 | 0,04155954323 | 0,03531877362 | 0,006240769616 | 17,30733287 | 0,003250466448 |
| AF_1727 | CEP126 | 0,0106435014 | 0,009887081362 | 0,0007564200379 | 17,51882338 | 0,00303242132 |
| AF_1784 | SLC15A1 | 0,1155202955 | 0,1138052254 | 0,001715070054 | 19,08108845 | 0,001495208243 |
| AF_1795 | NUDT14 | 7,346380748 | 7,342040484 | 0,004340263939 | 12,72341343 | 0,02637023142 |
| AF_1885 | TGS1 | 0,9212590184 | 0,9190020599 | 0,002256958548 | 20,16956129 | 0,0009675710921 |
| AF_2026 | SCLT1 | 0,0362501338 | 0,03573915626 | 0,0005109775401 | 14,04905263 | 0,01435926029 |
| AF_2078 | DAGLA | 0,3889938515 | 0,3877082609 | 0,001285590582 | 24,11500136 | 0,0001907040367 |
| AF_2094 | ECE1 | 0,1768274972 | 0,1732710105 | 0,003556486673 | 14,12523029 | 0,013905064 |
| AF_2274 | CCDC80 | 0,8409098272 | 0,8394889408 | 0,001420886431 | 18,94700597 | 0,001586524715 |
| AF_2331 | ZPLD1 | 0,01138594996 | 0,01137010762 | 0,00001584234518 | 17,31808283 | 0,003250466448 |
| AF_2397 | SOS2 | 0,01197343273 | 0,01163223884 | 0,0003411938901 | 31,07172932 | 0,00001211700761 |
| AF_2403 | POLE2 | 0,4648494109 | 0,4640828638 | 0,0007665471081 | 11,31862209 | 0,04626827607 |
| AF_2501 | CBX7 | 0,1115915889 | 0,1115746732 | 0,00001691570334 | 22,90411252 | 0,0003135345715 |
| AF_2516 | TNRC6B | 0,09650260734 | 0,09176857126 | 0,004734036083 | 31,8771465 | 0,000009072908296 |
| AF_2550 | MEI1 | 0,8980263035 | 0,8978212401 | 0,0002050634317 | 28,8349229 | 0,00002866744234 |
| AF_2630 | ATP7B | 5,385564968 | 5,381311937 | 0,004253030723 | 15,10593516 | 0,008874247141 |
| AF_2637 | CKAP2 | 0,07329966265 | 0,07142948562 | 0,001870177034 | 21,20869589 | 0,0006132203986 |
| AF_2702 | ALOX5AP | 12,85255407 | 12,85021811 | 0,002335959454 | 29,51480309 | 0,00002289533282 |
| AF_2715 | FLT3 | 86,9053855 | 86,89511285 | 0,01027265134 | 39,84005141 | 0,0000002408430043 |
| AF_2734 | NUP58 | 0,646803744 | 0,6462148664 | 0,0005888775714 | 27,62047299 | 0,00004685178587 |
| AF_2837 | BAZ1A | 0,004478273113 | 0,00315739915 | 0,001320873963 | 19,43910981 | 0,001301960977 |
| AF_2954 | JMJD1C | 0,004825940621 | 0,004214545055 | 0,0006113955661 | 11,87052306 | 0,03707797701 |
| AF_3110 | DYSF | 0,0190449074 | 0,01211991786 | 0,006924989539 | 26,02817922 | 0,00008546331066 |
| AF_3231 | ZDHHC1 | 0,02957741287 | 0,02876729329 | 0,0008101195774 | 11,94496346 | 0,03609706339 |
| AF_3359 | COL3A1 | 0,06392492151 | 0,06379783217 | 0,0001270893435 | 12,65159143 | 0,02702669133 |
| AF_3417 | CNTN5 | 0,008446959317 | 0,007104054928 | 0,001342904389 | 11,4932885 | 0,04323623222 |
| AF_3573 | PSMA5 | 0,1272491479 | 0,1271773918 | 0,00007175613363 | 12,34330626 | 0,03057114021 |
| AF_3806 | PPP1R21 | 2,310173053 | 2,307885904 | 0,002287148705 | 16,96122623 | 0,003840600344 |
| AF_3816 | SYCP1 | 0,5968355815 | 0,5960697152 | 0,000765866358 | 11,87359714 | 0,03707797701 |
| AF_3839 | MAGI3 | 0,02640441398 | 0,02560561342 | 0,00079880056 | 17,48113501 | 0,003044648715 |
| AF_3906 | SLC26A9 | 0,8401014525 | 0,8379518741 | 0,002149578351 | 25,49443551 | 0,000104307093 |
| AF_4207 | CACNB4 | 0,8133276157 | 0,8130654314 | 0,0002621843164 | 61,37504353 | 1,92E-11 |
| AF_4405 | COPB2 | 0,04975699746 | 0,04812351352 | 0,00163348394 | 15,31447996 | 0,008095727715 |
| AF_4494 | LNX2 | 0,00999752087 | 0,00882379806 | 0,00117372281 | 27,79587967 | 0,00004371597081 |
| AF_4545 | GOLGA4 | 0,01043916717 | 0,009585539135 | 0,0008536280333 | 20,3811921 | 0,0008780463366 |
| AF_4651 | CFI | 0,009802179791 | 0,008954659774 | 0,0008475200172 | 13,52973498 | 0,01820145221 |
| AF_4689 | ADGRL3 | 11,28889889 | 11,28473103 | 0,004167859745 | 35,8354935 | 0,000001525397869 |
| AF_4724 | C3 | 0,1210142544 | 0,1209499008 | 0,00006435357467 | 12,08231755 | 0,03404816296 |
| AF_4869 | CAPN8 | 0,02306175111 | 0,02212865713 | 0,0009330939781 | 25,51920655 | 0,000104307093 |
| AF_4955 | OGFR | 0,2139703472 | 0,2100508983 | 0,003919448902 | 32,65089359 | 0,000006553463965 |
| AF_4961 | CABLES2 | 10,29838566 | 10,29568735 | 0,002698310482 | 38,17475937 | 0,0000005102870979 |
| AF_4973 | TAF4 | 0,07434304415 | 0,0720697597 | 0,002273284451 | 41,83413208 | 0,0000001020140432 |
| AF_5019 | CA12 | 0,04989817566 | 0,04845814259 | 0,001440033068 | 14,4427141 | 0,01207669995 |
| AF_5091 | DPM1 | 0,9940052605 | 0,9938876027 | 0,000117657811 | 54,48970093 | 0,0000000004013031574 |
| AF_5241 | PTPN1 | 0,3932645643 | 0,3926030608 | 0,0006615034975 | 22,26084629 | 0,0003879292634 |
| AF_5347 | CHFR | 0,0466541824 | 0,04511326915 | 0,001540913249 | 18,22301085 | 0,002194349056 |
| AF_5419 | CD3D | 0,3261298979 | 0,3102344846 | 0,01589541327 | 16,01016986 | 0,005940428242 |
| AF_5565 | ZNF654 | 0,1488430892 | 0,1487683151 | 0,00007477401938 | 28,94230134 | 0,00002774271789 |
| AF_5570 | CD200 | 15,73677197 | 15,73052902 | 0,006242956027 | 19,81071991 | 0,001128026379 |
| AF_5615 | GAK | 0,08683904118 | 0,08571334788 | 0,001125693307 | 52,87304854 | 0,0000000008112921834 |
| AF_5865 | LY75 | 3,512209313 | 3,511070099 | 0,001139213737 | 18,55725516 | 0,001874189439 |
| AF_5896 | SLC23A3 | 0,01875259688 | 0,01720124022 | 0,001551356663 | 11,16096182 | 0,0496015022 |
| AF_5947 | TMEM169 | 0,5468122407 | 0,5465149977 | 0,0002972429473 | 34,96450476 | 0,000002275769571 |
| AF_5970 | TFCP2L1 | 0,005271244559 | 0,005267530807 | 0,000003713752527 | 19,98777543 | 0,001041240433 |
| AF_6157 | ARHGAP21 | 0,9851172736 | 0,9849471124 | 0,0001701612221 | 29,33093431 | 0,00002416332797 |
| AF_6166 | DNAJC13 | 0,8634884026 | 0,8619839937 | 0,0015044089 | 24,86767695 | 0,0001360173224 |
| AF_6198 | COL4A5 | 0,00708270317 | 0,005500055117 | 0,001582648053 | 12,33965266 | 0,03057114021 |
| AF_6340 | PPIF | 5,946557408 | 5,944009458 | 0,002547950001 | 16,11612038 | 0,005671162177 |
| AF_6369 | CDH17 | 0,4458470753 | 0,4453726632 | 0,0004744120952 | 19,52918736 | 0,001265805263 |
| AF_6433 | FAM83D | 0,6181919159 | 0,6173526063 | 0,0008393096024 | 18,68390608 | 0,001796307321 |
| AF_6452 | FAM49A | 0,1245999598 | 0,1231064365 | 0,001493523275 | 11,4470776 | 0,04402981684 |
| AF_6454 | DDX1 | 0,084665123 | 0,0816170281 | 0,003048094904 | 49,54142743 | 0,000000003909759153 |
| AF_6591 | FAM188A | 3,043542224 | 3,041021535 | 0,00252068987 | 44,03719957 | 0,00000004512560925 |
| AF_6649 | SLC17A5 | 0,006489439712 | 0,006478549352 | 0,00001089035999 | 14,64739178 | 0,01103452145 |
| AF_6680 | UGGT2 | 0,05938174984 | 0,05746137467 | 0,001920375175 | 19,20868438 | 0,001425530652 |
| AF_6686 | PDC | 8,822326155 | 8,819536929 | 0,002789226166 | 30,78681608 | 0,00001326053273 |
| AF_6753 | TOR1AIP2 | 0,747478393 | 0,7468068629 | 0,0006715300893 | 23,51808116 | 0,0002502762899 |
| AF_6791 | CRYD | 0,0339350755 | 0,03138360271 | 0,002551472788 | 12,28429642 | 0,0312621719 |
| AF_6901 | TNFSF13B | 0,1753293474 | 0,1689726398 | 0,006356707562 | 19,16958051 | 0,00144190114 |
| AF_7006 | EIF5B | 0,06438983034 | 0,06411205399 | 0,0002777763449 | 13,73440845 | 0,01651796065 |
| AF_7288 | TMOD3 | 6,680221854 | 6,677833329 | 0,002388525235 | 17,38557355 | 0,003171026065 |
| AF_7443 | EXOC3 | 0,01992274379 | 0,01842913429 | 0,001493609501 | 31,68155704 | 0,000009713131936 |
| AF_7486 | CNDP1 | 13,77947704 | 13,77547716 | 0,003999873677 | 12,59792596 | 0,02760851487 |
| AF_7552 | DPEP2 | 2,477066367 | 2,476090856 | 0,0009755113167 | 21,6348358 | 0,0004999456272 |
| AF_7615 | ENDOV | 0,2092887314 | 0,2076667221 | 0,001622009238 | 30,67394 | 0,00001359944136 |
| AF_7639 | SLC30A10 | 0,4408796802 | 0,4321053288 | 0,008774351395 | 30,63671497 | 0,00001359944136 |
| AF_7884 | SMARCD3 | 0,04219367715 | 0,04086487596 | 0,001328801196 | 24,7717462 | 0,0001389685219 |
| AF_7903 | TMEFF2 | 0,114813153 | 0,1134745615 | 0,001338591459 | 15,62412941 | 0,007067866967 |
| AF_8067 | USP10 | 0,008389001634 | 0,006990045023 | 0,001398956611 | 19,30746193 | 0,001379303658 |
| AF_8091 | GCSH | 114,2037949 | 114,1987998 | 0,004995117907 | 26,6949493 | 0,00006938885791 |
| AF_8125 | CENPU | 1,276125107 | 1,27499598 | 0,001129127848 | 12,50645047 | 0,02858347334 |
| AF_8128 | CFAP97 | 1,155060648 | 1,15337955 | 0,001681097249 | 19,49854761 | 0,001274455305 |
| AF_8314 | RTF1 | 0,01616115098 | 0,01612500485 | 0,00003614612483 | 11,54430061 | 0,04235534591 |
| AF_8336 | PLA2G4E | 0,1612694654 | 0,1608109976 | 0,0004584678594 | 23,49716465 | 0,0002502762899 |
| AF_8402 | SYNPO2L | 0,08197724841 | 0,07866437743 | 0,003312870972 | 62,95063231 | 1,04E-11 |
| AF_8533 | RBP4 | 2,363669703 | 2,362139384 | 0,001530318449 | 22,6432056 | 0,000345360127 |
| AF_8573 | GOT1 | 2,422247909 | 2,419506566 | 0,002741343911 | 13,43529678 | 0,01886002144 |
| AF_8702 | CCDC186 | 0,01002221987 | 0,007353311669 | 0,0026689082 | 14,16759755 | 0,01378376332 |

| | | | | | | |
|---|---|---|---|---|---|---|
| AF_8752 | WDR11 | 0,1461561642 | 0,1438428488 | 0,002313315401 | 23,3415679 | 0,0002608237381 |
| AF_8774 | ADAM9 | 0,006535936278 | 0,005501729257 | 0,001034207022 | 11,75965813 | 0,0383904482 |
| AF_8799 | BCCIP | 0,3273939273 | 0,3273036919 | 0,00009023545302 | 36,04471603 | 0,000001426414755 |
| AF_8923 | TRPV1 | 0,3330752988 | 0,3325756602 | 0,0004996386053 | 18,55226595 | 0,001874189439 |
| AF_8924 | TRPV3 | 0,02583313376 | 0,02204617717 | 0,003786956594 | 22,38489193 | 0,000376266613 |
| AF_8965 | RPS6KB1 | 0,1695594819 | 0,1694306037 | 0,000128878211 | 26,2429858 | 0,00008020581505 |
| AF_9307 | OSBPL6 | 0,7087270231 | 0,7079967564 | 0,0007302666988 | 43,34042205 | 0,00000005330741007 |
| AF_9437 | HSDL2 | 9,831365043 | 9,829351684 | 0,002013358979 | 26,54191774 | 0,00007242736882 |
| AF_9565 | KCNC2 | 0,0385876983 | 0,03853338366 | 0,00005431463228 | 11,85111612 | 0,03707922591 |
| AF_9572 | KRR1 | 0,06214049059 | 0,06100617066 | 0,001134319925 | 12,94767748 | 0,02368938086 |
| AF_9658 | SLCO1C1 | 16,19914585 | 16,19393326 | 0,005212583656 | 29,30452619 | 0,00002416332797 |
| AF_9667 | ETNK1 | 1,927112457 | 1,925392604 | 0,001719852722 | 21,72439402 | 0,0004826945679 |
| AF_9693 | ASUN | 0,01328404667 | 0,01324965872 | 0,00003438794802 | 15,88288159 | 0,006289965573 |
| AF_9717 | SCUBE1 | 0,02071437324 | 0,01940862847 | 0,001305744772 | 15,5836303 | 0,007164986827 |
| AF_9727 | PARVG | 2,765459629 | 2,765208466 | 0,0002511632831 | 56,28562407 | 0,00000000002098989554 |
| AF_9977 | MICU2 | 0,03369875291 | 0,02839806267 | 0,005300690241 | 14,51478196 | 0,01172091522 |
| AF_10112 | TBCD | 1,473139253 | 1,47150554 | 0,001633712746 | 22,82651015 | 0,0003222417228 |
| AF_10515 | DIAPH2 | 0,01093249756 | 0,009600196701 | 0,001332300855 | 17,69152841 | 0,002822662853 |
| AF_10527 | DDX51 | 0,9378915608 | 0,9369184145 | 0,0009731462513 | 41,91745476 | 0,000001020140432 |
| AF_10579 | CUNH12orf43 | 0,3067495143 | 0,2972392045 | 0,009510309743 | 13,26596401 | 0,02029717954 |
| AF_10588 | RNF10 | 0,4436387246 | 0,4426528457 | 0,0009858789244 | 19,24383315 | 0,001412246449 |
| AF_10741 | GALNT1 | 0,05249943537 | 0,05242477381 | 0,00007466156625 | 26,81436721 | 0,00006650121601 |
| AF_10753 | FH | 9,109293291 | 9,1050282 | 0,004265090676 | 18,66067928 | 0,001802974854 |
| AF_10875 | FAM189A1 | 0,009046161982 | 0,00822764125 | 0,0008185207323 | 11,24250753 | 0,04783957162 |
| AF_10955 | CTDSPL2 | 0,2586738739 | 0,2586127689 | 0,00006110493966 | 23,07278526 | 0,000291471255 |
| AF_11068 | PHC3 | 0,1043667433 | 0,1042776234 | 0,00008911992996 | 22,03450677 | 0,0004204624574 |
| AF_11070 | SKIL | 0,03518970408 | 0,03470647056 | 0,0004832335171 | 29,50270627 | 0,00002289533282 |
| AF_11086 | GOLGA1 | 0,2649574131 | 0,2641556125 | 0,0008018005803 | 16,60798332 | 0,004522251395 |
| AF_11248 | GPATCH2L | 0,5272433474 | 0,526413675 | 0,0008296723463 | 22,01912677 | 0,0004204624574 |
| AF_11322 | PSMC1 | 0,03608322582 | 0,0360594605 | 0,00002376531881 | 34,3364666 | 0,000003010004238 |
| AF_11503 | SSBP2 | 2,889753203 | 2,886874064 | 0,002879139425 | 17,4881303 | 0,003044648715 |
| AF_11754 | NCAPD3 | 0,004277597829 | 0,003415605102 | 0,0008619927269 | 26,02197676 | 0,00008546331066 |
| AF_12130 | DIS3 | 0,0265844982 | 0,02619206382 | 0,000392434382 | 16,47673277 | 0,004796872524 |
| AF_12249 | LMOD1 | 1,707949253 | 1,701768289 | 0,006180963946 | 23,7982331 | 0,0002206959013 |
| AF_12343 | UBE2J1 | 0,08217479008 | 0,08214845041 | 0,00002633967597 | 45,11499681 | 0,00000003088874603 |
| AF_12372 | NUDCD1 | 0,6474017875 | 0,64694441 | 0,00045737748 | 44,95066892 | 0,00000003088874603 |
| AF_12511 | TNMD | 0,2487106681 | 0,2486043251 | 0,0001063429585 | 33,31839975 | 0,000004846414014 |
| AF_12856 | VRK2 | 0,7497148463 | 0,7490378137 | 0,0006770325602 | 28,29904589 | 0,00003462003615 |
| AF_12981 | LARP7 | 2,197850282 | 2,195756042 | 0,002094240158 | 20,87719327 | 0,0007037982417 |
| AF_13114 | HM13 | 4,578041731 | 4,574028862 | 0,004012869542 | 13,46165636 | 0,01872325621 |
| AF_13220 | IKZF1 | 0,4960034325 | 0,4952452513 | 0,0007581811588 | 22,42235955 | 0,000376266613 |
| AF_13259 | CCT8 | 8,997546764 | 8,994026772 | 0,003519992113 | 29,11579565 | 0,00002598415929 |
| AF_13272 | SCAF4 | 0,4716506825 | 0,4707024848 | 0,0009481976874 | 25,78741068 | 0,00009343959957 |
| AF_13292 | SON | 0,2346211319 | 0,223208303 | 0,01141282895 | 71,12322062 | 4,24E-13 |
| AF_13382 | CUNH21orf33 | 0,3145793075 | 0,3144467905 | 0,000132517005 | 25,21814982 | 0,0001172534902 |
| AF_13596 | SRP72 | 0,02897017681 | 0,02869252851 | 0,0002776483007 | 20,8297916 | 0,0007140178182 |
| AF_13611 | SH3D19 | 3,013563446 | 3,011529417 | 0,002034029729 | 11,78537422 | 0,0380948584 |
| AF_13616 | TMEM154 | 0,1356337981 | 0,130445048 | 0,005188750067 | 26,06113563 | 0,00008546331066 |
| AF_13632 | DCHS2 | 4,894872716 | 4,8853898 | 0,009482915365 | 69,9531184 | 5,30E-13 |
| AF_13635 | FGA | 0,02268006755 | 0,02146318058 | 0,001216886974 | 22,5340846 | 0,0003607525068 |
| AF_13646 | CTSO | 0,1320539404 | 0,1313226428 | 0,000731297664 | 24,88376938 | 0,0001360173224 |
| AF_13984 | TALDO1 | 1,556107402 | 1,55500152 | 0,00110588189 | 22,07535572 | 0,0004171007351 |
| AF_14038 | MRPL14 | 212,5345299 | 212,5193111 | 0,01521882664 | 43,74964358 | 0,00000004598401072 |
| AF_14073 | FEZ2 | 0,09731074674 | 0,09726594276 | 0,00004480397809 | 26,24128272 | 0,00008020581505 |
| AF_14104 | MYO1H | 0,00950198102 | 0,008570097574 | 0,0009318834465 | 26,57231915 | 0,00007242736882 |
| AF_14160 | TCTN1 | 1,345407591 | 1,344529592 | 0,0008779987846 | 13,37036521 | 0,01937140393 |
| AF_14262 | PLIN3 | 20,67591391 | 20,66950574 | 0,006408168858 | 23,46570251 | 0,0002511206374 |
| AF_14348 | PSMG2 | 0,06679916162 | 0,06539511107 | 0,001404050548 | 15,8543084 | 0,006337645332 |
| AF_14366 | RALBP1 | 5,387881296 | 5,38587416 | 0,002007135953 | 31,50036849 | 0,00001033269931 |
| AF_14429 | YTHDC2 | 2,561987549 | 2,56095862 | 0,001028929029 | 55,79094924 | 0,00000000002353561721 |
| AF_14480 | SEPT2 | 0,2494469262 | 0,2482360804 | 0,001210845809 | 23,20354862 | 0,00027620468 |
| AF_14481 | FARP2 | 0,01651661482 | 0,0123462809 | 0,004170333927 | 28,76834849 | 0,00002903567217 |
| AF_14563 | IWS1 | 0,01279021242 | 0,007457027493 | 0,005333184923 | 13,95422893 | 0,01496884219 |
| AF_14715 | UROC1 | 2,000200281 | 1,999309926 | 0,000890354592 | 43,78477964 | 0,00000004598401072 |
| AF_15048 | PPIE | 0,5395259464 | 0,5394727057 | 0,00005324072776 | 22,79339639 | 0,0003239195542 |
| AF_15058 | PLEKHJ1 | 28,89213649 | 28,88745705 | 0,00467943846 | 26,25672607 | 0,00008020581505 |
| AF_15086 | COG3 | 0,5369797312 | 0,5364435958 | 0,0005361353999 | 22,40473377 | 0,000376266613 |
| AF_15093 | CPB2 | 0,1281658489 | 0,1280722853 | 0,00009356360449 | 14,14427404 | 0,01385893593 |
| AF_15149 | MAP3K13 | 0,04484568375 | 0,04092908707 | 0,003916596681 | 16,77068296 | 0,004196674108 |
| AF_15160 | CLDN18 | 0,3560735771 | 0,3011926911 | 0,05488088593 | 27,19663795 | 0,00005587893808 |
| AF_15461 | PIP5K1B | 0,02356036353 | 0,01945238511 | 0,004107978418 | 12,08549077 | 0,03404816296 |
| AF_15487 | ATF6 | 9,895815174 | 9,891212037 | 0,004603136747 | 18,13050126 | 0,002281115606 |
| AF_15529 | HTT | 4,085212757 | 4,082987655 | 0,002225102845 | 11,58363008 | 0,04173899905 |
| AF_15558 | LETM1 | 0,02425854547 | 0,02142006516 | 0,002838480302 | 38,62300544 | 0,0000004246402379 |
| AF_15623 | CAMK1D | 0,1351989764 | 0,1299962754 | 0,005202700978 | 11,8493755 | 0,03707922591 |
| AF_15642 | PRPF18 | 0,3555883672 | 0,3554802453 | 0,0001081218483 | 22,19351729 | 0,0003971292174 |
| AF_15711 | WDR43 | 0,2422732853 | 0,2417170643 | 0,0005562209572 | 11,96470837 | 0,03593343789 |
| AF_15753 | MEP1B | 40,57629195 | 40,56990371 | 0,006388235188 | 70,11910142 | 5,30E-13 |
| AF_15815 | LCP2 | 0,01263990306 | 0,01261164507 | 0,00002825798433 | 24,83930242 | 0,0001361280674 |
| AF_15900 | BRD8 | 0,02948859475 | 0,02947729122 | 0,00001130353151 | 17,56778632 | 0,002981094261 |
| AF_15959 | NIPAL4 | 7,924006813 | 7,920356133 | 0,003650679888 | 40,45011765 | 0,0000001854923599 |
| AF_16001 | ATOX1 | 1,483983821 | 1,483858444 | 0,0001253774638 | 23,37064844 | 0,0002601791948 |
| AF_16408 | CALM2 | 0,6508186902 | 0,6497704949 | 0,001048195239 | 12,36721934 | 0,03048965803 |
| AF_16658 | PLK1 | 0,05615625732 | 0,054427514 | 0,001728743319 | 22,33124933 | 0,0003824248869 |
| AF_16805 | IL17REL | 13,66595343 | 13,66206709 | 0,003886340809 | 16,1713283 | 0,005554379625 |
| AF_17266 | ATP6V0B | 9,827781262 | 9,825749531 | 0,002031730657 | 22,30875436 | 0,0003826968443 |
| AF_17368 | ORC1 | 0,6165499894 | 0,6105594671 | 0,005990522262 | 30,89050128 | 0,00001292017622 |
| AF_17412 | USP24 | 0,002603733902 | 0,002164618161 | 0,0004391157419 | 13,82968248 | 0,01584038194 |
| AF_17513 | FAM122B | 1,96981135 | 1,965795706 | 0,004015643797 | 11,36589829 | 0,04540622131 |
| AF_17750 | TBL1X | 1,940134343 | 1,937430497 | 0,002703846587 | 19,6254957 | 0,001216502581 |
| AF_17802 | SCN4A | 1,401322444 | 1,400118561 | 0,001203883282 | 15,25039157 | 0,008306136691 |
| AF_17859 | NLRX1 | 0,4709383127 | 0,4704025835 | 0,0005357291877 | 31,39029 | 0,00001061556539 |
| AF_17888 | HEPH | 2,785244099 | 2,784785105 | 0,0004589934151 | 25,78459945 | 0,00009343959957 |
| AF_18206 | KIAA1551 | 0,003909069604 | 0,003178850134 | 0,0007302194707 | 20,93846262 | 0,0006889718636 |
| AF_18227 | AGK | 6,087107303 | 6,084514386 | 0,002592916646 | 28,68015833 | 0,00002974152019 |
| AF_18243 | SVOPL | 14,90774948 | 14,90194904 | 0,005800432981 | 14,73519546 | 0,01062585061 |
| AF_18275 | HSP90B1 | 0,1708640743 | 0,1707363217 | 0,0001277526462 | 15,533971 | 0,007297335766 |
| AF_18433 | MCPH1 | 0,1376417404 | 0,1376232463 | 0,00001849417126 | 27,31769241 | 0,00005354338037 |
| AF_18616 | LDHA | 0,07563674084 | 0,07465889987 | 0,0009778409774 | 25,47923096 | 0,000104307093 |
| AF_18858 | TRMU | 0,01637471855 | 0,01636704877 | 0,000007669783809 | 11,39126431 | 0,04505711381 |
| AF_18859 | GTSE1 | 24,75972041 | 24,75315282 | 0,006567588785 | 20,60242586 | 0,0007929791936 |

**Table S4. Candidate genes predicted to be under positive selection by CODEML branch models in *A. forsteri* and *A. patagonicus* (FDR < 0.05)**

| Lineage | Gene ID | Gene name | Gene description | Likelihood Null Model | Likelihood Alternative Model | dN | dS | ω background | ω foreground | Adjusted p-value |
|---|---|---|---|---|---|---|---|---|---|---|
| *A. forsteri* | AF_4464 | POLR1D | DNA-directed RNA polymerases I and III subunit RPAC2 | -1073650663 | -106755323 | 0,028517 | 0,036234 | 0,05409 | 0,787 | 0,04364804417 |
| *A. forsteri* | AF_4207 | CACNB4 | Calcium voltage-gated channel auxiliary subunit beta 4 | -2433426797 | -2418667129 | 0,011123 | 0,008163 | 0,0308 | 1,3626 | 6,05E-05 |
| *A. forsteri* | AF_4640 | SGMS2 | Sphingomyelin synthase 2 | -297537272 | -2966389517 | 0,007836 | 0,003428 | 0,06737 | 2,2862 | 0,005059561294 |
| *A. forsteri* | AF_4559 | ASAP1 | ArfGAP with SH3 domain, ankyrin repeat and PH domain 1 | -6296899247 | -6290455269 | 0,002263 | 0,002701 | 0,05016 | 0,8377 | 0,03520841431 |
| *A. forsteri* | AF_4973 | TAF4 | TATA-box binding protein associated factor 4 | -2421578647 | -2411889524 | 0,0101 | 0,001486 | 0,05708 | 6,7988 | 0,002646258774 |
| *A. forsteri* | AF_5071 | TFAP2C | Transcription factor AP-2 gamma | -2007770972 | -1995712463 | 0,009457 | 0,009425 | 0,02139 | 1,0033 | 0,0004080053271 |
| *A. forsteri* | AF_5352 | ZDHHC8 | Zinc finger DHHC-type palmitoyltransferase 8 | -627925246 | -62729554 | 0,005392 | 0,00724 | 0,0939 | 0,7448 | 0,04000852649 |
| *A. forsteri* | AF_5369 | SEPTIN2 | Septin-2 | -232271942 | -2307164315 | 0,00471 | 0,008085 | 0,03002 | 0,5826 | 3,81E-05 |
| *A. forsteri* | AF_5373 | UFD1L | Ubiquitin recognition factor in ER associated degradation 1 | -2206514633 | -2197794535 | 0,007362 | 0,009361 | 0,02117 | 0,7865 | 0,006301236083 |
| *A. forsteri* | AF_5984 | ADCY5 | Adenylate cyclase 5 | -4773199909 | -4765414796 | 0,00906 | 0,016187 | 0,07598 | 0,5597 | 0,01267149056 |
| *A. forsteri* | AF_6996 | RPL31 | Ribosomal protein L31 | -1191786381 | -1184957119 | 0,019555 | 0,031658 | 0,04402 | 0,6177 | 0,027956754 |
| *A. forsteri* | AF_7017 | GMEB1 | Glucocorticoid modulatory element binding protein 1 | -2945127035 | -2939114773 | 0,003496 | 0,011999 | 0,01184 | 0,2914 | 0,04617422471 |
| *A. forsteri* | AF_6879 | AP2A2 | Adaptor related protein complex 2 subunit alpha 2 | -6843745376 | -6835199098 | 0,002413 | 0,003286 | 0,01918 | 0,7344 | 0,007058734128 |
| *A. forsteri* | AF_6831 | WNT3A | Wnt family member 3A | -2339459998 | -2319832376 | 0,01204 | 0,011843 | 0,01389 | 1,0166 | 1,42E-06 |
| *A. forsteri* | AF_7895 | GLS | Glutaminase | -3206570464 | -3196047445 | 0,007334 | 0,003744 | 0,04538 | 1,959 | 0,00117305132 |
| *A. forsteri* | AF_7908 | ATP6V1C1 | ATPase H+ transporting V1 subunit C1 | -3093405935 | -3086738859 | 0,011505 | 0,02447 | 0,07101 | 0,4702 | 0,03216134871 |
| *A. forsteri* | AF_8749 | MCMBP | Minichromosome maintenance complex binding protein | -5495090419 | -5485779913 | 0,007091 | 0,006285 | 0,07152 | 1,1281 | 0,003696824091 |
| *A. forsteri* | AF_8543 | IDE | Insulin degrading enzyme | -6904648351 | -6896112142 | 0,005545 | 0,027849 | 0,03653 | 0,1991 | 0,007058734128 |
| *A. forsteri* | AF_8972 | INTS2 | Integrator complex subunit 2 | -10334032792 | -10327810599 | 0,002343 | 0,002821 | 0,05648 | 0,8306 | 0,04157785835 |
| *A. forsteri* | AF_8983 | APPBP2 | Amyloid beta precursor protein binding protein 2 | -4370923681 | -4354724513 | 0,004569 | 0,002776 | 0,00889 | 1,646 | 3,20E-05 |
| *A. forsteri* | AF_9132 | HDHD2 | Haloacid dehalogenase like hydrolase domain containing 2 | -2586355115 | -2580233995 | 0,013557 | 0,005209 | 0,13516 | 2,6025 | 0,04359203902 |
| *A. forsteri* | AF_9261 | DYNC1I2 | Dynein cytoplasmic 1 intermediate chain 2 | -4339346636 | -4331810605 | 0,003983 | 0,004721 | 0,03087 | 0,8437 | 0,0158345096 |
| *A. forsteri* | AF_9769 | SLIT2 | Slit guidance ligand 2 | -8311327808 | -8305127747 | 0,001522 | 0,00295 | 0,02838 | 0,5159 | 0,04157785835 |
| *A. forsteri* | AF_9862 | PHLPP1 | PH domain and leucine rich repeat protein phosphatase 1 | -9858850225 | -9850650474 | 0,002592 | 0,001983 | 0,06277 | 1,3074 | 0,009436597667 |
| *A. forsteri* | AF_9984 | XPO4 | Exportin 4 | -7237695777 | -7227132988 | 0,002114 | 0,002742 | 0,02107 | 0,771 | 0,00117305132 |
| *A. forsteri* | AF_9961 | ANKRD17 | Ankyrin repeat domain 17 | -15007969513 | -14992613022 | 0,003813 | 0,003778 | 0,05029 | 1,0093 | 3,81E-05 |
| *A. forsteri* | AF_10572 | SFI1 | SFI1 centrin binding protein | -3940653339 | -3932635808 | 0,001847 | 0,060968 | 0,54726 | 0,0303 | 0,01081225409 |
| *A. forsteri* | AF_10519 | SMARCB1 | Actin dependent regulator of chromatin | -2590451276 | -2582187836 | 0,006447 | 0,012508 | 0,01855 | 0,5154 | 0,009174505375 |
| *A. forsteri* | AF_11846 | PCMT1 | Protein-L-isoaspartate (D-aspartate) O-methyltransferase | -1193624067 | -1187040217 | 0,005729 | 0,003375 | 0,02551 | 1,6972 | 0,033011879 |
| *A. forsteri* | AF_13292 | SON | SON DNA binding protein | -4358166602 | -43499761 | 0,028991 | 0,072681 | 0,10877 | 0,3989 | 0,009436597667 |
| *A. forsteri* | AF_12993 | ARSJ | Arylsulfatase family member J | -4474783053 | -4467715932 | 0,004169 | 0,005722 | 0,02725 | 0,7286 | 0,02348588544 |
| *A. forsteri* | AF_13270 | TIAM1 | TIAM Rac1 associated GEF 1 | -13787238749 | -13779297985 | 0,002102 | 0,002069 | 0,05724 | 1,016 | 0,01121603335 |
| *A. forsteri* | AF_262 | CTNNB1 | Catenin beta 1 | -5927657146 | -5916428017 | 0,002978 | 0,002427 | 0,01269 | 1,2269 | 0,00066380076 |
| *A. forsteri* | AF_475 | ZNF326 | Zinc finger protein 326 | -3086139466 | -3074893861 | 0,005437 | 0,00556 | 0,03129 | 0,9779 | 0,00066380076 |
| *A. forsteri* | AF_13872 | TEC | Tec protein tyrosine kinase | -483256932 | -4825957223 | 0,003355 | 0,004604 | 0,0502 | 0,7287 | 0,033011879 |
| *A. forsteri* | AF_14165 | RSRC2 | Arginine and serine rich coiled-coil 2 | -2871993365 | -2864573572 | 0,00697 | 0,00595 | 0,0417 | 1,1714 | 0,0175597951 |
| *A. forsteri* | AF_13907 | SLC2A1 | Solute carrier family 2 member 1 | -4273606712 | -426656548 | 0,00519 | 0,005713 | 0,02372 | 0,9085 | 0,02348588544 |
| *A. forsteri* | AF_14293 | SIN3B | SIN3 transcription regulator family member B | -8947294844 | -8931805361 | 0,003435 | 0,007034 | 0,01865 | 0,4884 | 3,81E-05 |
| *A. forsteri* | AF_14863 | CREB1 | cAMP responsive element binding protein 1 | -2182817514 | -2176441134 | 0,0045 | 0,003487 | 0,03012 | 1,2903 | 0,03725722575 |
| *A. forsteri* | AF_14970 | AGO4 | Protein argonaute-4 | -5831073447 | -5824543194 | 0,002199 | 0,002652 | 0,01364 | 0,8293 | 0,033011879 |
| *A. forsteri* | AF_14805 | CBFA2T2 | CBFA2/RUNX1 partner transcriptional co-repressor 2 | -486238352 | -4850439733 | 0,008462 | 0,020958 | 0,03234 | 0,4038 | 0,0004112613842 |
| *A. forsteri* | AF_15874 | UBTD2 | Ubiquitin domain containing 2 | -1426090087 | -1411632319 | 0,015211 | 0,012841 | 0,02579 | 1,1845 | 7,23E-05 |
| *A. forsteri* | AF_16001 | ATOX1 | Antioxidant 1 copper chaperone | -698565592 | -692311733 | 0,024614 | 0,070414 | 0,0778 | 0,3496 | 0,0413429436 |
| *A. forsteri* | AF_16232 | PAK1 | p21 (RAC1) activated kinase 1 | -3902088349 | -3893452186 | 0,005017 | 0,002471 | 0,0505 | 2,0305 | 0,006697106405 |
| *A. forsteri* | AF_16462 | TRRAP | Transformation/transcription domain associated protein | -1100710008 | -1094267791 | 0,007178 | 0,010158 | 0,007 | 0,7067 | 0,03520841431 |
| *A. forsteri* | AF_16779 | TMEM136 | Transmembrane protein 136 | -2554230876 | -2547536678 | 0,011799 | 0,005151 | 0,10675 | 2,2907 | 0,03175666705 |
| *A. forsteri* | AF_16783 | TRIM29 | Tripartite motif containing 29 | -4563715231 | -4556368765 | 0,005731 | 0,006242 | 0,05422 | 0,9181 | 0,01861547154 |
| *A. forsteri* | AF_17377 | ZYG11B | Protein zyg-11 homolog B | -5952338226 | -5941118625 | 0,004142 | 0,004208 | 0,03019 | 0,9843 | 0,00066380076 |
| *A. forsteri* | AF_17652 | PSMD12 | Proteasome 26S subunit, non-ATPase 12 | -2457245614 | -2449415142 | 0,006546 | 0,007682 | 0,03985 | 0,8522 | 0,01233487709 |
| *A. forsteri* | AF_17836 | CDC27 | Cell division cycle 27 | -5874928607 | -5868764267 | 0,003246 | 0,0087 | 0,03319 | 0,3731 | 0,04213245123 |
| *A. forsteri* | AF_867 | CLCN7 | Chloride voltage-gated channel 7 | -7135912477 | -7125413896 | 0,004365 | 0,00829 | 0,02712 | 0,5265 | 0,00117305132 |
| *A. forsteri* | AF_980 | DHX36 | DEAH-box helicase 36 | -8387828847 | -8379740228 | 0,002498 | 0,001433 | 0,04758 | 1,7426 | 0,0102635496 |
| *A. forsteri* | AF_18295 | CRNKL1 | Crooked neck pre-mRNA splicing factor 1 | -5307866494 | -5301030989 | 0,004075 | 0,00459 | 0,03059 | 0,8878 | 0,027956754 |
| *A. forsteri* | AF_18677 | IPO7 | Importin 7 | -6939111933 | -692536631 | 0,003364 | 0,001617 | 0,03016 | 2,081 | 0,0001121914818 |
| *A. forsteri* | AF_1408 | NUP205 | Nucleoporin 205 | -6807066793 | -6800013146 | 0,001181 | 0,007783 | 0,0716 | 0,1518 | 0,02348588544 |
| *A. forsteri* | AF_1588 | AQP2 | Aquaporin 2 | -3064184948 | -3057602995 | 0,008733 | 0,004594 | 0,07474 | 1,9011 | 0,033011879 |
| *A. forsteri* | AF_2852 | BRMS1L | BRMS1 like transcriptional repressor | -1970591685 | -1959043846 | 0,008633 | 0,008542 | 0,04364 | 1,0107 | 0,0005615834 |
| *A. forsteri* | AF_3784 | NEDD4 | NEDD4 E3 ubiquitin protein ligase | -6386335381 | -6380399578 | 0,004129 | 0,001973 | 0,11255 | 2,0925 | 0,04955413591 |
| *A. forsteri* | AF_15280 | TRPM8 | Transient receptor potential cation channel subfamily M member 8 | -9444417165 | -9435416073 | 0,001314 | 0,01346 | 0,06013 | 0,0976 | 0,01875770167 |
| *A. forsteri* | AF_2954 | JMJD1C | Jumonji domain containing 1C | -13288222845 | -132622255 | 0,001813 | 0,006211 | 0,12672 | 0,2919 | 4,26E-09 |
| *A. patagonicus* | AF_4494 | LNX2 | Ligand of Numb protein X 2 | -6658627707 | -6649558437 | 0,005742 | 0,004195 | 0,08009 | 1,3688 | 0,01208858 |
| *A. patagonicus* | AF_13259 | CCT8 | Chaperonin Containing TCP1 Subunit 8 | -3462889754 | -3455267095 | 0,009668 | 0,021091 | 0,0495 | 0,4584 | 0,038017819 |
| *A. patagonicus* | AF_15160 | CLDN18 | Claudin 18 | -3346636113 | -3331828403 | 0,030984 | 0,243117 | 0,07568 | 0,1274 | 0,000201542642 |
| *A. patagonicus* | AF_4689 | ADGRL3 | Adhesion G protein-coupled receptor L3 | -310995719 | -3099585622 | 0,008685 | 0,021118 | 0,02611 | 0,4113 | 0,005022785863 |
| *A. patagonicus* | AF_5762 | HNRNPLL | Heterogeneous nuclear ribonucleoprotein L like | -3456725962 | -3446531149 | 0,00538 | 0,002824 | 0,02437 | 1,9055 | 0,00503879558 |

**Table S5. Candidate genes for which RELAX found signals of intensified selection (K > 1) in *A. forsteri* (FDR < 0.05)**

| Gene ID | Gene name | Gene description | K | P-value | Adjusted p-value |
|---|---|---|---|---|---|
| AF_10572 | SFI1 | SFI1 Centrin Binding Protein | 5,135876433 | 0,0000000266 | 0,00009690635 |
| AF_10879 | TARS3 | Threonyl-TRNA Synthetase 3 | 24,97403162 | 0,0000000367 | 0,00009690635 |
| AF_10888 | MYO1E | Myosin IE | 40,54140993 | 0,0000000801 | 0,0001256878 |
| AF_11206 | COQ6 | Coenzyme Q6, Monooxygenase | 50 | 0,0000000952 | 0,0001256878 |
| AF_11502 | ACOT12 | Acyl-CoA Thioesterase 12 | 5,020793978 | 0,000000144 | 0,0001520928 |
| AF_12020 | OTOA | Otoancorin | 45,73485854 | 0,000000192 | 0,000168992 |
| AF_12479 | SNX13 | Sorting Nexin 13 | 3,675279974 | 0,000000299 | 0,0002255741429 |
| AF_12625 | SLC25A3 | Solute Carrier Family 25 Member 3 | 4,621722815 | 0,000000469 | 0,000309598625 |
| AF_12863 | KIAA1841 | SANT And BTB Domain Regulator Of CSR | 2,559263135 | 0,000000653 | 0,0003831658889 |
| AF_13392 | EFHC2 | EF-Hand Domain Containing 2 | 4,016546382 | 0,00000118 | 0,000623158 |
| AF_13473 | CRB1 | Crumbs Cell Polarity Complex Component 1 | 10,54184525 | 0,00000213 | 0,001022593636 |
| AF_13872 | TEC | Tec Protein Tyrosine Kinase | 50 | 0,00000307 | 0,001351055833 |
| AF_14164 | KNTC1 | Kinetochore Associated 1 | 50 | 0,00000363 | 0,001474617692 |
| AF_14805 | CBFA2T2 | CBFA2/RUNX1 Partner Transcriptional Co-Repressor 2 | 6,349842484 | 0,00000517 | 0,001946928667 |
| AF_14936 | INPP5B | Inositol Polyphosphate-5-Phosphatase B | 45,74236636 | 0,00000553 | 0,001946928667 |
| AF_1497 | ATG5 | Autophagy Related 5 | 11,97966152 | 0,00000666 | 0,00219821625 |
| AF_15354 | AGXT2 | Alanine--Glyoxylate Aminotransferase 2 | 50 | 0,0000072 | 0,002236658824 |
| AF_15426 | VPS13A | Vacuolar Protein Sorting 13 Homolog A | 4,692523499 | 0,00000912 | 0,002675706667 |
| AF_15710 | FAM179A | TOG Array Regulator Of Axonemal Microtubules 2 | 21,04335833 | 0,0000136 | 0,003780084211 |
| AF_15874 | UBTD2 | Ubiquitin Domain Containing 2 | 5,526435947 | 0,000015 | 0,00396075 |
| AF_16568 | DRC3 | Dynein Regulatory Complex Subunit 3 | 7,066727352 | 0,0000169 | 0,004249947619 |
| AF_16779 | TMEM136 | TLC Domain Containing 5 | 10,86567826 | 0,0000185 | 0,004440840909 |
| AF_16799 | LTA4H | Leukotriene A4 Hydrolase | 4,703763483 | 0,0000218 | 0,004906929167 |
| AF_17653 | HELZ | Helicase With Zinc Finger | 50 | 0,0000223 | 0,004906929167 |
| AF_1775 | PCCA | Propionyl-CoA Carboxylase Subunit Alpha | 50 | 0,0000259 | 0,005471116 |
| AF_17836 | CDC27 | Cell Division Cycle 27 | 4,318131157 | 0,000029 | 0,005890346154 |
| AF_1783 | DOCK9 | Dedicator Of Cytokinesis 9 | 4,53879479 | 0,0000321 | 0,006278522222 |
| AF_18346 | PNPT1 | Polyribonucleotide Nucleotidyltransferase 1 | 5,359372452 | 0,0000343 | 0,006301993333 |
| AF_18686 | RNF141 | Ring Finger Protein 141 | 50 | 0,0000357 | 0,006301993333 |
| AF_1887 | TMEM68 | Transmembrane Protein 68 | 50 | 0,0000358 | 0,006301993333 |
| AF_2282 | NME7 | NME/NM23 Family Member 7 | 8,933255746 | 0,0000415 | 0,006848796875 |
| AF_2359 | WDHD1 | WD Repeat And HMG-Box DNA Binding Protein 1 | 3,463530025 | 0,0000595 | 0,00952180303 |
| AF_2812 | COCH | Cochlin | 4,984052411 | 0,0000709 | 0,01101243824 |
| AF_2889 | AGL | Amylo-Alpha-1, 6-Glucosidase, 4-Alpha-Glucanotransferase | 50 | 0,0000734 | 0,01107501143 |
| AF_2954 | JMJD1C | Jumonji Domain Containing 1C | 50 | 0,0000781 | 0,01145683611 |
| AF_3445 | TMA16 | Translation Machinery Associated 16 Homolog | 5,608371484 | 0,0000863 | 0,01231757568 |
| AF_3520 | SUSD5 | Sushi Domain Containing 5 | 6,091471237 | 0,000092 | 0,01278557895 |
| AF_3784 | NEDD4 | NEDD4 E3 Ubiquitin Protein Ligase | 3,408788434 | 0,0001034193348 | 0,01400403864 |
| AF_3816 | SYCP1 | Synaptonemal Complex Protein 1 | 45,98022099 | 0,0001324912403 | 0,017492156 |
| AF_3995 | ITPR3 | Inositol 1,4,5-Trisphosphate Receptor Type 3 | 2,343454673 | 0,0001435518409 | 0,01810492095 |
| AF_4510 | NME8 | NME/NM23 Family Member 8 | 50 | 0,000143989146 | 0,01810492095 |
| AF_4536 | AVL9 | AVL9 Cell Migration Associated | 4,035448742 | 0,0001482325433 | 0,01820502468 |
| AF_4576 | HERC5 | HECT And RLD Domain Containing E3 Ubiquitin Protein Ligase 5 | 50 | 0,000162617641 | 0,01951781277 |
| AF_4632 | NPNT | Nephronectin | 5,143337172 | 0,0001908419878 | 0,0223963675 |
| AF_4804 | EVC2 | EvC Ciliary Complex Subunit 2 | 5,857175477 | 0,0002091719408 | 0,0227100099 |
| AF_5022 | HERC1 | HECT And RLD Domain Containing E3 Ubiquitin Protein Ligase | 12,18225009 | 0,0002092371906 | 0,0227100099 |
| AF_5029 | Hsp40 | DnaJ Heat Shock Protein Family | 5,696852219 | 0,0002202738908 | 0,0227100099 |
| AF_5042 | RBM19 | RNA Binding Motif Protein 19 | 5,275586623 | 0,0002208192394 | 0,0227100099 |
| AF_5623 | IDUA | Alpha-L-Iduronidase | 5,857869718 | 0,0002248507643 | 0,0227100099 |
| AF_5943 | SMARCAL1 | Actin Dependent Regulator Of Chromatin, Subfamily A Like 1 | 5,061129038 | 0,0002249640859 | 0,0227100099 |
| AF_6396 | MATN2 | Matrilin 2 | 50 | 0,0002257383151 | 0,0227100099 |
| AF_6608 | SLC39A12 | Solute Carrier Family 39 Member 12 | 2,990206511 | 0,0002316498662 | 0,0227100099 |
| AF_6996 | RPL31 | Ribosomal Protein L31 | 50 | 0,0002322174843 | 0,0227100099 |
| AF_7116 | HMGCL | 3-Hydroxy-3-Methylglutaryl-CoA Lyase | 50 | 0,0002587200563 | 0,02454949982 |
| AF_7276 | UNC13C | Unc-13 Homolog C | 4,541280096 | 0,0002632738049 | 0,02454949982 |
| AF_7370 | TMC3 | Transmembrane Channel Like 3 | 4,870346811 | 0,0002649728252 | 0,02454949982 |
| AF_7436 | ELP2 | Elongator Acetyltransferase Complex Subunit 2 | 50 | 0,0002777042813 | 0,02528545361 |
| AF_7503 | MYLK3 | Myosin Light Chain Kinase 3 | 7,40606769 | 0,000309483481 | 0,027396877 |
| AF_755 | PGM2L1 | Phosphoglucomutase 2 Like 1 | 50 | 0,0003112691953 | 0,027396877 |
| AF_7564 | SLC7A6 | Solute Carrier Family 7 Member 6 | 3,588501998 | 0,0003317227172 | 0,02871848639 |
| AF_7895 | GLS | Glutaminase | 4,076720222 | 0,0004077740001 | 0,03473313701 |
| AF_8156 | ASAH1 | N-Acylsphingosine Amidohydrolase 1 | 48,43880498 | 0,0004240274354 | 0,03554426803 |
| AF_8173 | SLC7A2 | Solute Carrier Family 7 Member 2 | 38,12050412 | 0,0004489436309 | 0,0370448643 |
| AF_8703 | TDRD1 | Tudor Domain Containing 1 | 5,16834605 | 0,0004968792884 | 0,04002047086 |
| AF_8782 | CPXM2 | Carboxypeptidase X, M14 Family Member 2 | 3,865603805 | 0,0005001611583 | 0,04002047086 |
| AF_8972 | INTS2 | Integrator Complex Subunit 2 | 17,57559971 | 0,0005828530974 | 0,0451634167 |
| AF_8983 | APPBP2 | Amyloid Beta Precursor Protein Binding Protein 2 | 2,937027461 | 0,0005979239328 | 0,0451634167 |
| AF_9101 | LRPPRC | Leucine Rich Pentatricopeptide Repeat Containing | 3,107315367 | 0,0006002563028 | 0,0451634167 |
| AF_9269 | PPP1R9A | Protein Phosphatase 1 Regulatory Subunit 9A | 9,516051849 | 0,0006039642174 | 0,0451634167 |
| AF_9464 | NRG3 | Neuregulin 3 | 6,285501843 | 0,0006071960965 | 0,0451634167 |
| AF_980 | DHX36 | DEAH-Box Helicase 36 | 4,556533591 | 0,0000412 | 0,006848796875 |

**Table S6. Candidate genes for which RELAX found signals of intensified selection (K > 1) in *A. patagonicus* (FDR < 0.05)**

| Gene ID | Gene name | Gene description | K | P-value | Adjusted p-value |
|---|---|---|---|---|---|
| AF_15753 | A2ML1 | Alpha-2-Macroglobulin Like 1 | 28,96233276 | 1,71E-13 | 0,000000000891936 |
| AF_11047 | GOLGA1 | Golgin A1 | 3,844974319 | 2,99E-12 | 0,00000000779792 |
| AF_8752 | SPTLC3 | Serine Palmitoyltransferase Long Chain Base Subunit 3 | 14,30144609 | 0,000000164 | 0,00026732 |
| AF_8402 | CCT8 | Chaperonin Containing TCP1 Subunit 8 | 3,84427245 | 0,000000205 | 0,00026732 |
| AF_2550 | PLEKHH1 | Pleckstrin Homology, MyTH4 And FERM Domain Containing H1 | 11,09394818 | 0,000000361 | 0,0003765952 |
| AF_9727 | SEPT2 | Septin 2 | 47,28415224 | 0,00000124 | 0,001077973333 |
| AF_17888 | UROC1 | Urocanate Hydratase 1 | 10,52857883 | 0,00000308 | 0,00204076 |
| AF_4689 | AIM1 | Crystallin Beta-Gamma Domain Containing 1 | 50 | 0,00000313 | 0,00204076 |
| AF_2516 | CPB2 | Carboxypeptidase B2 | 5,115426218 | 0,00000383 | 0,002219697778 |
| AF_4955 | CLDN18 | Claudin 18 | 50 | 0,00000702 | 0,003661632 |
| AF_7552 | LETM1 | Leucine Zipper And EF-Hand Containing Transmembrane Protein 1 | 38,8346529 | 0,00000788 | 0,003736552727 |
| AF_5241 | MEP1B | Meprin A Subunit Beta | 29,50697063 | 0,0000124 | 0,005389866667 |
| AF_5889 | RARS | Arginyl-TRNA Synthetase 1 | 3,038511142 | 0,0000135 | 0,005416615385 |
| AF_8924 | RBBP6 | RB Binding Protein 6, Ubiquitin Ligase | 38,87934461 | 0,000017 | 0,006333714286 |
| AF_1498 | HEPH | Hephaestin | 3,810692817 | 0,0000211 | 0,007337173333 |
| AF_15558 | ECE1 | Endothelin Converting Enzyme 1 | 3,117268144 | 0,0000266 | 0,007997866667 |
| AF_2837 | TNRC6B | Trinucleotide Repeat Containing Adaptor 6B | 2,891330917 | 0,0000268 | 0,007997866667 |
| AF_12888 | MEI1 | Meiotic Double-Stranded Break Formation Protein 1 | 14,12997858 | 0,0000276 | 0,007997866667 |
| AF_3806 | BAZ1A | Bromodomain Adjacent To Zinc Finger Domain 1A | 10,40442415 | 0,0000382 | 0,01048690526 |
| AF_13823 | WDR35 | WD Repeat Domain 35 | 4,690558028 | 0,000043 | 0,0112144 |
| AF_9658 | PPP1R21 | Protein Phosphatase 1 Regulatory Subunit 21 | 50 | 0,0000609 | 0,0146048 |
| AF_14715 | ADGRL3 | Adhesion G Protein-Coupled Receptor L3 | 47,00028235 | 0,000063 | 0,0146048 |
| AF_2094 | OGFR | Opioid Growth Factor Receptor | 46,1 | 0,0000644 | 0,0146048 |
| AF_13259 | PTPN11 | Protein Tyrosine Phosphatase Non-Receptor Type 11 | 10,07599425 | 0,0000708 | 0,0153872 |
| AF_285 | ANKZF1 | Ankyrin Repeat And Zinc Finger Peptidyl TRNA Hydrolase 1 | 8,37681621 | 0,0000919 | 0,019174016 |
| AF_15160 | DDX1 | DEAD-Box Helicase 1 | 4,647765809 | 0,0001058074996 | 0,02114278623 |
| AF_8479 | PDC | Phosducin | 4,806996748 | 0,0001162518623 | 0,02114278623 |
| AF_6454 | DPEP2 | Dipeptidase 2 | 18,77623626 | 0,0001207897653 | 0,02114278623 |
| AF_15827 | CFAP97 | Cilia And Flagella Associated Protein 97 | 12,97242384 | 0,0001224644622 | 0,02114278623 |
| AF_15093 | SYNPO2L | Synaptopodin 2 Like | 50 | 0,0001225926527 | 0,02114278623 |
| AF_8128 | ZNF488 | Zinc Finger Protein 488 | 29,48334142 | 0,0001256568967 | 0,02114278623 |
| AF_14480 | WDR11 | WD Repeat Domain 11 | 47,32465792 | 0,000184692065 | 0,0301048066 |
| AF_16647 | TRPV3 | Transient Receptor Potential Cation Channel Subfamily V Member 3 | 50 | 0,0002285431435 | 0,03612366777 |
| AF_11086 | SLCO1C1 | Solute Carrier Organic Anion Transporter Family Member 1C1 | 50 | 0,000247227765 | 0,03792764771 |
| AF_6686 | PARVG | Parvin Gamma | 49,74626866 | 0,0003276404311 | 0,04882778539 |

**Table S7. Candidate genes with positively selected sites in *A.forsteri* and *A.patagonicus*, predicted by CODEML branch-site models (FDR < 0.05)**

| Lineage | Gene ID | Gene name | Gene description | Likelihood Null Model | Likelihood Alternative Model | LRT | d.f | Adjusted p-value | BEB Analysis |
|---|---|---|---|---|---|---|---|---|---|
| *A. forsteri* | AF_4494 | LNX2 | Ligand of Numb protein X 2 | -6578,64252 | -6575,726939 | 5,831162 | 1 | 0,0324436654545455 | 12 W 0.951* |
| | | | | | | | | | 18 T 0.827 |
| | | | | | | | | | 26 D 0.819 |
| *A. forsteri* | AF_4640 | SGMS2 | Sphingomyelin synthase 2 | -2943,85942 | -2933,884229 | 19,950382 | 1 | 3,86036485714286e-05 | 281 L 0.999** |
| | | | | | | | | | 282 L 1.000** |
| | | | | | | | | | 283 T 0.977* |
| | | | | | | | | | 284 I 0.976* |
| *A. forsteri* | AF_5071 | TFAP2C | Transcription factor AP-2 gamma | -1987,396934 | -1974,065937 | 26,661994 | 1 | 2,35416e-06 | 158 C 0.996** |
| | | | | | | | | | 159 F 0.998** |
| | | | | | | | | | 160 R 0.873 |
| *A. forsteri* | AF_5373 | UFD1L | Ubiquitin recognition factor in ER associated degradation 1 | -2194,528675 | -2188,770013 | 11,517324 | 1 | 0,00195360033333333 | 1 M 0.998** |
| | | | | | | | | | 2 T 0.936 |
| | | | | | | | | | 3 V 0.933 |
| *A. forsteri* | AF_7017 | GMEB1 | Glucocorticoid modulatory element binding protein 1 | -2937,594909 | -2931,005189 | 13,17944 | 1 | 0,000916500761904762 | 1 M 0.997** |
| | | | | | | | | | 2 N 0.817 |
| *A. forsteri* | AF_6831 | WNT3A | Wnt family member 3A | -2311,378253 | -2300,910396 | 20,935714 | 1 | 2,75173333333333e-05 | 71 R 1.000** |
| | | | | | | | | | 72 A 0.884 |
| | | | | | | | | | 73 V 0.878 |
| | | | | | | | | | 77 P 0.948 |
| | | | | | | | | | 78 P 0.830 |
| *A. forsteri* | AF_7895 | GLS | Glutaminase | -3109,236662 | -3094,430938 | 29,611448 | 1 | 7,17944e-07 | 23 R 0.965* |
| | | | | | | | | | 218 A 1.000** |
| | | | | | | | | | 220 R 1.000** |
| | | | | | | | | | 221 H 0.963* |
| *A. forsteri* | AF_8749 | MCMBP | Minichromosome maintenance complex binding protein | -5446,9791 | -5439,253415 | 15,45137 | 1 | 0,00028782904 | 49 V 0.908 |
| | | | | | | | | | 50 K 0.853 |
| | | | | | | | | | 51 E 0.655 |
| | | | | | | | | | 146 G 0.942 |
| | | | | | | | | | 147 C 0.941 |
| | | | | | | | | | 331 Q 0.938 |
| *A. forsteri* | AF_8543 | IDE | Insulin degrading enzyme | -6780,555004 | -6769,523983 | 22,062042 | 1 | 1,99450044444444e-05 | 384 Q 0.749 |
| | | | | | | | | | 548 H 0.547 |
| | | | | | | | | | 557 R 0.774 |
| | | | | | | | | | 566 L 0.529 |
| *A. forsteri* | AF_8972 | INTS2 | Integrator complex subunit 2 | -10187,45183 | -10167,08127 | 40,741114 | 1 | 9,52e-09 | 484 S 0.997** |
| | | | | | | | | | 485 V 0.974* |
| | | | | | | | | | 486 A 0.998** |
| *A. forsteri* | AF_9132 | HDHD2 | Haloacid dehalogenase like hydrolase domain containing 2 | -2559,627097 | -2545,489555 | 28,275084 | 1 | 1,19272e-06 | 21 L 0.791 |
| | | | | | | | | | 22 R 0.950 |
| | | | | | | | | | 23 S 0.950 |
| | | | | | | | | | 107 C 0.797 |
| *A. forsteri* | AF_9261 | DYNC1I2 | Dynein cytoplasmic 1 intermediate chain 2 | -4323,901549 | -4307,812753 | 32,177592 | 1 | 3,1892e-07 | 299 Q 0.999** |
| | | | | | | | | | 300 Q 0.998** |
| *A. forsteri* | AF_9961 | ANKRD17 | Ankyrin repeat domain 17 | -14885,71439 | -14879,29238 | 12,844022 | 1 | 0,00100095704347826 | 177 T 0.599 |
| | | | | | | | | | 1081 V 0.724 |
| | | | | | | | | | 1150 A 0.563 |
| *A. forsteri* | AF_11846 | PCMT1 | Protein-L-isoaspartate (D-aspartate) O-methyltransferase | -1167,054052 | -1158,865431 | 16,377242 | 1 | 0,0002144912 | 195 D 0.909 |
| *A. forsteri* | AF_13292 | SON | SON DNA binding protein | -4086,87545 | -4066,972694 | 39,805512 | 1 | 9,52e-09 | 6 S 0.694 |
| | | | | | | | | | 26 R 0.629 |
| *A. forsteri* | AF_13270 | TIAM1 | TIAM Rac1 associated GEF 1 | -13709,29822 | -13704,80283 | 8,990766 | 1 | 0,00683392948148148 | 963 A 0.821 |
| | | | | | | | | | 1319 E 0.538 |
| | | | | | | | | | 1485 V 0.728 |
| *A. forsteri* | AF_262 | CTNNB1 | Catenin beta 1 | -5888,046367 | -5881,508793 | 13,075148 | 1 | 0,000924923636363636 | 1 Q 0.643 |
| | | | | | | | | | 703 A 0.889 |
| | | | | | | | | | 704 V 0.885 |
| | | | | | | | | | 705 S 0.777 |
| *A. forsteri* | AF_475 | ZNF326 | Zinc finger protein 326 | -3074,599753 | -3070,989783 | 7,21994 | 1 | 0,0169057660689655 | 219 K 0.946 |
| | | | | | | | | | 220 K 0.998** |
| | | | | | | | | | 221 K 0.819 |
| | | | | | | | | | 223 K 0.946 |
| | | | | | | | | | 226 T 0.947 |
| *A. forsteri* | AF_13872 | TEC | Tec protein tyrosine kinase | -4800,546021 | -4796,991208 | 7,109626 | 1 | 0,0173787826666667 | 149 M 0.862 |
| | | | | | | | | | 172 K 0.995** |
| | | | | | | | | | 477 G 0.868 |
| *A. forsteri* | AF_14805 | CBFA2T2 | CBFA2/RUNX1 partner transcriptional co-repressor 2 | -4820,717669 | -4815,448563 | 10,538212 | 1 | 0,0031805368 | 359 E 0.675 |
| | | | | | | | | | 362 G 0.657 |
| | | | | | | | | | 363 G 0.704 |
| | | | | | | | | | 364 G 0.996** |
| | | | | | | | | | 368 A 0.967* |
| *A. forsteri* | AF_15280 | TRPM8 | Transient receptor potential cation channel | -9342,079787 | -9334,315259 | 15,529056 | 1 | 0,00028782904 | 1049 T 0.808 |
| | | | | | | | | | 1060 M 0.870 |
| *A. forsteri* | AF_16232 | PAK1 | p21 (RAC1) activated kinase 1 | -3875,695768 | -3865,027497 | 21,336542 | 1 | 2,6201488e-05 | 471 F 0.999** |
| | | | | | | | | | 473 K 0.998** |
| | | | | | | | | | 474 F 0.945 |
| *A. forsteri* | AF_16462 | TRRAP | Transformation/transcription domain associated protein | -1093,238656 | -1089,565938 | 7,345436 | 1 | 0,0163280677142857 | 41 Q 0.997** |
| *A. forsteri* | AF_16779 | TMEM136 | Transmembrane protein 136 | -2514,418204 | -2505,261884 | 18,31264 | 1 | 8,49814133333333e-05 | 61 F 0.999** |
| | | | | | | | | | 62 S 0.979* |
| | | | | | | | | | 65 Q 0.960* |
| | | | | | | | | | 66 V 0.998** |
| *A. forsteri* | AF_16783 | TRIM29 | Tripartite motif containing 29 | -4507,973276 | -4500,069787 | 15,806978 | 1 | 0,0002649858 | 487 D 0.934 |
| | | | | | | | | | 489 S 0.932 |
| | | | | | | | | | 490 L 0.999** |
| | | | | | | | | | 491 K 0.995** |
| | | | | | | | | | 492 G 0.933 |
| *A. forsteri* | AF_17377 | ZYG11B | Protein zyg-11 homolog B | -5910,2849 | -5907,044897 | 6,480006 | 1 | 0,0239304327741935 | 1 M 0.597 |
| | | | | | | | | | 53 V 0.795 |
| *A. forsteri* | AF_17652 | PSMD12 | Proteasome 26S subunit, non-ATPase 12 | -2445,634772 | -2437,477009 | 16,315526 | 1 | 0,0002144912 | 113 T 0.997** |
| | | | | | | | | | 114 R 0.995** |
| | | | | | | | | | 115 H 0.856 |
| *A. forsteri* | AF_17836 | CDC27 | Cell division cycle 27 | -5856,389289 | -5846,05131 | 20,675958 | 1 | 2,84526123076923e-05 | 179 G 0.835 |
| | | | | | | | | | 688 G 0.995** |
| | | | | | | | | | 690 T 0.840 |
| | | | | | | | | | 691 F 0.627 |
| *A. forsteri* | AF_867 | CLCN7 | Chloride voltage-gated channel 7 | -7091,645003 | -7075,991059 | 31,307888 | 1 | 3,7434e-07 | 683 P 0.862 |
| | | | | | | | | | 687 G 0.856 |
| | | | | | | | | | 689 C 0.999** |
| | | | | | | | | | 692 R 0.966* |
| | | | | | | | | | 693 Q 0.832 |
| *A. forsteri* | AF_980 | DHX36 | DEAH-box helicase 36 | -8350,885653 | -8346,101466 | 9,568374 | 1 | 0,00517736307692308 | 40 G 0.952* |
| | | | | | | | | | 261 S 1.000** |
| | | | | | | | | | 340 V 0.950* |
| | | | | | | | | | 798 T 0.953* |
| *A. forsteri* | AF_1408 | NUP205 | Nucleoporin 205 | -6746,865304 | -6735,552885 | 22,624838 | 1 | 1,673837e-05 | 448 S 0.984* |
| *A. forsteri* | AF_2954 | JMJD1C | Jumonji domain containing 1C | -13118,79475 | -13108,34812 | 20,893262 | 1 | 2,75173333333333e-05 | 255 Q 0.970* |
| *A. patagonicus* | AF_2954 | JMJD1C | Jumonji domain containing 1C | -13116,26543 | -13110,24293 | 12,044998 | 1 | 0,007062689848 | 818 P 0.908 |
| | | | | | | | | | 1543 K 0.914 |
| | | | | | | | | | 1544 S 0.992** |
| *A. patagonicus* | AF_4689 | ADGRL3 | Adhesion G protein-coupled receptor L3 | -3095,661257 | -3077,545649 | 36,231216 | 1 | 5,95e-08 | 230 H 0.999** |
| | | | | | | | | | 231 S 0.809 |
| | | | | | | | | | 232 D 0.817 |

| | | | | | | | | | |
|---|---|---|---|---|---|---|---|---|---|
| | | | | | | | | | 236 D 1.000** |
| *A. patagonicus* | AF_13259 | CCT8 | Chaperonin Containing TCP1 Subunit 8 | -3424,544245 | -3410,031141 | 29,026208 | 1 | 1,61862666666667e-06 | 248 I 0.832 |
| | | | | | | | | | 302 T 0.998** |
| | | | | | | | | | 303 C 1.000** |
| | | | | | | | | | 304 P 0.840 |
| | | | | | | | | | 379 I 0.837 |
| *A. patagonicus* | AF_4494 | LNX2 | Ligand of Numb protein X 2 | -6581,073176 | -6572,401156 | 17,34404 | 1 | 0,00053018716 | 191 D 0.924 |
| | | | | | | | | | 192 L 0.904 |
| | | | | | | | | | 193 P 0.904 |
| | | | | | | | | | 194 Q 0.910 |
| | | | | | | | | | 260 V 0.945 |

**Table S8. Candidate genes for which RELAX found signals of relaxed selection (K < 1) in *A. forsteri* (FDR < 0.05)**

| Gene ID | Gene name | Gene description | Gene function | K | P-value | Adjusted p-value |
|---|---|---|---|---|---|---|
| AF_1808 | SHROOM3 | Shroom Family Member 3 | This gene encodes a PDZ-domain-containing protein that belongs to a family of Shroom-related proteins. This protein may be involved in regulating cell shape in certain tissues. A similar protein in mice is required for proper neurulation | 0,061487 | 0 | 0,00000040874 |
| AF_9961 | ANKRD17 | Ankyrin Repeat Domain 17 | It has been suggested that this protein plays a role in both DNA replication and in both anti-viral and anti-bacterial innate immune pathways. | 0,089538 | 0,0000038 | 0,001935467 |
| AF_16351 | FOLH1B | Putative N-acetylated-alpha-linked Acidic Dipeptidase | Metallopeptidase activity and dipeptidase activity | 0,366502 | 0,0000341 | 0,008288242 |
| AF_12856 | VRK2 | Vaccinia Related Kinase 2 | Serine/threonine kinase that regulates several signal transduction pathways. Isoform 1 modulates the stress response to hypoxia and cytokines. | 0,069616 | 0,0000382 | 0,008583765 |
| AF_12356 | RARS2 | Arginyl-tRNA Synthetase 2, mitochondrial | This nuclear gene encodes a protein that localizes to the mitochondria, where it catalyzes the transfer of L-arginine to its cognate tRNA, an important step in translation of mitochondrially-encoded proteins. | 0,129315 | 0,0000603 | 0,01212347 |
| AF_475 | ZNF326 | Zinc Finger Protein 326 | Core component of the DBIRD complex, a multiprotein complex that acts at the interface between core mRNP particles and RNA polymerase II (RNAPII) and integrates transcript elongation with the regulation of alternative splicing. May play a role in neuronal differentiation and is able to bind DNA. | 0,000031 | 0,0000729 | 0,01367746 |
| AF_10189 | FLVCR1 | Feline Leukemia Virus Subgroup C Cellular Receptor 1 | This gene encodes a member of the major facilitator superfamily of transporter proteins. The encoded protein is a heme transporter that may play a critical role in erythropoiesis by protecting developing erythroid cells from heme toxicity. | 0,335002 | 0,0000939 | 0,01594213 |
| AF_14165 | RSRC2 | Arginine/Serine-rich Coiled-coil 2 | Diseases associated with RSRC2 include Strabismus and Myopia | 0,243106 | 0,0001179351 | 0,01917073 |
| AF_18677 | IPO7 | Importin 7 | Functions in nuclear protein import, either by acting as autonomous nuclear transport receptor or as an adapter-like protein in association with the importin-beta subunit KPNB1 | 0 | 0,0002248237 | 0,02816098 |
| AF_9862 | PHLPP1 | PH Domain and Leucine Rich Repeat Protein Phosphatase 1 | The encoded protein promotes apoptosis by dephosphorylating and inactivating the serine/threonine kinase Akt, and functions as a tumor suppressor in multiple types of cancer.Involved in the hippocampus-dependent long-term memory formation (By similarity). Involved in circadian control by regulating the consolidation of circadian periodicity after resetting (By similarity). Involved in development and function of regulatory T-cells | 0 | 0,0002440689 | 0,02913572 |
| AF_964 | CCNL1 | Cyclin L1 | Involved in pre-mRNA splicing. Functions in association with cyclin-dependent kinases (CDKs) | 0,374898 | 0,0003203334 | 0,03435755 |
| AF_4559 | ASAP1 | ArfGAP with SH3 domain, Ankyrin repeat and PH domain 1 | May coordinate membrane trafficking with cell growth or actin cytoskeleton remodeling by binding to both SRC and PIP2. May function as a signal transduction protein involved in the differentiation of fibroblasts into adipocytes and possibly other cell types | 0,01036983409 | 0,0003301746439 | 0,05882572017 |
| AF_11868 | FBXO5 | F-box Protein 5 | Regulator of APC activity during mitotic and meiotic cell cycle | 0 | 0,0003327825771 | 0,05882572017 |
| AF_5373 | UFD1L | Ubiquitin Fusion Degradation 1 like | Essential component of the ubiquitin-dependent proteolytic pathway which degrades ubiquitin fusion proteins | 0,293617511 | 0,0003696194827 | 0,06067039795 |
| AF_8796 | EDRF1 | Erythroid Differentiation Regulatory factor 1 | Transcription factor involved in erythroid differentiation. Involved in transcriptional activation of the globin gene | 0,3761251066 | 0,0004624720448 | 0,0673835077 |
| AF_401 | PTCH1 | Patched 1 | This gene encodes a member of the patched family of proteins and a component of the hedgehog signaling pathway. Hedgehog signaling is important in embryonic development and tumorigenesis. | 0,04620246266 | 0,0004691628038 | 0,0673835077 |
| AF_16232 | PAK1 | p21 (RAC1) activated kinase 1 | Protein kinase involved in intracellular signaling pathways downstream of integrins and receptor-type kinases that plays an important role in cytoskeleton dynamics, in cell adhesion, migration, proliferation, apoptosis, mitosis, and in vesicle-mediated transport processes. Plays a role in the regulation of insulin secretion in response to elevated glucose levels. | 0,313 | 0,0005412858331 | 0,0731691085 |

**Table S9. Candidate genes for which RELAX found signals of relaxed selection (K < 1) in *A. patagonicus* (FDR < 0.05)**

| Gene ID | Gene name | Gene description | Gene function | K | P-value | Adjusted p-value |
|---|---|---|---|---|---|---|
| AF_10582 | ACADS | Acyl-CoA Dehydrogenase Short Chain | This gene encodes a tetrameric mitochondrial flavoprotein, which is a member of the acyl-CoA dehydrogenase family. This enzyme catalyzes the initial step of the mitochondrial fatty acid beta-oxidation pathway. | 0,270306 | 0,0000000331 | 0,000063 |
| AF_14861 | CPO | Carboxypeptidase O | This gene is a member of the metallocarboxypeptidase gene family. | 0 | 0,00000279 | 0,002174 |
| AF_6089 | ZNF438 | Zinc Finger Protein 438 | Isoform 1 acts as a transcriptional repressor. | 0 | 0,0000491 | 0,015632 |
| AF_8002 | TBC1D31 | TBC1 Domain Family Member 31 | Diseases associated with TBC1D31 include Branchiootorenal Syndrome 1; dominant disorder characterized by sensorineural, conductive, or mixed hearing loss, structural defects of the outer, middle, and inner ear, branchial fistulas or cysts, and renal abnormalities ranging from mild hypoplasia to complete absence | 0,045369 | 0,0000000261 | 0,000063 |

**Table S10. GO biological process terms in candidate genes for positive selection suggested by both CODEML and aBSREL**

| GO category | Term ID | Adjusted p-value | Genes Number | Genes names |
|---|---|---|---|---|
| Wnt signaling pathway involved in heart development | GO:0003306 | 0,02090722984 | 2 | WNT3A,CTNNB1 |
| biosynthetic process | GO:0009058 | 0,02090722984 | 16 | WNT3A,ZDHHC8,CTNNB1,NEDD4,ANKRD17,RPL31,ZNF326,SMARCB1,TRIM29,TFAP2C,MCMBP,ADCY5,GMEB1,SGMS2,CBFA2T2,SIN3B |
| positive regulation of muscle tissue development | GO:1901863 | 0,02753468434 | 2 | WNT3A,CTNNB1 |
| cellular nitrogen compound biosynthetic process | GO:0044271 | 0,03517376825 | 13 | WNT3A,CTNNB1,NEDD4,RPL31,ZNF326,SMARCB1,TRIM29,TFAP2C,ADCY5,GMEB1,SGMS2,CBFA2T2,SIN3B |
| cellular localization | GO:0051641 | 0,03517376825 | 9 | WNT3A,XPO4,CTNNB1,NEDD4,NUP205,TRIM29,ADCY5,APPBP2,AP2A2 |
| dopaminergic neuron differentiation | GO:0071542 | 0,03517376825 | 2 | WNT3A,CTNNB1 |
| regulation of nucleic acid-templated transcription | GO:1903506 | 0,03517376825 | 10 | WNT3A,CTNNB1,NEDD4,ZNF326,SMARCB1,TRIM29,TFAP2C,GMEB1,CBFA2T2,SIN3B |
| RNA metabolic process | GO:0016070 | 0,03517376825 | 12 | WNT3A,CRNKL1,CTNNB1,NEDD4,ZNF326,SMARCB1,TRIM29,TFAP2C,GMEB1,CBFA2T2,SIN3B,INTS2 |
| response to lipid | GO:0033993 | 0,03517376825 | 4 | CTNNB1,NEDD4,ADCY5,SLC2A1 |
| developmental induction | GO:0031128 | 0,03517376825 | 2 | WNT3A,CTNNB1 |
| sister chromatid cohesion | GO:0007062 | 0,03582990517 | 2 | CTNNB1,MCMBP |
| gene expression | GO:0010467 | 0,04040696313 | 13 | WNT3A,CRNKL1,CTNNB1,NEDD4,RPL31,ZNF326,SMARCB1,TRIM29,TFAP2C,GMEB1,CBFA2T2,SIN3B,INTS2 |
| ceramide phosphoethanolamine metabolic process | GO:1905371 | 0,04481644826 | 1 | SGMS2 |
| organic cyclic compound metabolic process | GO:1901360 | 0,04481644826 | 13 | WNT3A,CRNKL1,CTNNB1,NEDD4,ZNF326,SMARCB1,TRIM29,TFAP2C,ADCY5,GMEB1,CBFA2T2,SIN3B,INTS2 |
| nuclear transport | GO:0051169 | 0,04481644826 | 3 | XPO4,NEDD4,NUP205 |
| cellular response to glucose stimulus | GO:0071333 | 0,04721095806 | 2 | SMARCB1,ADCY5 |

**Table S11. GO biological process terms in candidate genes for positive selection suggested by either CODEML or aBSREL**

| GO category | Term ID | Adjusted p-value | Genes Number | Genes names |
|---|---|---|---|---|
| response to stress | GO:0006950 | 0,00004058722031 | 61 | INTS7,UBA5,PLD4,TBK1,UBASH3B,MAP3K5,ENPP3,CLEC16A,LGALS8,OPA1,FANCL,PPARA,P2RX4,CHAF1A,EDEM2,SUMO1,ATG5,RASGRP1,UIMC1,AQP11,EIF2AK1,MDFIC,ZYX,STXBP4,JCHAIN,TH,PNPT1,CELSR1,PLEKHM2,DACT1,PSMC6,CTNNB1,COCH,MBIP,NEDD4,MAGI3,DAPK1,FANCA,NME8,AOAH,C7,NEK4,IFIH1,SMARCAL1,CUL4B,UFL1,ATRIP,EDEM3,MAP3K3,DLEC1,WNT3A,HRAS,MYLK3,CAPN3,RAD51C,ANKRD17,PHLPP1,TEC,PAK1,DHX36,IPO7 |
| response to external stimulus | GO:0009605 | 0,00005231652761 | 45 | STRBP,ANKRD27,TBK1,UBASH3B,MAP3K5,ENPP3,CLEC16A,LGALS8,CRB1,GLRB,PPARA,P2RX4,CEP192,RHOA,ATG5,PDZD2,RASGRP1,RAC1,EIF2AK1,KYAT1,ZYX,STXBP4,JCHAIN,USP53,PLEKHM2,BOC,CBX7,GAB1,COCH,DAPK1,FANCA,AOAH,NPNT,C7,IFIH1,WNT3A,HRAS,CAPN3,NRP1,NRG3,ANKRD17,SLIT2,PHLPP1,DHX36,IPO7 |
| immune system process | GO:0002376 | 0,0002485224227 | 43 | FLVCR1,UBA5,MYO1E,PLD4,TBK1,UBASH3B,ENPP3,LGALS8,GPNMB,ATP11C,SIX4,TFRC,TRAT1,SPPL2B,RHOA,ATG5,RASGRP1,RAC1,EIF2AK1,ZYX,STXBP4,JCHAIN,PLEKHM2,CTNNB1,COCH,DAPK1,FANCA,IRAK1BP1,C7,IFIH1,UFL1,PLCG1,WNT3A,HRAS,LEPR,CACNA1C,ANKRD17,SLIT2,PHLPP1,TEC,CREB1,DHX36,IPO7 |
| regulation of catabolic process | GO:0009894 | 0,0007657388642 | 24 | TRAF5,TBK1,CLEC16A,KIF25,PPARA,WDR91,SUMO1,AQP11,PNPT1,VPS13D,DACT1,PSMC6,SGSM3,NEDD4,DAPK1,EGF,HERC1,UFL1,NBAS,ASB9,SLC25A4,LRPPRC,LEPR,DHX36 |
| reproduction | GO:0000003 | 0,006569724938 | 25 | MYCBPAP,STRBP,SIX4,IQCG,FANCL,GLRB,INPP5B,VPS13A,FNDC3A,WEE2,TH,ADAD1,CTNNB1,BRDT,FANCA,NME8,HFM1,TFAP2C,SELENOP,SMC3,RAD51C,LEPR,TAF4,SLIT2,DHX36 |
| gical process involved in interspecies interaction between organ | GO:0044419 | 0,007656893518 | 22 | SMARCB1,TBK1,LGALS8,CEP192,RHOA,ATG5,RASGRP1,KYAT1,ZYX,STXBP4,JCHAIN,PLEKHM2,COCH,NEDD4,DAPK1,C7,IFIH1,HRAS,NRP1,ANKRD17,DHX36,IPO7 |
| circulatory system development | GO:0072359 | 0,01032228054 | 24 | PROX1,FLVCR1,MYO1E,CRB2,LGALS8,RHOA,AGO3,ATG5,TH,CTNNB1,GAB1,EGF,PLCG1,ROCK2,MAP3K3,WNT3A,MYBPC3,NRP1,LEPR,CACNA1C,ANKRD17,SLIT2,CREB1,DHX36 |
| hepatocyte growth factor receptor signaling pathway | GO:0048012 | 0,01149124281 | 3 | RAC1,NRP1,PAK1 |
| defense response | GO:0006952 | 0,01465043969 | 22 | PLD4,TBK1,ENPP3,LGALS8,PPARA,RASGRP1,EIF2AK1,ZYX,STXBP4,JCHAIN,PLEKHM2,COCH,DAPK1,FANCA,AOAH,C7,IFIH1,DLEC1,HRAS,MYLK3,ANKRD17,IPO7 |
| organophosphate ester transport | GO:0015748 | 0,01656475837 | 7 | ATP11C,ABCC5,SLC35B2,SLC17A9,OSBPL2,ATP11A,SLC25A4 |
| mitotic cell cycle process | GO:1903047 | 0,01727762692 | 16 | DYNC1H1,KIF25,GPNMB,NAE1,KNTC1,CEP192,RHOA,TOM1L2,STIL,KIF20A,DACT1,EGF,SMC3,RAD51C,NCAPG,ANKRD17 |
| interferon-alpha production | GO:0032607 | 0,01727762692 | 3 | TBK1,IFIH1,DHX36 |
| ubiquinone metabolic process | GO:0006743 | 0,02100251913 | 3 | COQ6,COQ9,COQ3 |
| chemical synaptic transmission | GO:0007268 | 0,02127127929 | 14 | CYFIP1,GLRB,P2RX4,TH,CTNNB1,ITPR3,PLCG1,WNT3A,HRAS,UNC13C,PTCHD1,ITPKA,PPP1R9A,NRG3 |
| tissue remodeling | GO:0048771 | 0,02173939945 | 7 | UBASH3B,GPNMB,TFRC,CRB1,ATG5,CTNNB1,LEPR |
| locomotion | GO:0040011 | 0,02252432364 | 31 | PROX1,CRB2,LGALS8,GPNMB,SIX4,IQCG,P2RX4,ARF4,RHOA,INPP5B,PDZD2,VPS13A,WWC1,RAC1,CELSR1,BOC,CBX7,CTNNB1,GAB1,NME8,EGF,ACAP3,PLCG1,ROCK2,MAP3K3,WNT3A,HRAS,NRP1,NRG3,SLIT2,PAK1 |
| muscle structure development | GO:0061061 | 0,0267082668 | 14 | PROX1,SIX4,RHOA,ATG5,LMOD2,BOC,CTNNB1,NPNT,WNT3A,ATP11A,MYBPC3,CAPN3,ANKRD17,CREB1 |
| cellular calcium ion homeostasis | GO:0006874 | 0,0281040751 | 10 | UBASH3B,P2RX4,CHERP,ATG5,PACS2,ITPR3,ADCY5,PLCG1,CAPN3,CACNA1C |
| behavior | GO:0007610 | 0,03220405041 | 14 | STRBP,GLRB,GPR176,ARF4,VPS13A,TH,CELSR1,ITPR3,ZDHHC8,SELENOP,ADCY5,PTCHD1,LEPR,CREB1 |
| regulation of respiratory burst | GO:0060263 | 0,03840462882 | 2 | RAC1,JCHAIN |
| heart contraction | GO:0060047 | 0,0396681173 | 7 | P2RX4,CACNA1D,SUMO1,ATG5,TH,ATP1A1,CACNA1C |
| eye photoreceptor cell development | GO:0042462 | 0,04234538582 | 3 | CRB2,CRB1,TH |
| cellular divalent inorganic cation homeostasis | GO:0072503 | 0,04391777767 | 10 | UBASH3B,P2RX4,CHERP,ATG5,PACS2,ITPR3,ADCY5,PLCG1,CAPN3,CACNA1C |
| fatty acid metabolic process | GO:0006631 | 0,04495075639 | 9 | ACOT12,ACSBG2,PPARA,EPHX1,ACOX3,AOAH,ACOT7,ASAH1,PTGR2 |
| cellular pigmentation | GO:0033059 | 0,04803600723 | 3 | ANKRD27,MREG,KIF13A |
| glycerol metabolic process | GO:0006071 | 0,04988690227 | 2 | GK5,COQ3 |
| morphogenesis of a branching epithelium | GO:0061138 | 0,03716719742 | 8 | PROX1,SIX4,CELSR1,CTNNB1,NPNT,EGF,SLIT2,PAK1 |
| cellular localization | GO:0051641 | 0,000003537184997 | 59 | PKDCC,DYNC1H1,ANKRD27,UBASH3B,SERAC1,KIF25,SIX4,TFRC,CRB1,GLRB,WDR91,NUP205,P2RX4,CHERP,CEP192,ARF4,SUMO1,TRAK2,ATG5,MREG,PACS2,RASGRP1,RAC1,AQP11,TOM1L2,TRIM29,STIL,STXBP4,LIMD2,IPO5,CELSR1,LMAN1,PLEKHM2,CTNNB1,STARD3NL,NEDD4,ITPR3,COPB2,EGF,KIF13A,OSBPL2,ADCY5,PLCG1,ROCK2,AKAP11,STX6,WNT3A,HRAS,AP2A2,SRSF10,UNC13C,EXPH5,PEX16,CAPN3,SEC24C,APPBP2,TLK1,XPO4,IPO7 |
| regulation of signaling | GO:0023051 | 0,00001120879736 | 63 | PROX1,TRAF5,NCLN,SMARCB1,RALGPS1,UBA5,CRB2,TBK1,UBASH3B,CYFIP1,MAP3K5,CLEC16A,SIPA1L2,GPNMB,OPA1,TFRC,TRAT1,RGS9,PPARA,P2RX4,CHERP,RHOA,CBFA2T2,STK40,RASGRP1,WWC1,RASA2,MDFIC,STXBP4,DACT1,SGSM3,CTNNB1,MBIP,NEDD4,MAGI3,ITPR3,DAPK1,FANCA,NPNT,EGF,PRDM16,MAP4K3,GRB14,STK36,ADCY5,UFL1,PLCG1,MAP3K3,WNT3A,HRAS,UNC13C,ITPKA,CAPN3,SHOC2,PPP1R9A,NRP1,NRG3,ANKRD17,SLIT2,PHLPP1,CREB1,PAK1,DHX36 |
| cellular component biogenesis | GO:0044085 | 0,01053640548 | 48 | PROX1,FBLN5,DYNC1H1,ANKRD27,CYFIP1,SIX4,OPA1,TFRC,IQCG,NUP205,CHAF1A,CEP192,RHOA,SUMO1,AGO3,ATG5,PACS2,RAC1,AQP11,LMOD2,IQUB,ZYX,STIL,JCHAIN,WEE2,CRNKL1,PNPT1,TMEM41B,DACT1,CTNNB1,NME8,OSBPL2,MYO10,STK36,CUL4B,ROCK2,SLC39A12,WNT3A,HRAS,SRSF10,CDKL5,PEX16,CAPN3,SMC3,NRP1,ASAP1,SLIT2,PAK1 |

| | | | | |
|---|---|---|---|---|
| anatomical structure morphogenesis | GO:0009653 | 0,0165389987 | 44 | PROX1,FLVCR1,MYO1E,CRB2,PKDCC,ANKRD27,CYFIP1,LGALS8,SIX4,OPA1,TFRC,CRB1,PPARA,FOXN4,RHOA,STK40,AGO3,RAC1,LMOD2,TH,PNPT1,CELSR1,BOC,CTNNB1,GAB1,COCH,NPNT,EGF,HERC1,MYO10,PLCG1,ROCK2,MAP3K3,WNT3A,MYBPC3,CAPN3,NOLC1,NRP1,LEPR,NRG3,CACNA1C,SLIT2,CREB1,PAK1 |
| cell junction | GO:0030054 | 0,0006013854666 | 37 | MYO1E,CRB2,CYFIP1,CRB1,GLRB,PDZRN3,RGS9,P2RX4,ARF4,RHOA,RAC1,GRIK4,ZYX,ARR3,TH,USP53,SGSM3,CTNNB1,GAB1,MAGI3,TP53BP2,ATP1A1,PLCG1,WNT3A,HRAS,UNC13C,CDKL5,PTCHD1,SLC7A2,MCMBP,RAD51C,PPP1R9A,NRP1,NRG3,CACNA1C,ASAP1,PAK1 |
| autophagy | GO:0006914 | 0,001862677973 | 15 | UBA5,TBK1,CLEC16A,KIF25,LGALS8,ATG5,VPS13A,PACS2,TMEM41B,VPS13D,DAPK1,HERC1,UFL1,SLC25A4,LEPR |
| response to hormone | GO:0009725 | 0,02567556464 | 14 | UBA5,PPARA,RHOA,STXBP4,TH,NEDD4,ATP1A1,GRB14,UFL1,SLC2A1,LEPR,SLIT2,CREB1,PAK1 |
